# A brain atlas of synapse protein lifetime across the mouse lifespan

**DOI:** 10.1101/2021.12.16.472938

**Authors:** Edita Bulovaite, Zhen Qiu, Maximilian Kratschke, Adrianna Zgraj, David G. Fricker, Eleanor J. Tuck, Ragini Gokhale, Shekib A. Jami, Paula Merino-Serrais, Elodie Husi, Thomas J. O’Dell, Javier DeFelipe, Noboru H. Komiyama, Anthony Holtmaat, Erik Fransén, Seth G.N. Grant

## Abstract

Protein turnover is required for synapse maintenance and remodelling and may impact memory duration. We quantified the lifetime of postsynaptic protein PSD95 in individual excitatory synapses across the mouse brain and lifespan, generating the Protein Lifetime Synaptome Atlas. Excitatory synapses have a wide range of protein lifetimes that may extend from a few hours to several months, with distinct spatial distributions in dendrites, neuron types and brain regions. Short protein lifetime (SPL) synapses are enriched in developing animals and in regions controlling innate behaviors, whereas long protein lifetime (LPL) synapses accumulate during development, are enriched in the cortex and CA1 where memories are stored, and are preferentially preserved in old age. The protein lifetime synaptome architecture is disrupted in an autism model, with synapse protein lifetime increased throughout the brain. These findings add a further layer to synapse diversity in the brain and enrich prevailing concepts in behavior, development, ageing and brain repair.

---

Protein turnover is fundamental for cellular adaptive responses. It maintains the proteome (proteostasis) by replacing proteins that have been modified or damaged and enables proteome remodeling in response to transcriptional changes (Kaushik and Cuervo, 2015; Labbadia and Morimoto, 2015). Protein turnover in brain synapses controls adaptive processes including synaptic transmission, plasticity and learning (Colledge et al., 2003; Ehlers, 2003; Kato et al., 2005; Patrick et al., 2003), and mutations that disrupt protein turnover result in neurodevelopmental disorders including autism and intellectual disability (Labbadia and Morimoto, 2015; Louros and Osterweil, 2016). Reduced protein removal in old age is a key mechanism of senescence (Kaushik and Cuervo, 2015; Sabath et al., 2020; Santra et al., 2019) and contributes to synaptic pathology in Alzheimer’s, Parkinson’s and other neurodegenerative diseases (Helton et al., 2008; Kaushik and Cuervo, 2015; Tai et al., 2012; Vilchez et al., 2014).

In 1984, Francis Crick highlighted the importance of the lifetime of synaptic proteins for memory, reasoning that protein turnover would erase the protein modifications that occur with learning (Crick, 1984). Several decades later, the turnover of synapse proteins was measured in bulk preparations and found to have a half-life of days to weeks (Cohen et al., 2013; Dörrbaum et al., 2018; Ehlers, 2003; El-Husseini et al., 2002; Fornasiero et al., 2018; Heo et al., 2018; Price et al., 2010). However, because of the technical challenges of measuring protein lifetime in individual synapses in the intact brain, which so far has been investigated using overexpressed exogenous proteins in small populations of neurons (Gray et al., 2006; Steiner et al., 2008; Villa et al., 2016), it is not known whether there are differences in the lifetime of proteins between synapses, whether specialized subsets of synapses exist with long protein lifetimes, or whether synapse protein lifetime and turnover change during development and aging. That synapse protein turnover rates may differ across the dendritic tree of neurons is suggested by the observation that dendrites contain protein synthesis and degradation machinery (Biever et al., 2020; Bingol and Schuman, 2006; Ostroff et al., 2002; Steward and Levy, 1982).

To visualize and quantify protein lifetime in individual excitatory synapses across the mouse brain and lifespan, we engineered the genome so that mice express the HaloTag protein domain fused to endogenous PSD95 (PSD95-HaloTag mice) (Fig. 1A, S1). PSD95 is a highly abundant postsynaptic scaffold protein that interacts with and organizes more than 100 other postsynaptic proteins, including NMDA and AMPA subtypes of neurotransmitter receptors (Fernandez et al., 2017; Fernandez et al., 2009; Frank et al., 2016; Husi et al., 2000). PSD95 is essential for synaptic plasticity and learning (Carlisle et al., 2008; Fernandez et al., 2017; Fitzgerald et al., 2015; Frank et al., 2016; Migaud et al., 1998; Nithianantharajah et al., 2013). The HaloTag forms a covalent bond with a small-molecule synthetic ligand coupled to a fluorophore (England et al., 2015; Hoelzel and Zhang, 2020; Los et al., 2008). Injecting the HaloTag ligand into PSD95-HaloTag mice labels excitatory synapses with a fluorescent date-stamp. The lifetime of PSD95 in a synapse is then measured by visualizing the duration the labeled protein remains. We combined this labeling strategy with our synaptome mapping pipeline technology (SYNMAP), which is capable of quantifying the fluorescent signals from billions of individual synapses across the mouse brain and lifespan (Cizeron et al., 2020; Zhu et al., 2018). We report the creation of a single-synapse resolution atlas (Protein Lifetime Synaptome Atlas; https://brain-synaptome.org/Protein_Lifetime/) of PSD95 protein lifetime across the brain in young, mature and old mice and in a mouse model of neurodevelopmental disorders.

**Fig. 1.**
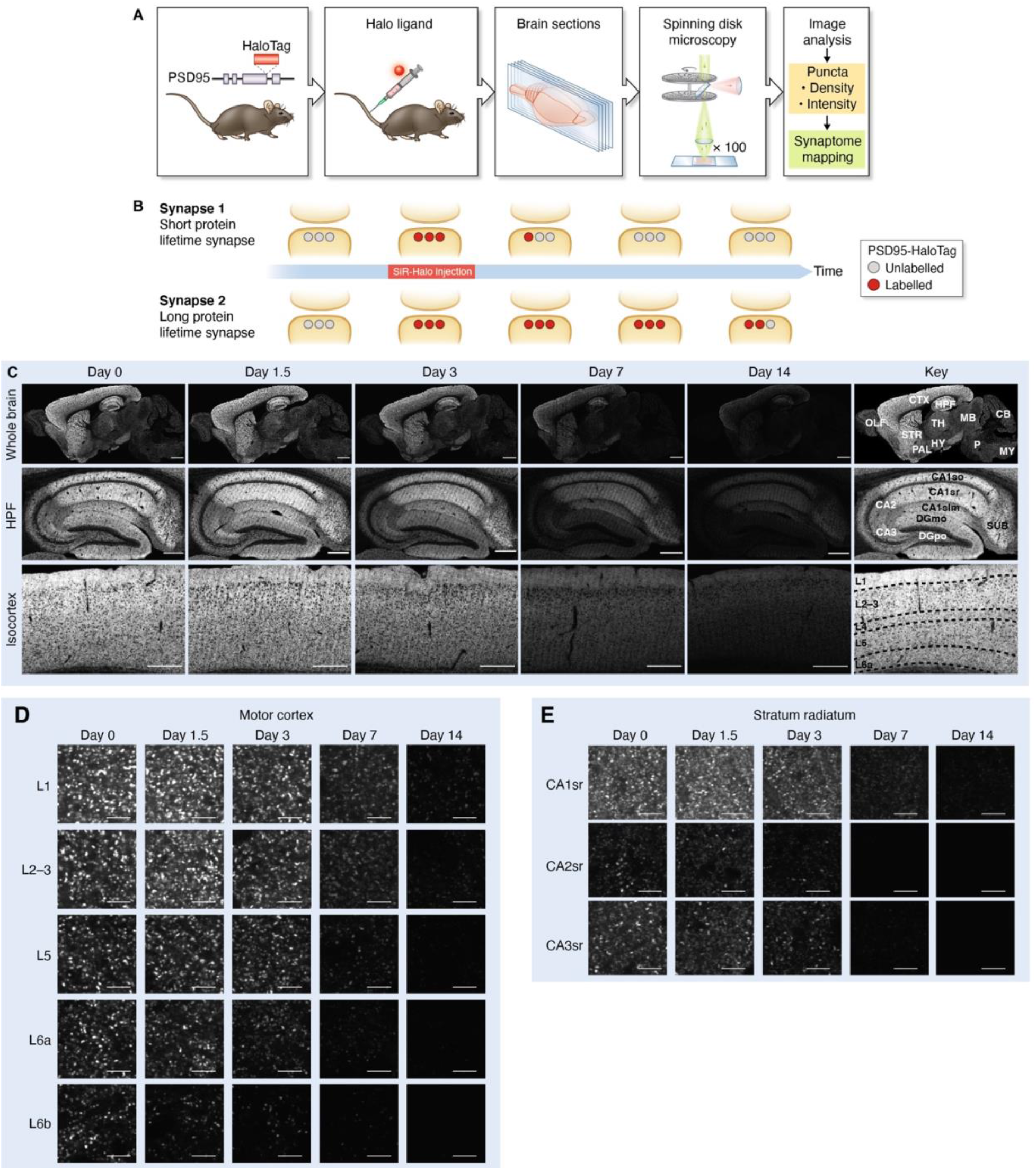
Visualizing synapse protein lifetime across the mouse brain. (**A**) Summary of the experimental flow. PSD95-HaloTag mice were injected with a Halo ligand, sagittal brain sections obtained and imaged using spinning disk confocal microscopy, and synaptic puncta analyzed and mapped. (**B**) Illustration of PSD95-HaloTag labeling in the postsynaptic terminal of synapses with short or long protein lifetime. (**C**) SiR-Halo fluorescence labeling at five time points post-injection, showing whole brain, hippocampal formation and layers of the isocortex. Key shows regions and layers. (**D**,**E**) High-magnification representative images of puncta fluorescence decay in layers of the motor cortex (D) and in stratum radiatum of CA1, CA2 and CA3 subregions of the HPF (E) over a 2-week period. CTX, isocortex; HPF, hippocampal formation; OLF, olfactory areas; STR, striatum; PAL, pallidum; HY, hypothalamus; TH, thalamus; MB, midbrain; CB, cerebellum; P, pons; MY, medulla; CA1so, CA1 stratum oriens; CA1sr, stratum radiatum; CA1slm, stratum lacunosum-moleculare; DGmo, dentate gyrus molecular layer; DGpo, dentate gyrus polymorphic layer; SUB, subiculum. Scale bars: 2 mm (C, whole brain), 500 µm (C, HPF and CTX), 5 µm (D,E).

## Genetic labeling and visualization of postsynaptic protein lifetime

In PSD95-HaloTag mice, PSD95 protein expression, protein complex assembly, and the physiology of synaptic transmission and synaptic plasticity are normal (Fig. S2, S3). To irreversibly label and visualize the PSD95-HaloTag *in vivo*, we injected a cell- and blood brain barrier-permeable fluorescent ligand (Silicon-Rhodamine-Halo, or SiR-Halo) (Lukinavicius et al., 2013; Masch et al., 2018) into the tail vein of PSD95-HaloTag mice and prepared sections of brain tissue for imaging on a spinning disk confocal microscope (pixel resolution 84 nm and optical resolution ∼260 nm) (Fig. 1A). Because SiR-Halo forms a covalent bond with the PSD95-HaloTag, we could examine the persistence of labeling after injection and identify the synapses and brain regions with the longest and shortest PSD95 protein lifetimes (Fig. 1B).

Imaging the brain of 3-month-old PSD95-HaloTag mice 6 hours (day 0) after injection revealed widespread labeling (Fig. 1C, S4), whereas none was detected in injected wild-type mice (Fig. S5). We found that 300 nmol of injected SiR-Halo was sufficient to saturate HaloTag binding sites (Fig. S6). We confirmed that the SiR-Halo ligand quantitatively labeled all brain regions by injecting it into compound heterozygous *Psd95*^HaloTag^;*Psd95*^eGFP^ (*Psd95*^HaloTag/eGFP^) mice, quantifying the SiR-Halo and eGFP synaptic puncta density in 110 brain regions, and then testing their correlation (R = 0.976, P < 0.0001) (Fig. S7). These results demonstrate that injection of a fluorescent ligand into PSD95-HaloTag mice efficiently and quantitatively labels PSD95 across all regions of the brain.

## Spatial diversity in synaptic protein lifetime

Images collected over the course of 2 weeks revealed a loss of labeling in all brain regions, with different regions showing different rates (Fig. 1C, S4). Synapses with the longest PSD95 protein lifetime were concentrated in the isocortex (neocortex) and hippocampal formation (HPF) (Fig. 1C, S4). Examination of images at single-synapse resolution revealed that each brain region is composed of populations of synapses with different PSD95 lifetimes (Fig. 1D, 1E, S8), some with long (weeks) and others with short (hours or days) protein lifetimes, which we hereafter refer to as LPL and SPL synapses, respectively. Moreover, these synapse populations were spatially organized. For example, within the laminar organization of the isocortex there was a clear gradient from superficial (layer 1) to deep (layer 6) layers, with synapses with the longest PSD95 lifetime located in layer 1 (Fig. 1C, 1D, S4, S8). In the HPF, the dendritic fields of pyramidal neurons in the CA1 exhibited synapses with longer PSD95 lifetimes than those in CA2 and CA3 (Fig. 1C, 1E, S4, S8). This pattern was evident in both basal (stratum oriens) and apical (stratum radiatum, stratum lacunosum, stratum moleculare) dendrites, indicating that synaptic protein lifetime is determined, at least in part, by cell-wide mechanisms that differ between pyramidal neuron subtypes (Fig. 1C, S8). Synapses in the dendritic fields of granule cell neurons in the dentate gyrus and olfactory bulb had short PSD95 protein lifetimes, further suggesting cell-type-specific turnover mechanisms (Fig. 1C, S8). Comparison of the large CA3 thorny excrescence synapses and those in the polymorphic layer of the dentate gyrus with the smaller CA1 synapses showed that the largest synapses of the HPF (Broadhead et al., 2016; Cizeron et al., 2020; Harris and Weinberg, 2012; Zhu et al., 2018) do not have the longest protein lifetimes (Fig. S8).

## A brainwide synaptome atlas of protein lifetime in excitatory synapses

To quantify PSD95 protein lifetime in brain subregions we calculated its half-life using single-synapse resolution data and SYNMAP synaptome mapping technology (Zhu et al., 2018). PSD95-HaloTag mice were injected with SiR-Halo ligand and parasagittal brain sections prepared and imaged at post-injection time points (6 hours, 1.5, 3, 7, 14 days) (Fig. 1C, S4). The density and intensity of SiR-Halo-labeled synaptic puncta were quantified in 12 overarching brain areas (isocortex, olfactory areas, HPF, cortical subplate, striatum, pallidum, thalamus, hypothalamus, midbrain, pons, medulla, cerebellum) and 110 subregions registered to the Allen Reference Atlas (Table S1). By fitting an exponential decay function to the data for every subregion we calculated the average PSD95 puncta density (^PSD95 density^t_1/2_) and intensity (^PSD95 intensity^t_1/2_) half-lives and plotted the values in brain maps (Fig. 2A, 2B, S9, S10). ^PSD95 density^t_1/2_ is derived from the reduction in the number of SiR-Halo-positive puncta, whereas ^PSD95 intensity^t_1/2_ is derived from the reduction in SiR-Halo signal within labeled synapses. The two half-life measures were highly correlated (R = 0.9088 and P < 0.0001; Fig. S11). These datasets were used to create the Protein Lifetime Synaptome Atlas (Bulovaite et al., 2021a).

**Fig. 2.**
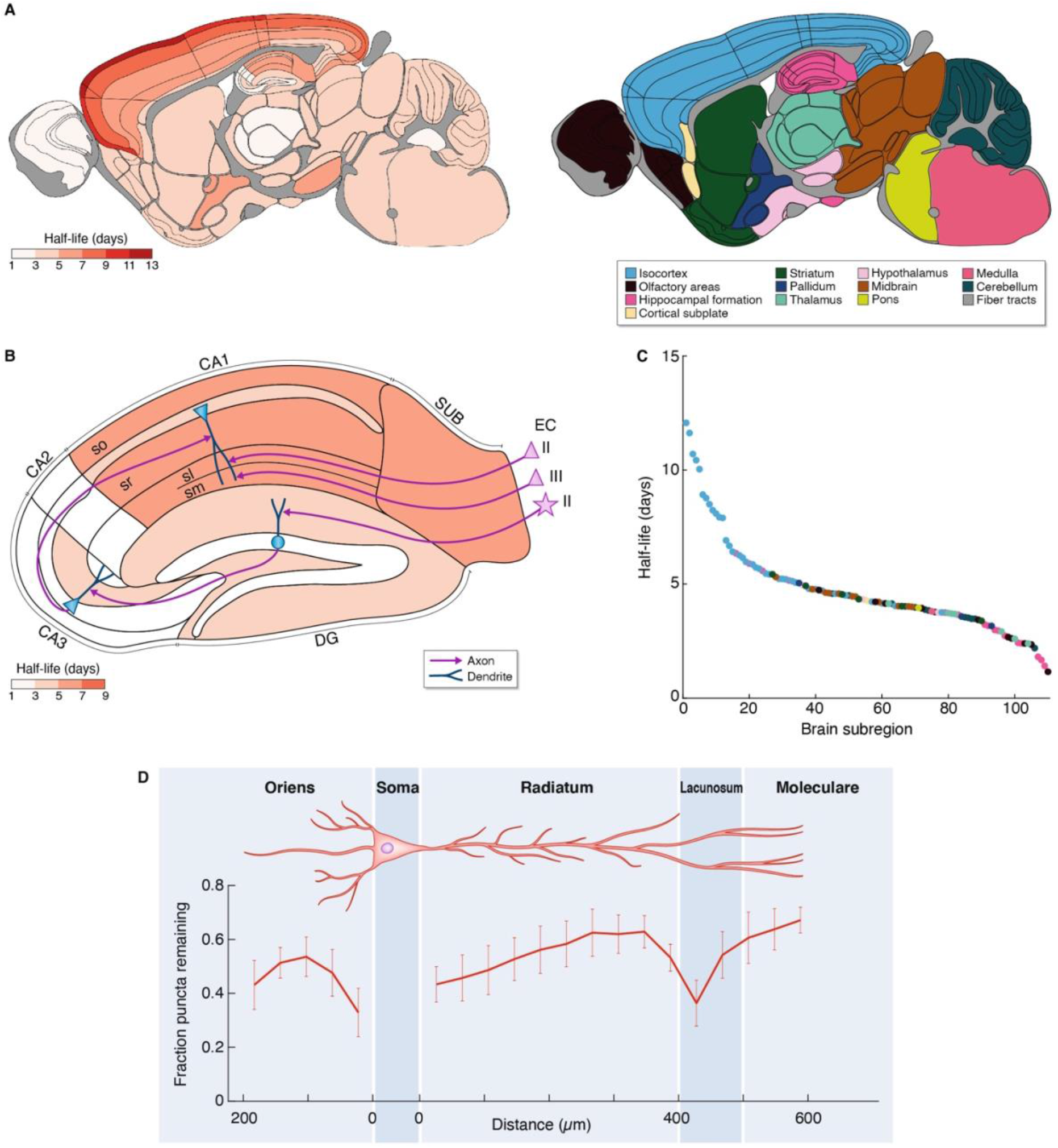
Brainwide atlas of synapse protein lifetimes. (**A**) Summary of PSD95 puncta half-life across 110 mouse brain subregions (Table S1). The twelve principal brain regions are color-coded to the right. (**B**) PSD95 puncta half-life in HPF, with key elements of the circuitry shown. The pyramidal neurons in entorhinal cortex layer II project to CA1sl, pyramidal neurons in entorhinal cortex layer III project to CA1sm, and stellate neurons in entorhinal cortex layer II project to dentate gyrus granule neurons, which in turn project to CA3, which project to CA1. CA, cornu ammonis; DG, dentate gyrus; EC, entorhinal cortex; sr, stratum radiatum; sl, stratum lacunosum; sm, stratum moleculare; so, stratum oriens. Adapted from (Marks et al., 2020). (**C**) The 110 brain subregions ranked according to their half-life, color-coded as in (A). (**D**) Fluorescent puncta decay (mean fraction of puncta remaining at day 7 compared with day 0, ± SD) across apical and basal dendrites of the CA1 pyramidal cells. Day 0, n=3; day 7, n=3 animals.

## Dendritic and subregional distribution of synapses with different PSD95 lifetimes

We observed a 10-fold difference in ^PSD95 density^t_1/2_ between the subregions with the longest (layer 1 of the motor cortex, 12.1 days) and shortest (olfactory bulb glomerular layer, 1.2 days) half-life (Fig. 2A, 2C, S9, S12). Comparison of ^PSD95 density^t_1/2_ for the 12 overarching brain areas showed that cortical structures are populated by synapses with the longest protein lifetimes, in contrast to subcortical structures which predominantly comprise synapses with the shortest protein lifetimes (Fig. 2A, 2C, S9, S10). All regions of the isocortex exhibited a gradient, with a 2.4-fold range in ^PSD95 density^t_1/2_ from layer 1 (10.6 days) to layer 6 (4.5 days) (Fig. 2A, S9). Similarly, the subregions within the HPF with the shortest ^PSD95 density^t_1/2_ were in initial pathways of the trisynaptic circuit (dentate gyrus, 3.5 days; CA2, 1.9 days; CA3, 3.0 days), whereas those with the longest ^PSD95 density^t_1/2_ were in the latter part of the circuit in the CA1 field (5.5 days) and subiculum (6.4 days) (Fig. 2B).

Although dendrites contain protein synthesis and degradation machinery (Biever et al., 2020; Bingol and Schuman, 2006; Ostroff et al., 2002; Steward and Levy, 1982) it is not known whether synapse protein lifetime varies across the dendritic tree. To determine whether synapse protein lifetime varies as a function of distance from the soma, we quantified ^PSD95 density^t_1/2_ in a series of parallel windows from the pyramidal neuron soma in the CA1 to the distal basal dendrites in stratum oriens and to the distal apical dendrites in stratum radiatum, stratum lacunosum and stratum moleculare (Fig. 2D). In apical dendrites, a gradient of increasing PSD95 protein lifetime was observed in stratum radiatum, which reversed at the boundary with stratum lacunosum (which receives inputs from entorhinal cortex layer II) before increasing again in distal stratum moleculare synapses (which receives inputs from entorhinal cortex layer III). Similarly, in the basal dendrites of stratum oriens there was an initial gradient in PSD95 protein lifetime before levelling and reversing in distal dendrites. Together, these data show that synapses with different protein lifetimes are differentially distributed throughout the dendritic tree and are dependent on the distance from the soma and the identity of the neuronal input.

## PSD95 turnover occurs in stable dendritic spines

Dendritic spines can be structurally stable for long or short durations (Grutzendler et al., 2002; Holtmaat et al., 2005; Trachtenberg et al., 2002; Zuo et al., 2005). We asked if protein turnover occurs within stable spines or whether the loss of HaloTag labeling might reflect spine elimination. To visualize dendritic spines housing PSD95 in individual apical dendrites, we dye filled CA1 pyramidal neurons from PSD95-HaloTag mice injected with SiR-Halo (Fig. 3A, S13). Consistent with previous results that show PSD95 is expressed in a subset of larger, more stable synapses (Cizeron et al., 2020; Fortin et al., 2014; Santuy et al., 2020; Zhu et al., 2018), a subset of spines expressed PSD95 at day 0 and labeled spines were still visible 10 days after SiR-Halo injection. If PSD95 were being turned over in synapses, we reasoned we could observe both ‘old’ and ‘new’ protein using a two-step labeling procedure. Three days after injecting mice with SiR-Halo we labeled brain sections with TMR-Halo and found that the two labels were colocalized in many synapses, especially in subregions with long PSD95 half-lives (Fig. 3B, S14).

**Fig. 3.**
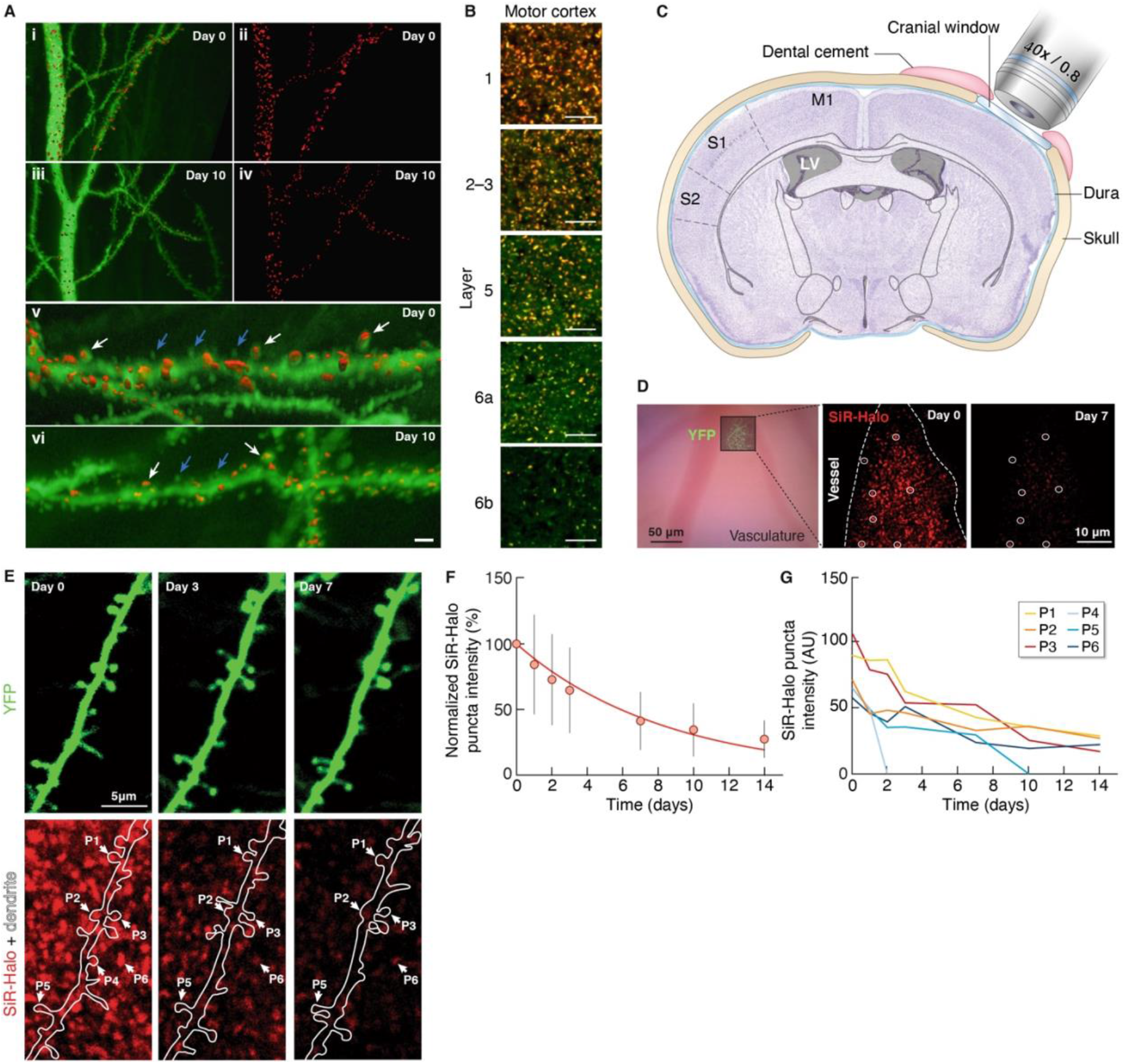
PSD95 turnover in dendritic spines. (**A**) 3D reconstructed dye-filled CA1 neurons and SiR-Halo-positive puncta. (i-iv) Confocal microscopy images of CA1 apical dendrites (stratum radiatum) intracellularly injected with Alexa 488 (green) at day 0 (i, ii) and at day 10 post-injection of SiR-Halo (red) (iii, iv). (v, vi) Higher-magnification of (i) and (iii), respectively, showing dendritic spines with SiR-Halo-positive puncta (white arrows) and without SiR-Halo signal (blue arrows). Scale bar: 5 µm (i-iv), 1.5 µm (v, vi). (**B**) Colocalization (orange) of SiR-Halo (red) injected at day 0 and post-fixation-applied TMR-Halo (green) at day 3 in synapses across layers of the motor cortex. Scale bars: 5 µm. (**C**) Schematic of the cranial window implant above the somatosensory cortex. (**D**) Bright-field image (left) of the superficial vasculature and, superposed, a fluorescence image of YFP-labeled axons and dendrites in cortical layer 1 that are used as fiducial points for tracking SiR-Halo ligand puncta over time, as shown for day 0 (center) and day 7 (right). (**E**) A YFP-labeled dendrite (top) containing SiR-Halo-labeled puncta (bottom), repeatedly imaged over days to weeks. SiR-Halo puncta can be tracked in stable and dynamic spines (e.g. P1-P5) and in the immediate surrounding of the dendrite (e.g. P6). (**F**) The mean (± SD) fluorescence intensities over 14 days of individual SiR-Halo puncta (n=295; N=3 mice) normalized to the first imaging time point. Red line, a single exponential fit of the means (λ=0.12/day; t_1/2_=5.8 days). (**G**) The measured fluorescence intensity (in arbitrary units) over time of the SiR-Halo puncta examples in (E).

To simultaneously visualize PSD95 turnover and spine turnover in the brain of living mice, we monitored individual synapses in layer 1 of somatosensory cortex through a cranial window (Holtmaat et al., 2009) in PSD95-HaloTag mice over weeks after injection of SiR-Halo (Fig. 3C). To visualize dendritic spines in individual neurons and provide fiducial markers for SiR-Halo-labeled synapses, we used adeno-associated viral vectors expressing YFP (Fig. 3D, 3E). Spine size was estimated based on YFP fluorescence intensities (Holtmaat et al., 2005). In individual synapses we observed decay profiles that were similar (Fig. 3F) to the population data (Fig. 2). A similar overall decay of fluorescence was observed when measured in regions that were imaged only once over the course of the experiment (Fig. S15), excluding photobleaching as a main factor in driving the fluorescence decay. In line with the dye injection experiment, not all spines expressed PSD95 to a similar extent (Fig. 3E, 3G), although in general SiR-Halo fluorescence intensities correlated well with spine size (Fig. S15) (Gray et al., 2006). Even within single dendrites, spines variably retained the SiR-Halo label. Label retention was weakly related to spine size (Fig. S15), and spines observed to disappear often had low levels of PSD95 or rapid loss of labeling (see examples in Fig. 3E, 3G), which is consistent with previous observations (Cane et al., 2014; Gray et al., 2006). Together, these data indicate that structurally stable spines differ in their PSD95 content and lifetime even within the same dendrite.

## Synapse classification by protein lifetime

The differences in protein lifetime between individual synapses raises important questions about the diversity of synapses that comprise the synaptome. In previous work we used a data-driven approach to classify synapses from across the whole brain and lifespan based on their ‘static’ protein composition (Cizeron et al., 2020; Zhu et al., 2018). Using two synaptic markers (PSD95eGFP and SAP102mKO2) we classified synapses into 3 types: type 1 express PSD95, type 2 express SAP102, and type 3 express both markers. Each of these types was further classified using morphological criteria into a total of 37 subtypes (Zhu et al., 2018). To determine whether the different types and/or subtypes of synapses have characteristic protein lifetimes, we combined synaptome mapping of PSD95-HaloTag with PSD95eGFP and SAP102mKO2 using triple knock-in mice (*Psd95*^HaloTag/eGFP^;*Sap102*^mKO2/y^). Six_-_month-old triple knock-in mice were injected with SiR-Halo and brain sections collected at day 0 and day 7. SiR-Halo-positive synapses were then categorized using PSD95eGFP and SAP102mKO2, allowing us to determine the PSD95 lifetime of two synapse types (type 1 and type 3) and 30 of the 37 known subtypes (all except subtypes 12-18, which do not express PSD95) in all brain areas. Ranking the synapse subtypes by their PSD95 protein lifetime from brainwide data (Fig. 4A) and individual regions (Fig. S15) showed that both type 1 and type 3 synapses contain subtypes with short and long protein lifetimes. Thus, populations of synapses with distinct molecular compositions were further divisible into subtypes with varying protein lifetimes. Synapse subtype 2 maintained the most (96.7%) and subtype 6 the least (23.3%) label after 7 days (Fig. 4A). To visualize the potential duration that any synapse subtype could retain copies of PSD95 we plotted exponential decay functions (Fig. 4B). This revealed that several synapse subtypes can retain copies of PSD95 for the natural lifespan of the mouse (Burt, 1940; Hamilton Jr, 1937).

**Fig. 4.**
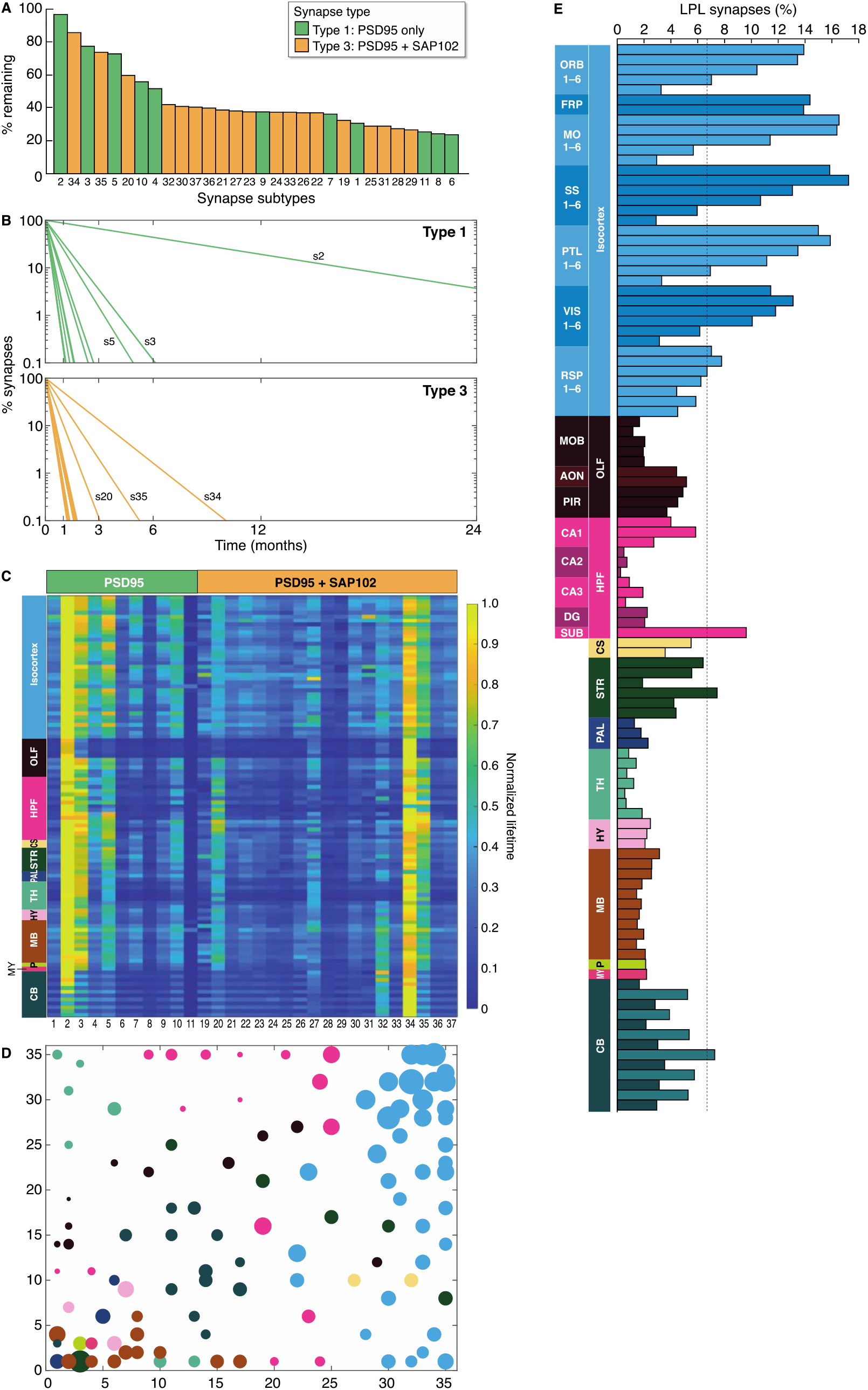
PSD95 protein lifetimes in synapse subtypes. (**A**) Ranked bar plot of synapse subtypes according to their associated PSD95 lifetimes using whole-brain data. Shown is the percentage of each SiR-Halo-positive subtype remaining at day 7 compared with day 0. Synapse subtypes (coded 1-37, of which thirty contain PSD95 and are encompassed by the present study) are classified as type 1 or type 3 according to whether they contain SAP102 in addition to PSD95. (**B**) Lifetime of PSD95 in type 1 and type 3 synapse subtypes, shown as percentage of synapses retaining PSD95. Longest-lived subtypes (s) are indicated. (**C**) Synapse subtype PSD95 protein lifetimes for 110 brain subregions represented as a heatmap. The values in each row are normalized. (**D**) Self-organizing feature map (SOM) clustering based on subtype densities. Each region is described by its 37 densities of the subtypes and a resulting two-dimensional SOM clustering is shown. Each subregion (see Fig. 2A for color key) is represented by a disc, the center position of which corresponds to the winning SOM node position and the radius codes for ^PSD95 intensity^t_1/2_. Axes indicate node index number in the 35×35 SOM node network. (**E**) Percentage of LPL synapses in brain regions and subregions. Regions: isocortex; OLF, olfactory areas; HPF, hippocampal formation; CS, cortical subplate; STR, striatum; PAL, pallidum; HY, hypothalamus; TH, thalamus; MB, midbrain; CB, cerebellum; P, pons; MY, medulla. Isocortex subregions: ORB, orbital area; FRP, frontal pole; MO, somatomotor area; SS, somatosensory area; PTL, posterior parietal association areas; VIS, visual areas; RSP, retrosplenial area. Olfactory area subregions: MOB, main olfactory bulb; AON, anterior olfactory nucleus; PIR, piriform area. HPF subregions: CA, cornu ammonis; DG, dentate gyrus; SUB, subiculum. The cortical layers in isocortex are numbered and shown from top to bottom.

## Building blocks of synaptome architecture

The protein lifetime of each subtype was independent of its location in the brain, with subtype lifetime rankings maintained across brain regions (Fig. 4C, S16). Crucially, this indicates that synapse subtypes with different protein lifetimes are general building blocks of whole-brain architecture. Furthermore, these findings suggest that the spatial distribution of synapse subtypes with different protein lifetimes underpins the half-life measurements from individual brain regions. To confirm this, we generated a self-organizing feature map (SOM), which reduces the high dimensionality of synaptome maps of the 37 subtypes to a two-dimensional representation (Fig. 4D, S17). The SOM shows a systematic organization in which anatomically related regions become closely clustered owing to their similarity in subtype density (Fig. S17). Consistent with our hypothesis, we observe a systematicity in half-lives when subregion half-life values are shown (Fig. 4D).

LPL synapses are particularly interesting because they could potentially maintain PSD95 modifications for long durations. When grouped together, the five subtypes with longest protein lifetimes (subtypes 2, 34, 3, 35, 5) represented 6.7% of PSD95-positive excitatory synapses in the whole brain; however, there were marked regional and subregional differences (Fig. 4E, S18), with a 15-fold range across the main brain regions: isocortex (11.9%), striatum (5.9%), cortical subplate (4.4%), cerebellum (3.9%), HPF (3.8%), olfactory areas (3.0%), midbrain (2.4%), hypothalamus (2.2%), medulla (2.2%), pons (2.1%), pallidum (1.8%), thalamus (0.8%). The subregion with the highest percentage of these LPL synapses was layer 2-3 of the somatosensory cortex (17.2%), while the lowest percentage was found in the CA2 stratum lacunosum-moleculare (0.2%). Isocortex regions had the highest percentages of LPL synapses, with superficial layers showing 4- to 5-fold more LPL synapses than the deepest layers. Within the HPF, the subiculum had the highest percentage (9.6%) of LPL synapses of all subregions, followed by CA1sr (5.8%). These results show that the neocortex (isocortex), which is the most recently evolved region of the cortex as compared with the ancient allocortex (olfactory areas and HPF) (Rakic, 2009), contains higher numbers of LPL synapses, with the superficial layers of the neocortex being particularly well populated.

## Synapse protein lifetime increases throughout the lifespan

Synapse protein synthesis and turnover are thought to be important during brain development and aging (Hipp et al., 2019; Kaushik and Cuervo, 2015; Labbadia and Morimoto, 2015; Yerbury et al., 2016), yet to our knowledge there have been no studies of synaptic protein turnover comparing mice of different ages. We injected SiR-Halo into PSD95-HaloTag mice at three ages (3 weeks (W), 3 and 18 months (M)) that correspond to childhood, mature adulthood and old age (Cizeron et al., 2020).

These lifespan studies revealed that PSD95 protein lifetime increases in the majority of brain regions from birth to old age (Fig. 5A, 5B, S19-S21). At 3W there was a paucity of subregions showing long protein lifetimes, with only the most superficial layers of the neocortex and CA1 stratum lacunosum/moleculare showing a ^PSD95 density^t_1/2_ above that of other regions (Fig. 5A). Between 3W and 3M, the ^PSD95 density^t_1/2_ increased in all subregions, with the greatest increases in subregions of the isocortex and cerebellum granular layer (Fig. 5A, 5B, S20). We also examined age-dependent changes in PSD95 protein lifetime in the dendritic tree of CA1 pyramidal cells (Fig. 5C). At 3W there were nascent gradients in both basal and apical dendrites formed from synapses with short protein lifetimes, and their protein lifetime increased ∼5-fold by 3M. To uncover which synapse subtypes contribute to this increase we measured the effect size (Cohen’s *d*) for the change in density for all subtypes between 3W and 3M. This identified subtype 2, which is the LPL subtype with the longest protein lifetime, as significantly increased across many brain regions (Fig. S21A, C, E, G). By contrast, no increase was observed for subtype 34 (a type 3 LPL subtype; Fig. S21F, H), indicating that LPL synapses with different molecular compositions are non-redundant and have distinct developmental trajectories. During development, brain regions differentiate from each other by the acquisition of sets of diverse synapses (Zhu et al., 2018) and we therefore asked if brain regions differentiate with respect to their synapse protein lifetime compositions. Comparison of the ^PSD95 density^t_1/2_ between all subregions showed a high similarity at 3W that dramatically decreased by 3M (Fig. 5D). Together, these data show that from weaning to maturity the mouse brain accumulates synapses with long protein lifetimes, and that brain regions acquire a distinct and thus specialized compositional signature of synapses with different protein lifetimes.

**Fig. 5.**
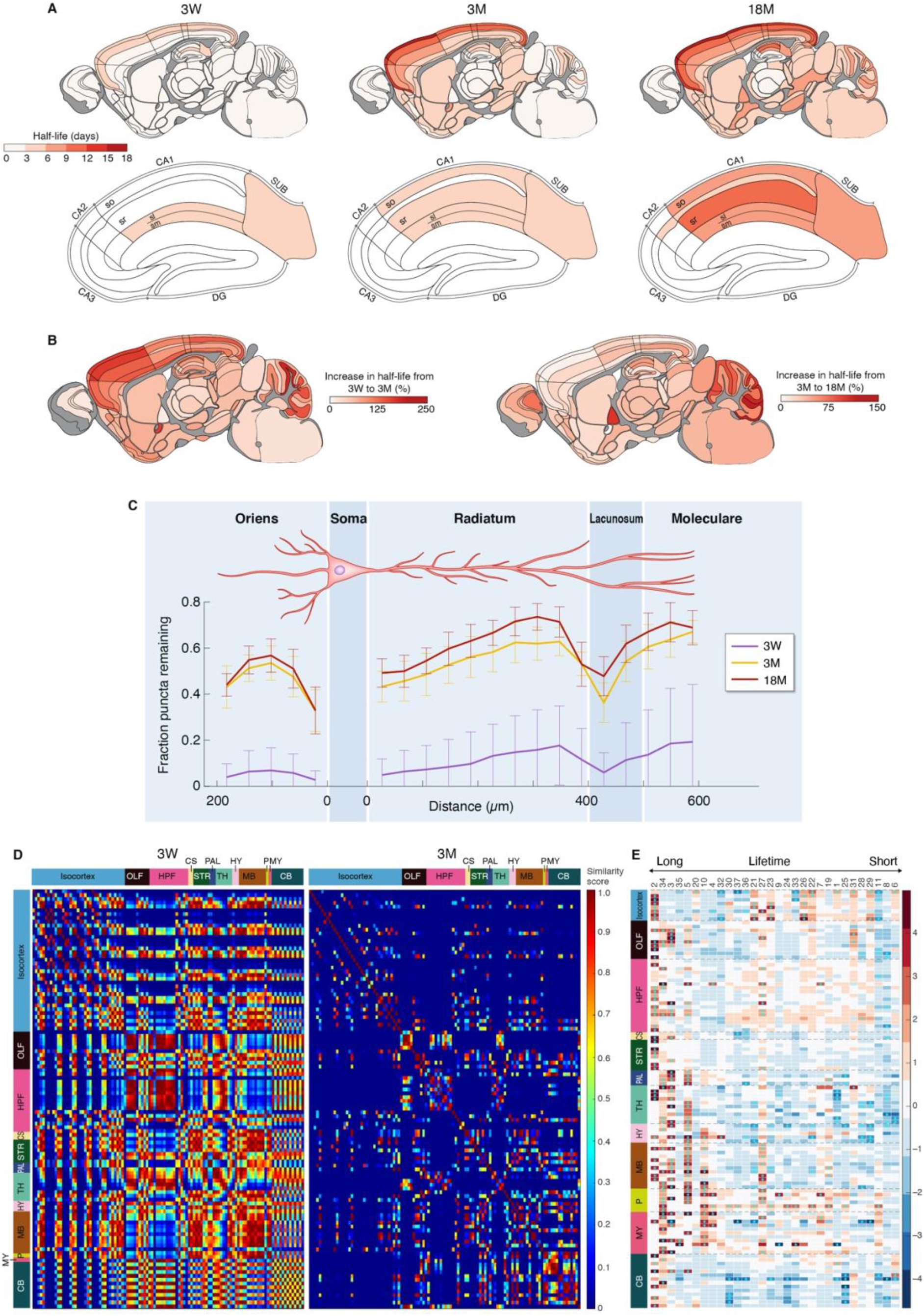
Synapse protein lifetime across the lifespan. (**A**) Summary of PSD95 puncta half-life across 110 mouse brain subregions in PSD95-HaloTag mice at 3 weeks (3W), 3 months (3M) and 18 months (18M) of age for the whole brain (top row) and HPF (bottom row). (**B**) Percentage increase in puncta half-life between 3W and 3M (left brain map) and between 3W and 18M (right brain map). (**C**) Mean percentage of puncta remaining at day 7 compared with day 0 (± SD) is plotted across the segments of basal and apical dendrites to cover the length of the CA1 dendritic tree at 3W, 3M and 18M (3 mice in each group). (D) Matrix of half-life similarities between pairs of subregions (rows and columns) for the 3W and 3M age groups. (**E**) Heatmap showing change (Cohen’s *d* effect size) in synapse subtype (subtype number shown at top) density across brain subregions between 3M and 18M. Subtypes are ranked with longest lifetime on the left and shortest on the right. Asterisks indicate significant differences (P<0.05, Benjamini-Hochberg correction). See Fig. S21 for detailed heatmaps.

We next examined how the brain changes with old age. Between 3M and 18M the regional ^PSD95 density^t_1/2_ increased in most areas of the brain (Fig. 5A, 5B, S20). By contrast, aging did not influence the spatial organization of synapses on the dendritic tree of CA1 pyramidal neurons, as the gradient and input-related pattern of synapse protein lifetime were maintained from 3M to 18M (Fig. 5C). Previously, we showed an overall loss of excitatory synapses between 3M and 18M but with a subtype specificity to the retention or loss (Cizeron et al., 2020). Here, we found that the increased regional ^PSD95 density^t_1/2_ in old age reflects the preservation of LPL synapses. Although most synapse subtypes decrease between 3M and 18M (Cizeron et al., 2020), the LPL synapses (subtypes 2, 34, 3, 5) were retained (Fig. 5E, S21B, D, I-L). Subtype 2 was preferentially retained in neocortex (isocortex), whereas in allocortex (olfactory cortex and HPF) subtypes 34, 3 and 5 were retained, revealing that LPL synapses with different molecular compositions are resilient to aging and populate distinct brain regions.

## Increased synaptic protein lifetime in *Dlg2* mutant mice

Mutations in the *DLG2* gene cause autism and schizophrenia in humans (Fernandez et al., 2009; Fromer et al., 2014; Ingason et al., 2015; Kirov et al., 2012; Nithianantharajah et al., 2013; Purcell et al., 2014; Reggiani et al., 2017; Ruzzo et al., 2019; Walsh et al., 2008) and the *Dlg2* mutant mouse model recapitulates the impairments in attention, learning and cognitive flexibility found in humans (Nithianantharajah et al., 2013). *Dlg2* encodes PSD93, a postsynaptic scaffold protein that binds to PSD95, and together these proteins assemble a ternary complex with the NMDA receptor subunit GluN2B (Frank et al., 2016). To test if *Dlg2* mutations affect the synaptome map of PSD95 lifetime we bred a cohort of PSD95-HaloTag mice containing heterozygous (*Dlg2*^+/−^;*Psd9*5^HaloTag/+^) or homozygous (*Dlg2*^−/−^;*Psd95*^HaloTag/+^) null alleles of *Dlg2* (Fig. 6, S22). Synaptome maps showed a robust increase in PSD95 protein lifetime across all 110 brain regions of both heterozygous and homozygous *Dlg2* mutant mice as compared with PSD95-HaloTag control mice, with a gene-dosage-dependent effect. PSD95 half-life increased up to 210% in heterozygous and 380% in homozygous *Dlg2* mutants. Most affected were subregions of the isocortex and HPF CA1, which are areas that show dysfunction in schizophrenia (Bähner and Meyer-Lindenberg, 2017; Heinz et al., 2019), intellectual disability and autism (Carper and Courchesne, 2005; Postema et al., 2019; Weston, 2019).

**Fig. 6.**
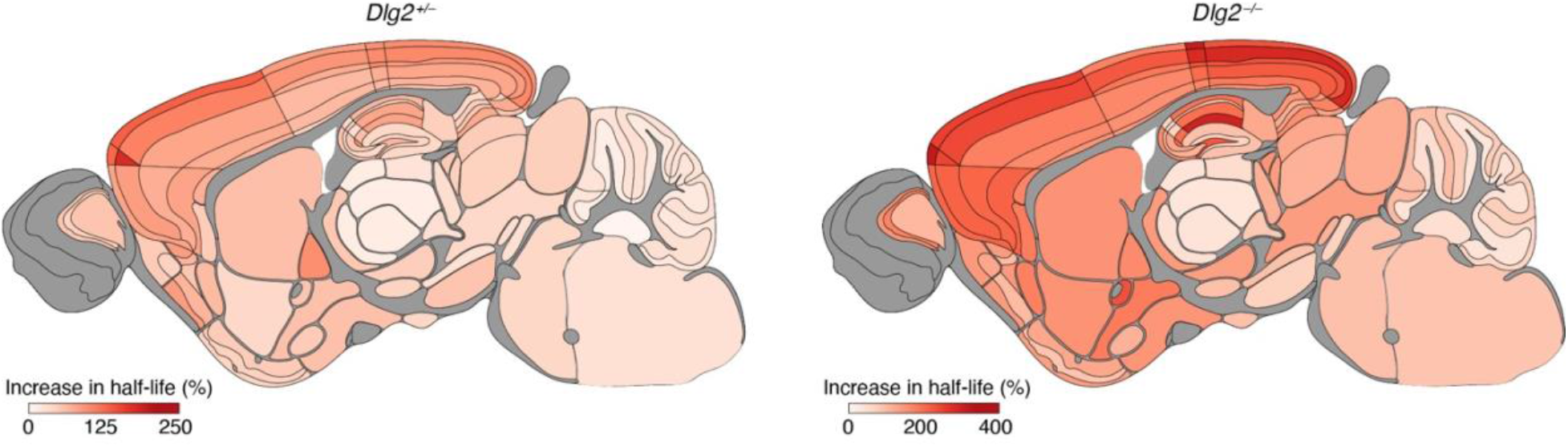
Synapse protein lifetime increases in *Dlg2* mutant mice. The percentage increase in PSD95 puncta density half-life across 110 adult (3M) brain subregions in mice carrying heterozygous or homozygous *Dlg2* mutations, as compared with control mice.

## Discussion

We systematically measured protein lifetime at single-synapse resolution across the brain and lifespan of the mouse. We focused on PSD95 because it is a highly abundant protein in the postsynaptic terminal of excitatory synapses and plays a crucial role in assembling glutamate receptors and other proteins into signaling complexes that control synaptic strength and plasticity (Carlisle et al., 2008; Fernandez et al., 2017; Fitzgerald et al., 2015; Frank et al., 2016; Migaud et al., 1998; Nithianantharajah et al., 2013). Excitatory synapses differed widely in their protein turnover rate and protein lifetime. Most excitatory synapses replaced PSD95 within a few days or weeks, but some could retain PSD95 for many months, indicative of a remarkable proteome stability. Synapse types 1 and 3, which are defined by their distinct protein compositions, each comprised a population of subtypes with a wide range of protein lifetimes. Thus, synapse diversity in the brain arises from differences in both proteome composition and protein lifetime.

### A brainwide architecture of synapse protein lifetime

When we looked at 30 excitatory synapse subtypes previously categorized by molecular and morphological parameters, they differed widely in their PSD95 protein lifetime. For simplicity, we designated the synapse subtypes with the longest protein lifetimes as LPL synapses and those with the shortest protein lifetime as SPL synapses. LPL and SPL subtypes are spatially distributed in all brain regions at differing abundance. Within individual dendrites, protein lifetime differed between adjacent synapses, was slower in distal dendrites and appeared to be influenced by synaptic inputs at different locations on the dendritic tree. Synapses within hippocampal subregions and layers showed different protein lifetimes, indicating projection and cell-type-specific regulation of turnover. These findings strongly support the conclusion that synapses with different protein lifetimes are building blocks of the synaptome architecture of the brain.

### A lifespan trajectory of synapse protein turnover

A striking finding was that the PSD95 turnover rate was highest in youngest animals and slowest in old age. This lifespan trajectory of slowing protein turnover and increasing protein lifetime occurs across almost all brain regions and has implications for understanding brain development, aging and disease. The enrichment of SPL synapses during early postnatal development could facilitate the rapid proteome remodeling that is instructed by dynamic transcriptome programs (Skene et al., 2017) and contribute to the dramatic expansion in synapse diversity in the first month of postnatal life (Cizeron et al., 2020). The slowing in protein turnover in old mice is consistent with the progressive reduction in protein clearance that underpins senescence and longevity (Kelmer Sacramento et al., 2020; Koyuncu et al., 2021). This reduction in protein clearance is known to contribute to the accumulation of toxic and misfolded proteins in neurodegenerative disorders (Labbadia and Morimoto, 2015; Yerbury et al., 2016) and our findings suggest that SPL and LPL synapses may have a differential vulnerability to these toxic proteins. In the future it will be important to uncover further molecular and biochemical characteristics of the SPL and LPL synapses and to consider the potential of targeted therapeutic interventions for neurodevelopmental and neurodegenerative disorders, and in senescence.

### Implications for synaptic physiology and behavior

Protein synthesis occurs within dendrites and the location of mRNA and ribosomes (Biever et al., 2020; Bingol and Schuman, 2006; Ostroff et al., 2002; Steward and Levy, 1982) may contribute to the dendritic distribution of synapses with different protein lifetimes. The lifetime of PSD95 in synapses may also be controlled by protein-protein interactions in the postsynaptic terminal. This is supported by our finding that in *Dlg2* mutant mice there was a marked increase in PSD95 lifetime. The spatial distribution of synapses with different protein turnover rates indicates that protein homeostasis (proteostasis) differs throughout the dendritic tree. The finding that synapses closest to the soma on apical dendrites of CA1 pyramidal neurons have the fastest turnover could potentially explain why homeostatic synaptic plasticity predominantly occurs in these synapses (Lee et al., 2013). Distal synapses, which have the slowest rates of turnover, would have slow homeostatic plasticity and maintain activity-dependent protein modifications for longer durations.

Our findings offer new insights into some long-standing issues in learning and memory research. Although it is accepted that synapses are the main locus of memory storage and that the initial step in programming a new memory is the activity-dependent modification of synaptic proteins within excitatory synapses, the mechanisms controlling the duration of memories and mechanisms of forgetting are less clear. In 1984 Crick speculated that the lifetime of synaptic proteins was too short relative to the duration of long-term memory, leading him to speculate about other mechanisms for long-term memory encoding (Crick, 1984). Our findings challenge Crick’s assumption that synaptic protein lifetime is far shorter than memory duration. We find that synapses vary widely in their protein lifetimes and that there are previously unknown subpopulations of synapses with very long protein lifetimes, potentially capable of maintaining copies of PSD95 for the lifetime of the mouse. A striking finding was that LPL synapses are heavily enriched in the neocortex where long-term memories are stored (Squire and Dede, 2015). Moreover, the lifetime of PSD95 in CA1, visual and piriform cortex corresponds well with the duration of spatial, visual and olfactory representations measured using calcium imaging of individual neurons (Driscoll et al., 2017; Rubin et al., 2015; Schoonover et al., 2021) and the duration of protein-synthesis-independent memories in dentate gyrus engram cells (Roy et al., 2017). Our data on synaptic protein lifetime at different ages also support the model that synapse protein lifetime is a mechanism relevant to memory duration. Very few, if any, long-term memories are retained from infancy (known as infantile amnesia) (Donato et al., 2021), and the capacity to form long-term memories increases during childhood and adolescence. Correspondingly, LPL synapses are scarce in 3W mice and, as they mature, they acquire increased numbers of LPL synapses. A hallmark of old age is the retention of long-term memories and a reduction in short-term and working memory capacity (Li et al., 2004; Reinert, 1970; Tucker-Drob, 2009), which is paralleled by our observations that LPL synapses are preferentially retained in old age and SPL synapses are lost. Finally, we found that *Dlg2* mutant mice show an increase in synaptic protein lifetime, which may be relevant to their inappropriate retention of learned information that impedes their cognitive flexibility (Nithianantharajah et al., 2013). A surprising and initially paradoxical finding was that those areas of the brain that are thought to be ‘hard wired’ and control innate behaviors were highly enriched in synapses with rapid protein turnover. The rapid protein turnover means that these regions have powerful proteostatic mechanisms, capable of erasing protein modifications and maintaining circuit homeostasis and thereby ensuring stable innate behaviors. Together, these findings raise the hypothesis that memory duration is dependent on the protein lifetime of synapses and that LPL synapses are specialized for storing information for long durations whereas SPL synapses retain modifications for short durations. Our proposal does not imply that other mechanisms do not contribute to long-term memory formation, such as Crick’s model of protein-protein interactions (Crick, 1984) or Kandel’s model of transcription-dependent new synapse formation (Kandel, 2004).

### Synaptic protein lifetime in disease

Genetic mutations that alter synapse protein synthesis and degradation are known to cause autism and other neurodevelopmental disorders (Louros and Osterweil, 2016). Such mutations might be expected to alter the architecture of synapse protein lifetime, as we have indeed demonstrated here for the autism and schizophrenia mutation *Dlg2*. Altering the anatomical distribution of synapses with different protein lifetimes could influence adaptive behaviors. In support of this, we have described changes in synaptome architecture in mouse models of autism and schizophrenia (Zhu et al., 2018), which have abnormal learning and impaired cognitive flexibility (Nithianantharajah et al., 2013). Many proteins that interact with PSD95 are encoded by genes conferring susceptibility to neurodevelopmental disorders including schizophrenia (Bayes et al., 2011; Fernandez et al., 2009; Fromer et al., 2014; Husi et al., 2000; Kirov et al., 2012; Purcell et al., 2014; Skene et al., 2017) and altering the lifetime of these proteins might contribute to disease phenotypes.

### The Protein Lifetime Synaptome Atlas

The atlas provides a versatile reference resource for understanding and modelling synaptic plasticity, memory, molecular and cellular anatomy, and brain disease. It can be integrated with other anatomical and molecular datasets, and expanded to encompass lifetime studies of further synaptic proteins, in both the pre- and post-synaptic terminal and in inhibitory synapses, and additional disease models. Data from the mouse atlas can be used to generate hypotheses for validation in other species. For example, it will be interesting to ask if the human neocortex is populated with synapses with longer protein lifetimes than those found in the mouse.

## STAR Methods

### Resource availability

#### Lead contact

Further information and requests for resources and reagents should be directed to and will be fulfilled by the lead contact, Seth Grant.

#### Materials availability

Newly generated materials are available from the lead contact upon request and, where appropriate, provision of a materials transfer agreement.

#### Data and code availability

- Synaptome map data have been deposited in Edinburgh DataShare and are publicly available as of the date of publication. The DOI is listed in the key resources table.
- All original code has been deposited at GitHub and is publicly available as of the date of publication. DOIs are listed in the key resources table.
- Any additional information required to reanalyze the data reported in this paper is available from the lead contact upon request.

### Experimental model and subject details

#### Gene Targeting and Mouse Generation

Animal procedures were performed in accordance with UK Home Office regulations and approved by Edinburgh University Director of Biological Services. The genetic targeting strategy adapted from Fernandez et al. (2009) was used to fuse HaloTag protein to the C-terminus of endogenous PSD95. HaloTag coding sequence (Promega) together with a short linker were inserted into the open reading frame of the mouse *Psd95* gene, followed by insertion of a loxP floxed PGK-EM7-neo-pA cassette, using recombination in *Escherichia coli*. E14Tg2a ES cells (from 129P2 ola) were used for gene targeting. Following identification of positive targeting clones using PCR, E3.5 blastocysts from C57BL/6 mice were injected with ES cells containing the target gene. Male chimeras were crossed with C57BL/6 females to produce first generation heterozygous animals. These mice were crossed with CAG Cre recombinase-expressing mice in order to remove the loxP floxed neo cassette in vivo. Animals were then cross-bred to produce a homozygous PSD95-HaloTag colony.

PSD95-HaloTag mice were genotyped by PCR using the following primers (5’-3’): exon F, GTCACATGTCTTTGTGACCTTG; 95UTR F, GATACATGCAGAGAGGAGTGTC; and Halogen F, CTGACTGAAGTCGAGATGGAC. Generation and characterization of the *Psd95*^eGFP/eGFP^;*Sap102*^mKO2/mKO2^ knock-in mouse line was described previously (Zhu et al., 2018). To establish the *Psd95*^HaloTag/eGFP^;*Sap102*^mKO2/y^ line, double homozygous *Psd95*^eGFP/eGFP^;*Sap102*^mKO2/mKO2^ mice were crossed with *Psd95*^HaloTag/HaloTag^ mice. Generation and characterization of the *Dlg2* mutant mouse line was described previously (McGee et al., 2001). To establish the *Dlg2*^+/−^;*Psd95*^HaloTag/+^ line, *Dlg2*^−/−^ mice were crossed with *Psd95*^HaloTag/HaloTag^ mice.

The mice were group housed and littermates were randomly assigned to experimental groups. The following groups of mice (m, male; f, female) were used for the study of PSD95 lifetime: 3M animals, day 0 n=9 (5m, 4f), day 1.5 n=8 (4m, 4f), day 3 and day 14 n=8 (3m, 5f), day 7 n=7 (2m, 5f); 3W animals, day 0 n=8 (5m, 3f), day 7 n=8 (6m, 2f); 18M animals, day 0 n=10 (7m, 3f), day 7 n=9 (8m, 1f). For triple colocalization the groups were: day 0 and day 7, n=8 (8f). For studies in *Dlg2* mutant animals, 3- to 4-month-old animals were used and the groups were: *Dlg2^+/+^*;*Psd95*^HaloTag/+^ animals, day 0 n=5 (2m, 3f), day 7 n=6 (2m, 4f); *Dlg2^+/−^*;*Psd95*^HaloTag/+^ animals, day 0 n=8 (6m, 2f), day 7 n=8 (3m, 5f); *Dlg2^−/−^*;*Psd95*^HaloTag/+^ animals, day 0 n=7 (3m, 4f), day 7 n=7 (2m, 5f). Sample size was estimated based on Cizeron et al. (2020).

### Method details

#### Western Blotting

PSD95 protein expression and abundance were tested in whole-brain synaptosome extracts from wild-type, *Psd95*^+/HaloTag^ and *Psd95*^HaloTag/HaloTag^ mice. Three extracts per genotype were loaded onto 4-12% gradient Bis-Tris gels, with 20 µl loaded per well. The gels were subjected to SDS-PAGE and western blotting. The membrane was probed with antibodies for PSD95, PSD95-HaloTag and alpha-Tubulin (loading control) and bands detected using the LI-COR Odyssey imaging system.

#### Electrophysiological recordings

Hippocampal slices obtained from the dorsal third of the hippocampus were prepared and maintained in vitro using previously described techniques (Babiec et al., 2017) approved by the Institutional Animal Care and Use Committee at the University of California, Los Angeles, USA. In experiments using extracellular recording techniques, slices were maintained at 30°C in an interface-slice type recording chamber perfused (2-3 ml/min) with an oxygenated (95% O_2_/5% CO_2_) artificial cerebrospinal fluid (ACSF) containing 124 mM NaCl, 4 mM KCl, 25 mM NaHCO_3_, 1 mM NaH_2_PO_4_, 2 mM CaCl_2_, 1.2 mM MgSO_4_, 10 mM glucose (all Sigma-Aldrich). Field EPSPs (fEPSPs) evoked by Schaffer collateral/commissural fiber stimulation were recorded in stratum radiatum of the CA1 region using low-resistance (5-10 MΩ) glass recording electrodes filled with ACSF (basal stimulation rate = 0.02 Hz). LTP was induced using two, one-second-long trains of 100 Hz stimulation (inter-train interval = 10 seconds). Whole-cell voltage-clamp recordings of evoked excitatory postsynaptic currents (EPSCs) and miniature EPSCs (mEPSCs) were performed using slices maintained in submerged-slice type recording chambers perfused with a modified ACSF containing 2.4 mM KCl, 3.0 mM CaCl_2_, 2.4 mM MgSO_4_, 100 µM picrotoxin. In these experiments, recording electrodes were filled with a solution containing 102 mM Cs-gluconate, 20 mM CsCl, 10 mM K-gluconate, 10 mM TEA-Cl, 5 mM QX-314, 0.2 mM EGTA, 4 mM Mg-ATP, 0.3 mM Na-GTP, 20 mM HEPES (pH 7.3, 290 mOsm). Evoked EPSCs (at 0.2 Hz) were recorded at membrane potentials of -80 mV and +40 mV and AMPAR-mediated and NMDAR-mediated components of the EPSCs were estimated by the amplitude of EPSCs 5 and 50 milliseconds after EPSC onset, respectively. Spontaneous mEPSCs were recorded at -80 mV in the presence of 1.0 µM TTX (Alomone Labs). Data collection and initial analysis were done blind to genotype.

#### HaloTag ligand generation

SiR-Halo coupling and purification were performed as previously described (Lukinavicius et al., 2013). 50 mg SiR-COOH dye (Spirochrome) and 36 mg Amine (O2) HaloTag building block (1.5 equivalent) were dissolved in 950 µl dimethylformamide (DMF). 90 µl N,N-diisopropylethylamine (DIPEA) were added into the solution. 70 mg benzotriazol-1-yl-oxytripyrrolidinophosphonium hexafluorophosphate (PyBOP) (1.5 equivalent) dissolved in 230 µl DMF was added to the SiR-COOH and Amine (O2) HaloTag solution. Solution was stirred at room temperature for 2 hours protected from light. Prior to purification, the compound was phase-separated from DMF. The compound was collected with the organic phase (diethyl ether) while the aqueous phase (brine) was discarded. Two to three tablespoons of dried magnesium sulphate were added to the organic phase solution, mixed well and filtered to remove any aqueous remains. Diethyl ether was evaporated, and the product was purified using silica gel column chromatography (2% MeOH in dichloromethane). The product yield: 63%.

#### In vivo application of HaloTag ligands

SiR-Halo ligand was dissolved in DMSO to a stock concentration of 5 mM. HaloTag ligand solution for injections was prepared as described (Grimm et al., 2017) but at 1.5 mM dye concentration and the total injection volume was adjusted by average weight for the animal group examined. 3W age group received 70 µl (21 µl HaloTag stock solution, 7 µl Pluronic F-127, 42 µl saline), 3M age group received 200 µl (60 µl HaloTag stock solution, 20 µl Pluronic F-127, 120 µl saline) and 18M age group received 300 µl (90 µl HaloTag stock solution, 30 µl Pluronic F-127, 180 µl saline) of HaloTag dye solution. Prior to injection, animals were placed in a heat box for 5-10 minutes to allow the blood vessels of the tail to dilate and become more visible. The mice were placed in a rodent restrainer for the injection. A bolus injection of HaloTag ligand solution was performed into the lateral tail vein. Following injection, the animals were monitored for any adverse effects twice daily for the length of the experiment. The experimenter was blinded to time points and genotypes at the time of injections.

#### Tissue collection and section preparation

Mice were fully anesthetized by intraperitoneal injection of 0.10-0.20 ml (according to age) pentobarbital (Euthatal). The thorax was opened and an incision made to the right atrium of the heart. A fine needle (26G, 0.45 x 100 mm) was inserted into the left ventricle and the animal was transcardially perfused with 10-15 ml 1xPBS followed by 10-15 ml fixative (4% paraformaldehyde (PFA)) and left at 4°C for 3-4 hours (according to age). Samples were transferred to a 30% sucrose solution and incubated for 48-72 hours at 4°C. Brains were then embedded in OCT (embedding matrix for frozen sections) solution (CellPath) inside a plastic mould (Sigma-Aldrich), the moulds placed in beakers containing isopentane (Sigma-Aldrich), and the beakers moved to a container containing liquid nitrogen for freezing. Frozen brains were stored at -80°C for up to 4-5 months. Frozen brain samples were cut at 18 µm thickness using a cryostat (NX70 Thermo Fisher) to obtain sagittal sections referring to 12-13/21 bregma level from the Allen Mouse Brain Atlas (sagittal, https://mouse.brain-map.org/experiment/thumbnails/100042147?image_type=atlas). Cut brain sections were placed on Superfrost Plus glass slides (Thermo Fisher). A drop of 1xPBS was placed on a glass slide prior to picking up the brain sections to ensure the brain tissue lay flat. After cutting, brain sections were left to dry in the dark at room temperature overnight and were then stored at -20°C. Frozen brain sections were placed in a dark chamber and incubated at room temperature for 1 hour prior to mounting the coverslips. Sections were washed with 800 µl 1xPBS to remove any remaining OCT on or around the brain tissue and allowed to dry. A 12 µl drop of MOWIOL solution was applied on top of the brain section. A glass coverslip (18 mm diameter, thickness #1.5, VWR) was carefully lowered on top of the sample to avoid any bubbles forming in between the glass slide and the coverslip. The sections were left to dry in the dark at room temperature overnight and then stored at 4°C for up to 1 week. The experimenter was blinded at the stage of tissue sectioning and section preparation for imaging.

#### Post-fixation labeling with HaloTag ligands

Brain sections were first washed with 800 µl PBS to remove any remaining OCT and left to dry in the dark. Hydrophobic marker pen was used to draw around sections to contain the solutions. 50 µl 10 µM TMR-Halo in 1xPBS was added to each brain section and samples incubated for 1 hour at room temperature in a wet dark chamber. Brain sections were then washed twice for 10 minutes each with 1xTBS (Tris buffered saline) containing 0.2% Triton X-100 detergent to remove any unbound TMR-Halo ligand and once with 1xTBS for 10 minutes.

#### CA1 pyramidal neuron filling

Mice (n=14) were perfused intracardially with 1xPBS followed by 4% PFA and, after removal, brains were subsequently post-fixed in PFA for 24 hours. After washing in PBS, brains were cut in coronal sections (200 µm thick) with a vibratome, and to identify cell nuclei the sections were prelabelled with 4’,6-diamidino-2-phenylindole (DAPI) (Sigma). Then, CA1 pyramidal neurons were individually injected with Alexa Fluor 488 dye solution (Life Technologies) by continuous current until the dendrites were filled completely as previously described (Benavides-Piccione et al., 2013). After injections, sections were mounted in ProLong Gold Antifade mounting medium (Life Technologies). Imaging was performed with a Zeiss LSM 710 confocal microscope. The fluorescence of DAPI, Alexa 488 and 633 (SiR-Halo-positive puncta) was recorded through separate channels. Stacks of images were acquired at high magnification (63x oil immersion; pixel size 0.057 x 0.057 µm; z-step 0.14 µm) and no pixels were saturated within the spines. For intracellular injection experiments, the animals were randomly assigned into groups for different time points for sacrifice.

#### Spinning disk confocal microscopy

Imaging was performed using an Andor Revolution XDi spinning disk microscope equipped with CSU-X1 (pinhole size 50 µm) and 2x post-magnification lens. Images of 512 x 512 pixels in size and 16-bit depth were obtained using Andor iXon Ultra back-illuminated EMCCD camera and Olympus UPlanSAPO 100x oil-immersion lens (NA 1.4). To cover the whole area of the sagittal brain section, multi-tile single-plane image acquisition was arranged with no overlap between adjacent tiles. Focus in z-plane was achieved by selecting and recording desired z-position for four points on the brain section, which were then used to calculate the intermediate z-positions for in-between image tiles.

#### Imaging parameters

High-magnification (x100) images at a single-synapse resolution covering the whole brain section were obtained using 250 EM gain, 2-frame averaging and 5000 millisecond acquisition speed. SiR-Halo was excited at 640 nm, TMR-Halo at 561 nm, eGFP at 488 nm and mKO2 at 561 nm. Emitted light was filtered with a QUAD filter (BP 440/40, BP 521/21, BP 607/34 and BP 700/45).

#### Long-term imaging in vivo

5-month-old male (n=3) and female (n=1) homozygous PSD95-HaloTag mice were used in accordance with the guidelines of the Federal Food Safety and Veterinary Office of Switzerland and in agreement with the veterinary office of the Canton of Geneva (license number GE12219A). To generate sparse expression of YFP in the supragranular layers of cortex, a mix of AAV9.CamKII.Cre.SV40 (2.4×10^13^ GC/ml; Addgene #105558) and AAV2.Ef1α.DIO.eYFP (4.6×10^12^ GC/ml; UNC GTC vector #42585) (100 nl in a ratio 1:1500) was injected in the primary somatosensory cortex. Cranial windows were implanted as described previously (Holtmaat et al., 2009) using a circular (3 mm diameter) coverglass pressed down onto the intact dura and glued to the skull using dental acrylic cement. 2-3 weeks after window implantation, SiR-Halo ligand was injected in the tail vein, as described above. Images were acquired in head-fixed mice under anesthesia (0.1 mg/kg Medetomidin and 5 mg/kg Midazolam, i.p.) using a custom-built, two-photon laser-scanning microscope (2PLSM) (Holtmaat et al., 2009) controlled by custom software written in MATLAB (Scanimage, Vidrio Technologies) (Pologruto et al., 2003). A tunable Ti:sapphire laser (Chameleon ultra II, Coherent) was used as a light source, tuned to λ=840 nm for simultaneous excitation of YFP and SiR-Halo ligand, or λ=910 nm for YFP alone. Excitation power was kept constant over time. Fluorescent images were collected using a 20x, 0.95 numerical aperture water-immersion objective (Olympus) and GaAsP photomultiplier tubes (10770PB-40, Hamamatsu). Emitted light was spectrally separated using a 565 nm dichroic mirror (Chroma) and two band-pass emission filters (ET510/50 nm, Chroma; 675/67 nm, Semrock). Field-of-views (FOVs) with YFP-labeled dendrites were first imaged at 6 hours post-injection, and then re-imaged after 1, 2, 3, 7, 10 and 14 days. Additional FOVs were imaged only once over the course of the experiment to measure population fluorescence decay similarly to the *in situ* experiments (Fig. S15). For each FOV, two separate image stacks (typically 10-30 planes, separated by 1 μm) were acquired at 2 milliseconds/line (image size, 1024 x 1024 pixels; pixel size, 0.06 x 0.06 μm), one at excitation λ=840 nm and one at λ=910 nm. The dim YFP signal that was collected at λ=840 nm served to align the dendrites in both image stacks.

### Quantification and statistical analysis

#### Analysis of electrophysiological data

Acquisition and analysis of electrophysiological data were performed using pCLAMP software (Molecular Devices). Results are reported as mean ± SEM.

#### Detection of synaptic puncta

Synaptic puncta detection from fluorescence images was performed using a machine learning-based ensemble method developed in-house by Dr Zhen Qiu (Cizeron et al., 2020; Zhu et al., 2018). The classifiers/detectors of PSD95eGFP and SAP102mKO2 were trained in a previous study (Zhu et al., 2018). The training of the image detector for PSD95-HaloTag was required. We first collected a training set of 249 images (10.8 × 10.8 µm) by randomly sampling across 12 main brain regions using bootstrapping. All puncta in the training set were manually located by three independent individuals of varying scientific expertise. Manual annotation of each punctum was then weighted for each individual. The ground truth was generated by weighted averaging over all individuals; an average weight greater than 0.7 was considered as true puncta. The ensemble learning was applied for training and a K-folder method was applied to validate and test the trained detector.

#### Measurement of synaptic parameters

Upon detection and localization, synaptic puncta were segmented by applying an intensity threshold. The threshold for each detected punctum was set adaptively at 10% of the height of its fluorescence intensity profile. A set of parameters were then quantified for each segmented punctum, including mean pixel intensity, size, skewness, kurtosis, circularity and aspect ratio (Zhu et al., 2018). Additionally, quantification of puncta density per unit area was performed. For calculations of PSD95 lifetime, primarily the measurements of puncta density and total fluorescence intensity content (punctum size x punctum mean intensity) were used.

#### SiR-Halo-positive puncta 3D reconstruction

The z-stacks of SiR-Halo and dye-filled neuron images were analyzed with 3D image processing software Imaris 7.6.5 (Bitplane). SiR-Halo-positive puncta were manually reconstructed along apical dendrites (stratum radiatum). For detailed information regarding 3D reconstruction of SiR-Halo-positive puncta, see Figure S13.

#### Analysis of long-term imaging

The images from λ=840 nm and λ=910 nm acquisitions were overlaid using the Fiji pairwise stitching algorithm (BigStitcher) (Hörl et al., 2019). The images at different time points were aligned using the ‘Correct_3D_Drift’ plugin in Fiji (Parslow et al., 2014). To correct for slight variations in excitation efficiency and image quality at different time points, the images were normalized using the YFP fluorescence signal from the dendrites, under the assumption that YFP levels did not substantially change over time, and background subtracted using the blood vessel interior as the baseline signal. For single SiR-Halo punctum decay analysis, the total fluorescence of each punctum at every time point was measured from the plane that yielded the brightest signal. For spine SiR-Halo puncta, the spines were selected in the YFP channel. YFP and SiR-Halo fluorescence intensity (from 910 nm and 840 nm excitation, respectively) was integrated in the spine head. To correct for bleaching due to repeated imaging of the puncta, on the first day of imaging a series of seven images were taken of a control FOV in rapid succession (within 5 minutes). For each point in the series the level of bleaching was estimated (∼3%) and applied as a correction factor to the corresponding time point in the long-term imaging series.

#### Colocalization of synaptic puncta

Object-based colocalization method was employed to assess double (between SiR-Halo and eGFP) and triple (between SiR-Halo, eGFP and mKO2) colocalization of markers of synaptic puncta. The double colocalization between two channels (SiR-Halo and eGFP) was obtained by minimizing the summation of the total centroid distance between all eGFP and SiR-Halo using linear programming (Zhu et al., 2018). The triple localization is computationally much more complex and an NP-hard problem, thus it cannot be resolved efficiently using linear programming. New multiple-channel localization algorithms were developed by adapting the multiple hypothesis tracking algorithm (Chenouard et al., 2013) to the particle tracking problem: the image channel of different markers were considered as temporal image frames in tracking, but more image matching between different markers was added because there was no temporal sequence in multilocalization. A distance threshold of 400 nm was applied to identify markers that were colocalized or triple localized. 400 nm is within the range of the previously measured diameters of postsynaptic density as observed by electron microscopy (Harris et al., 1992).

#### Classification of synaptic puncta

Using classifiers trained and built in a previous study (Cizeron et al., 2020), *Psd95*^eGFP/+^;*Sap102*^mKO2/y^ synapses were classified into 37 subtypes based on the parameters measured in the previous steps. A small percentage of puncta (∼0.002%) that did not get classified into any subtype group were denoted ‘other subtype’. The classification results were then combined with the triple localization result of PSD95-HaloTag to quantify the lifetime of different synapse subtypes over all brain regions/subregions at time points post SiR-Halo injection.

#### Segmentation of brain regions and subregions

An overview montage of each imaged sagittal brain section was stitched from ∼45,000-50,000 image tiles using a custom MATLAB script and downsized by a factor of 16. Quantification of PSD95-Halo labelling was performed on anatomical brain regions as defined by the Allen Mouse Brain Atlas. A combination of manual and semi-automated delineation methods was employed for defining 110 anatomical regions in each sagittal brain image. Manual delineations of subcortical brain regions (subregions of the olfactory areas, cortical subplate, striatum, pallidum, hypothalamus, thalamus, midbrain, cerebellum, pons, medulla) were performed using a polygon selection tool in Fiji/ImageJ (Schindelin et al., 2012; Schneider et al., 2012). Semi-automated delineations of isocortex and hippocampal formation were performed with in-house custom-built software. The program provides a user interface for a freeform deformation of a surface on which we map a specially prepared image that contains a multitude of delineated regions (Sederberg and Parry, 1986). The specially prepared images were, in most cases, delineation templates obtained from the Allen Mouse Brain Atlas. The delineation templates mapped on a digital surface are subdivided into a uniform grid so that each cell contains a subdivision of the delineation template. The software allows the user to manipulate the grid points, which results in the deformation of the grid cells and in turn leads to the deformation of the delineation template (Catmull, 1974). The software superimposes the delineation template on top of the brain image being delineated. The user can deform the delineation template to ‘fit’ the underlying montage image of the brain section. The software has a function allowing the user to paint over any damaged areas on the tissue to exclude them from the finalized delineation. The final delineation is saved as a set of .roi files representing individual brain regions, which are used in the later steps of the mapping pipeline.

#### Decay rate estimation

PSD95 decay rate was estimated by two alternative methods: (a) puncta count-based estimation and (b) total punctum intensity-based estimation. Method (a) relies purely on the presence or absence of fluorescent synaptic puncta, whereas method (b) considers the fluorescence content within each punctum. For both methods, the mean at day 0 was considered a reference point to which all the subsequent values were normalized. Single-phase exponential decay function was fitted to fractions of puncta/synapse fluorescence remaining:

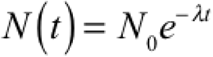

where N(t) denotes the puncta density/fluorescence intensity at time t, N_0_ = N(0) is quantity at t = 0 and λ is the decay rate constant.

##### Puncta count-based estimation

The detected number of fluorescent puncta per 100 µm^2^ (puncta density) was estimated for each ROI from every brain examined using the SYNMAP pipeline. Normalized puncta densities (y-axis) for each animal were plotted against time (x-axis) and an exponential decay function was fitted to estimate half-life and ± 95% confidence interval.

##### Total puncta intensity-based estimation

For each detected punctum within an ROI, mean fluorescence intensity was multiplied by punctum size to obtain the total fluorescence content for each punctum. The resulting fluorescence intensities were summed to obtain the total synaptic fluorescence content within a given brain region. Normalized fluorescence intensities (y-axis) for each animal were plotted against time (x-axis) and an exponential decay function was fitted to estimate half-life and ± 95% confidence interval.

##### PSD95 half-life calculations

PSD95 half-life was defined as the time required for the PSD95-Halo puncta density/fluorescence intensity to fall to half its initial value. Half-life was calculated based on the decay constant obtained from exponential curve fitting via the following formula:

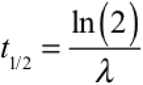

##### PSD95 fraction remaining calculation

In order to compare the decay of puncta number/fluorescence intensity between two time points, for each ROI we estimated the fraction of puncta/fluorescence intensity remaining at day 7 compared with day 0. The mean values for fraction remaining were calculated using:

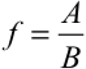

where A represents the mean of day 7 measurements and B is the mean of day 0 measurements. The error in the measurement of fraction remaining was estimated by following the statistical rules for propagation of uncertainty. The formula used for estimating the standard deviation of a ratio of means was:

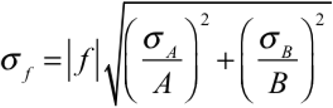

where σ_f_ is a standard deviation of the function f (ratio of mean (day 7)/mean (day 0)), σ_A_ is the standard deviation of A and σ_B_ is the standard deviation of B.

#### Statistical analysis

Covariance of pairs of synaptic parameters (density, half-life) was assessed using Pearson’s correlation function in MATLAB (Figs. S7 and S11). Correlation coefficient (R) values are presented alongside P-values that determine the statistical significance (P < 0.05) of the correlation between the two variables.

##### Bayesian analysis

Changes in subtype densities between 3M and18M were established in a previous study (Fig. S13 in Cizeron et al, 2020) and the heatmaps (Figs. 5E and S21) were then permutated column-wisely based on the ranking order of the subtype lifetime in Fig. 4A. The changes in subtype densities (Figs. 5E and S21) were tested using Bayesian estimation (Kruschke, 2013). Bayesian testing was also used to compare PSD95 lifetime between the *Dlg2* mouse mutant and control (Fig. S22). The input data were modelled assuming a t-distribution, and a Markov chain Monte Carlo algorithm was then performed to estimate the posterior distribution of the changes. P-values were then calculated for the distribution to complete the test. Each data point in each subregion was modelled and tested separately. The results were finally corrected over all subregions using the Benjamini-Hochberg procedure.

##### Cohen’s d effect size

The difference between two groups was measured by calculating a Cohen’s *d* effect size (Figs. 5E, S21 and S22), which is based on the difference between two means divided by their pooled standard deviation:

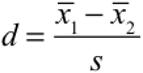

The pooled standard deviation was calculated as follows:

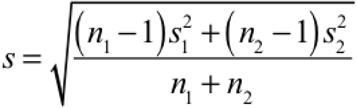

#### Self-organizing feature maps

Self-organizing feature maps (SOMs) (Kohonen, 1982) were constructed based on the subtype density of 110 subregions. SOMs preserve the local neighborhood so that subregions which are similar (neighbors) in the 37-dimensional subtype space become neighbors in the 2-dimensional output space. Network nodes were connected on a 2-dimensional rectangular grid of size 35 x 35 nodes. Input consisted of the 37 subtype densities of a subregion, where each subtype was normalized to [0, 1]. Number of training iterations was set to 45 and starting neighborhood size was set to 15 and reduced by a factor 0.9 for each iteration and weights were initialized to [0, 1]. After training, the strongest responding node was identified for each subtype and the position of this node was used for positioning the name of the subregion (Fig. S17A) (half-height of left-hand side of first letter of subregion name) as well as the position of the disc (disc center) representing the subregion (Fig. 4D, S17B). Radius of discs was proportional to the intensity half-life (^PSD95 intensity^t_1/2_) of the subregion represented by the disc. Colors of subregions used for presentations were the same as in Figure 2A.

#### Similarity matrices

Each row/column in the matrix represents one delineated brain subregion (Fig. 5D). Elements in the matrix are the synaptome similarities between two subregions quantified by differences in the PSD95 lifetime. A Gaussian kernel function was applied to convert the differences into similarities (Zhu et al., 2018).

#### Protein Lifetime Synaptome Atlas website

The Protein Lifetime Synaptome Atlas website (Bulovaite et al., 2021a) provides users with detailed information on PSD95 lifetime in regions and subregions of the mouse brain across the lifespan. The main panel shows the heatmap of protein half-life for the selected punctum parameter (density, intensity) and age or age comparison. The brain region navigator provides a collapsible tree structure that allows the user to pinpoint any subregion, its half-life values, and linked punctum parameter and synapse type/subtype prevalence data from the Lifespan Synaptome Atlas (Cizeron et al., 2020). Half-life data for further synaptic proteins will be added as they become available.

#### Additional resources

All brain maps generated for the study are freely available at (Bulovaite et al., 2021).

## Acknowledgments

**General:** Sarah Lempriere, Cathy McLaughlin, Emma Sigfridsson, Rand Dahan, Gabor Varga, Emily Robson, Theresa Wong, Bev Notman, Ian Hawes, Dimitra Koukaroudi, William Mungall, Lorena Tapia Mendive, Fabio De Moliner, Marc Vendrell and Babis Koniaris for technical assistance. Colin Davey for editing. Debbie Maizels for artwork. Trevor Robbins, Szu-Han Wang, Tim Bussey for comments on the manuscript.

## Author contributions

EB and AZ planned the experiments, collected, imaged and analyzed brain samples, performed image and data analysis. ZQ developed software and performed image analyses. MK analyzed protein extracts and developed methods. NHK, DGF and EJT constructed the PSD95-HaloTag mice. RG built the Protein Lifetime Synaptome Atlas website and performed statistical analysis. SAJ and TJO performed electrophysiology. PMS and JD performed neuron filling. EH and AH performed in vivo neuronal imaging. EF provided advice on experimental design and analyzed data. SGNG conceived the project and wrote the paper with input from all authors.

## Data and materials availability

Data are available at the Protein Lifetime Synaptome Atlas website (Bulovaite et al., 2021a) and Edinburgh DataShare (Bulovaite et al., 2021b).

## SUPPLEMENTAL INFORMATION

**Fig. S1. Related to Fig. 1.**
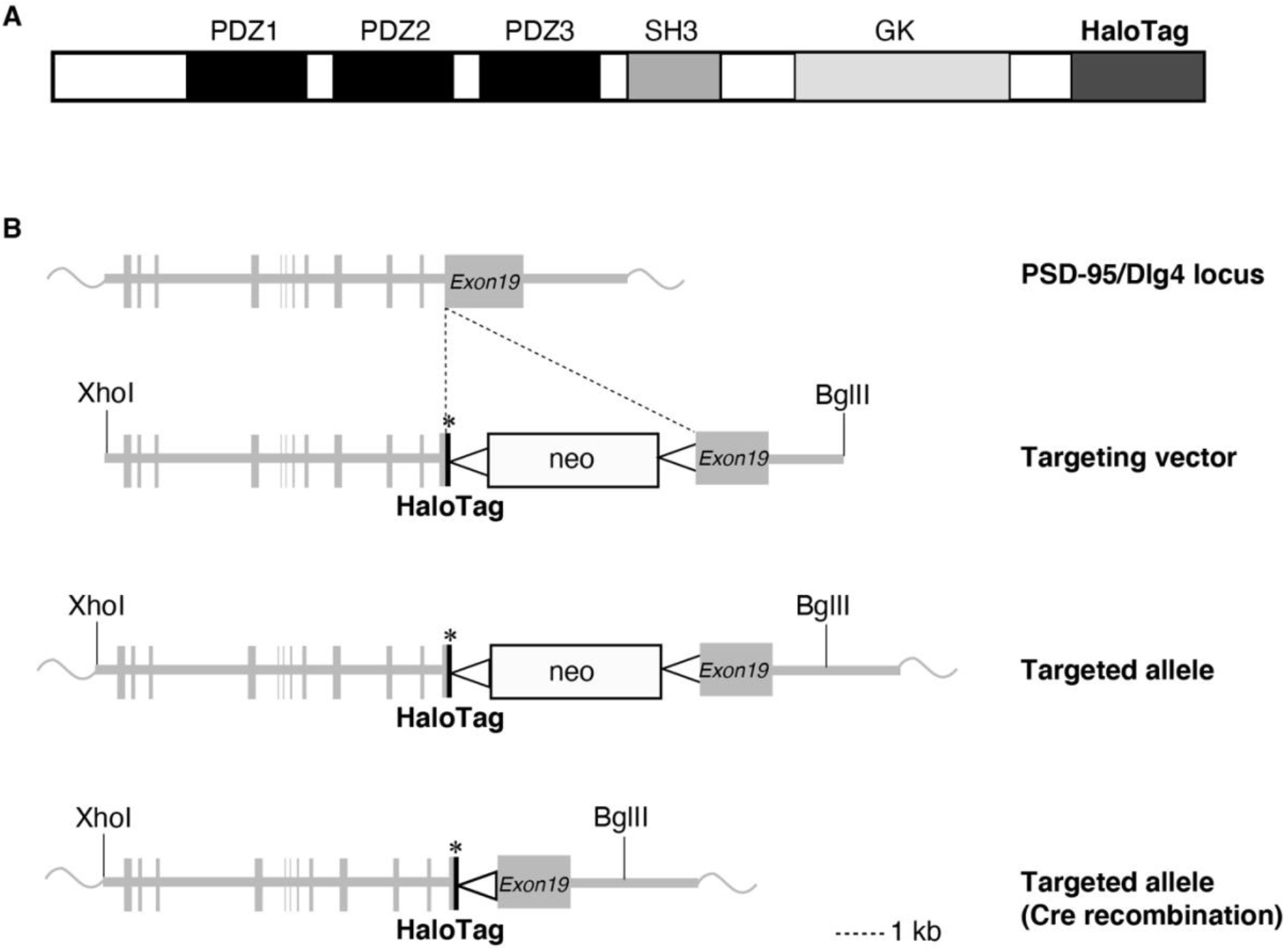
Generation of the PSD95-HaloTag knock-in mouse. (**A**) PSD95-HaloTag protein domain structure. PSD95 contains three PDZ domains followed by SRC homology 3 (SH3) and guanylate kinase (GK) domains. The HaloTag is fused at the C-terminus. (**B**) Targeting strategy used for the genomic *Psd95* (*Dlg4*) locus. The *Dlg4* allele was targeted with the HaloTag sequence inserted before the stop codon (asterisk). The neomycin resistance cassette (neo) was deleted between loxP sites (triangles) by crossing the PSD95-HaloTag mouse with a transgenic Cre recombinase-expressing mouse.

**Fig. S2. Related to Fig. 1.**
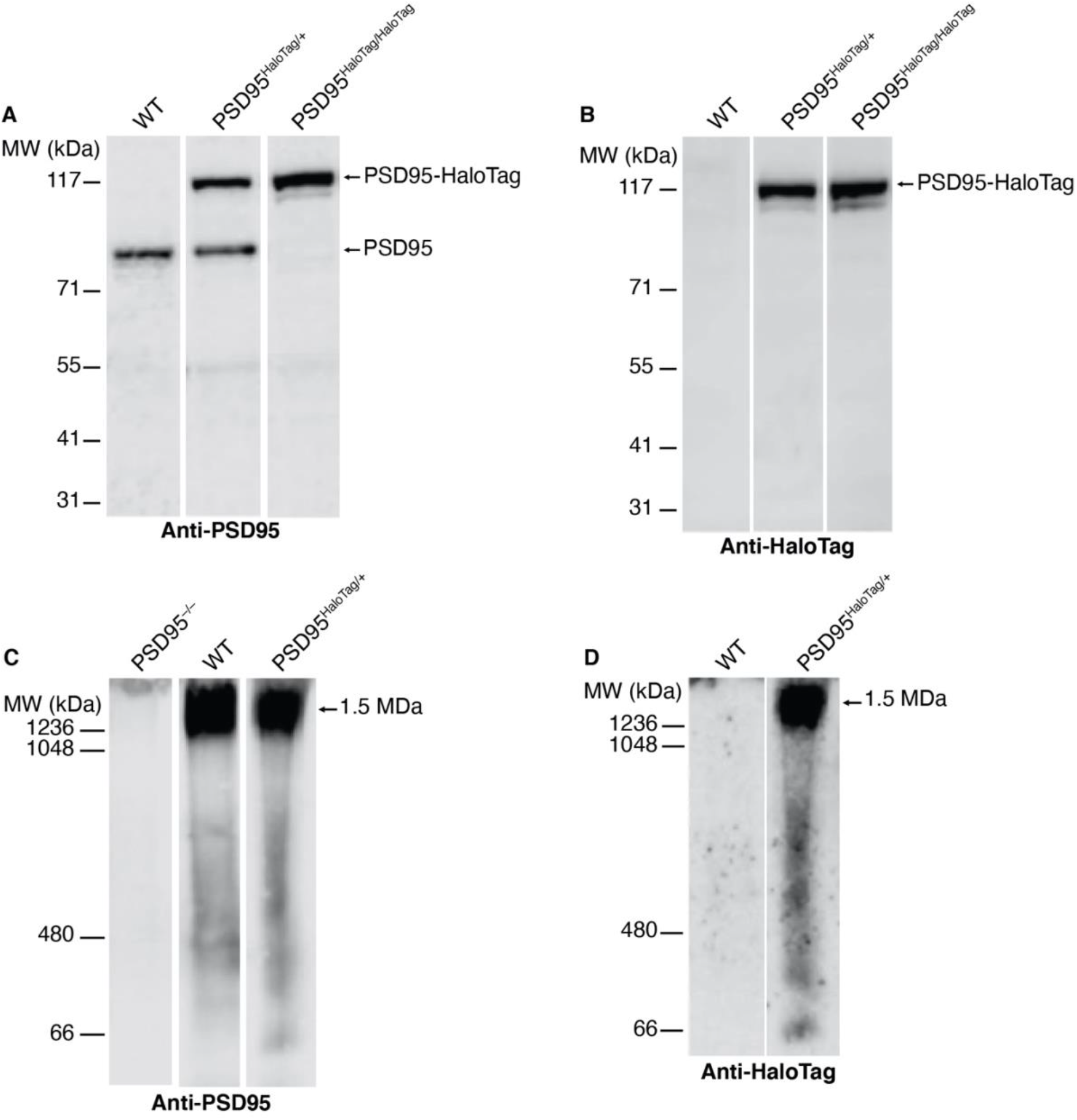
PSD95 expression is unaffected by the HaloTag. (**A**) Western blot analysis on crude synaptosome preparations from wild-type (WT), *Psd95*^HaloTag/+^ and *Psd95*^HaloTag/HaloTag^ mice using antibody against PSD95. PSD95 and PSD95-HaloTag fusion protein are indicated. (**B**) The same set of synaptosome preparations as in (A) were probed with anti-HaloTag antibody. Only PSD95-HaloTag protein is detected. (**C**,**D**) PSD95-HaloTag is assembled into 1.5 MDa postsynaptic supercomplexes. Complexes were separated using Blue native PAGE of the synaptosome preparations described in (A,B) and immunoblotted with antibodies to PSD95 (C) and HaloTag (D).

**Fig. S3. Related to Fig. 1.**
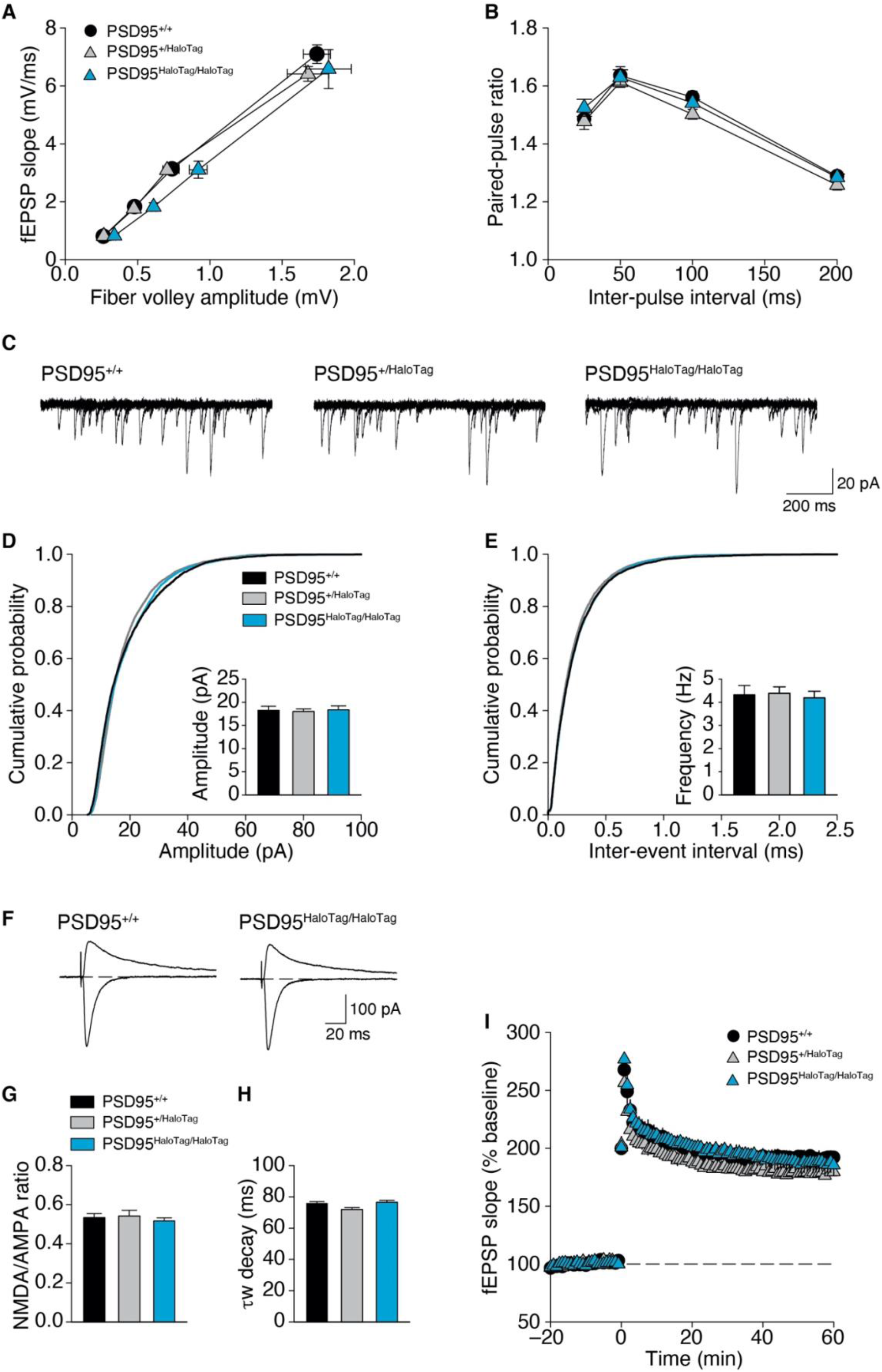
PSD95-HaloTag knock-in mice are electrophysiologically normal. (**A**) Fiber volley/field excitatory postsynaptic potential (fEPSP) input/output curves generated by eliciting fEPSPs with intensities of presynaptic stimulation corresponding to 100, 75, 50, and 25% of max. (**B**) Paired-pulse facilitation is normal in PSD95-HaloTag mutants. Results in (A) and (B) from 5 WT (20 slices), 5 heterozygous (21 slices), and 4 homozygous mutant (18 slices) mice. (**C**) Example mEPSCs (five superimposed 1-second-long recordings for each) showing no difference between WT and mutant. (**D**,**E**) Cumulative probability plots of mEPSC amplitudes (D) and inter-event intervals (E). *Psd95*^+/+^ (WT) n=4 (11 cells), heterozygous (*Psd95*^+/HaloTag^) n=3 (11 cells), homozygous (*Psd95*^HaloTag/HaloTag^) mutant n=4 (15 cells) mice. One-way ANOVA, F(2,8) = 0.049, P = 0.952 for mEPSC amplitude, and F(2,8) = 0.836, P = 0.921 for mEPSC frequency. (**F**) Example excitatory postsynaptic currents (EPSCs) recorded at membrane potential of -80 and +40 mV illustrate similarity of WT and homozygous mutants. (**G**) Ratio of NMDAR- to AMPAR-mediated currents recorded at V_m_ = +40 mV. No difference was detected between WT and mutants (one-way ANOVA, F(2,11) = 0.345, P = 0.717). (**H**) Weighted decay time constants for currents elicited at V_m_ = +40 mV. No difference in decay times was detected (F(2,11) = 3.602, P = 0.071). In (G) and (H), WT n=5 (19 cells), heterozygous n=3 (12 cells), homozygous mutant n=4 (16 cells) mice. (**I**) LTP induced by two 1-second-long trains of 100 Hz stimulation (delivered at time = 0, inter-train interval = 10 seconds). Sixty minutes post-100 Hz stimulation, fEPSPs were potentiated to 191 ± 4.5% of baseline in WT slices (*n*=10 slices from 5 mice), 179 ± 1.3% of baseline in heterozygote slices (*n*=10 slices from 5 mice), and 186 ± 8.2% of baseline in homozygous mutant slices (*n*=9 slices from 4 mice). One-way ANOVA, F(2,11) = 1.706, P = 0.226. Dashed line represents baseline synaptic strength. All panels report mean ± SEM.

**Fig. S4. Related to Fig. 1.**
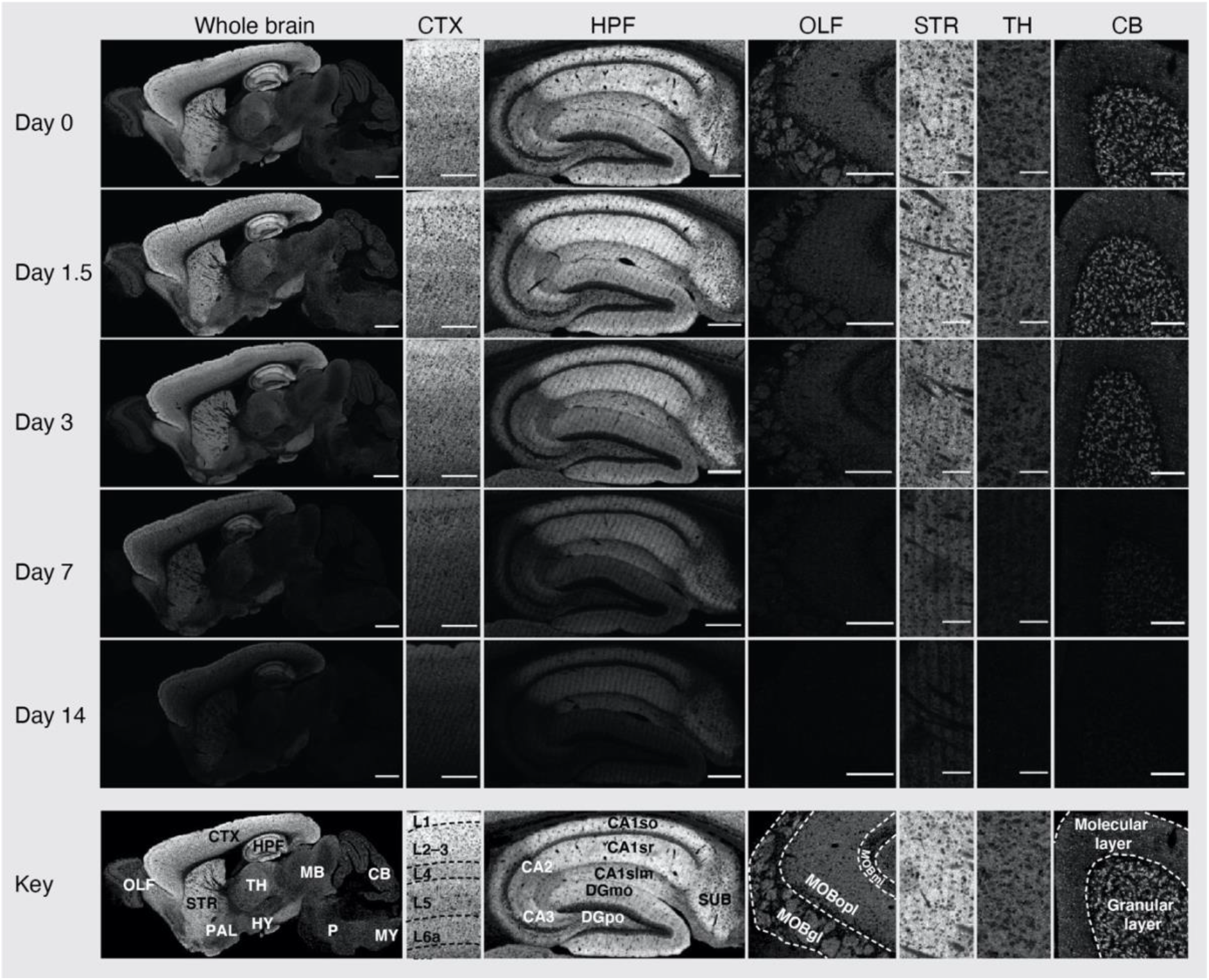
SiR-Halo fluorescence signal in different brain regions across time. Representative images of SiR-Halo-labeled PSD95-HaloTag expression in the whole brain, isocortex (CTX), hippocampal formation (HPF), olfactory areas (OLF), striatum (STR), thalamus (TH) and cerebellum (CB) at the indicated days post-injection of SiR-Halo. Scale bars: 2 mm (whole brain), 500 µm (CTX, HPF, OLF), 200 µm (STR, TH, CB). PAL, pallidum; HY, hypothalamus; MB, midbrain; CB, cerebellum; P, pons; MY, medulla; CA1so, CA1 stratum oriens; CA1sr, stratum radiatum; CA1slm, stratum lacunosum-moleculare; DGmo, dentate gyrus molecular layer; DGpo, dentate gyrus polymorphic layer; SUB, subiculum; MOBmi, mitral layer of the main olfactory bulb; MOBopl, outer plexiform layer of the main olfactory bulb; MOBgl, glomerular layer of the main olfactory bulb.

**Fig. S5. Related to Fig. 1.**
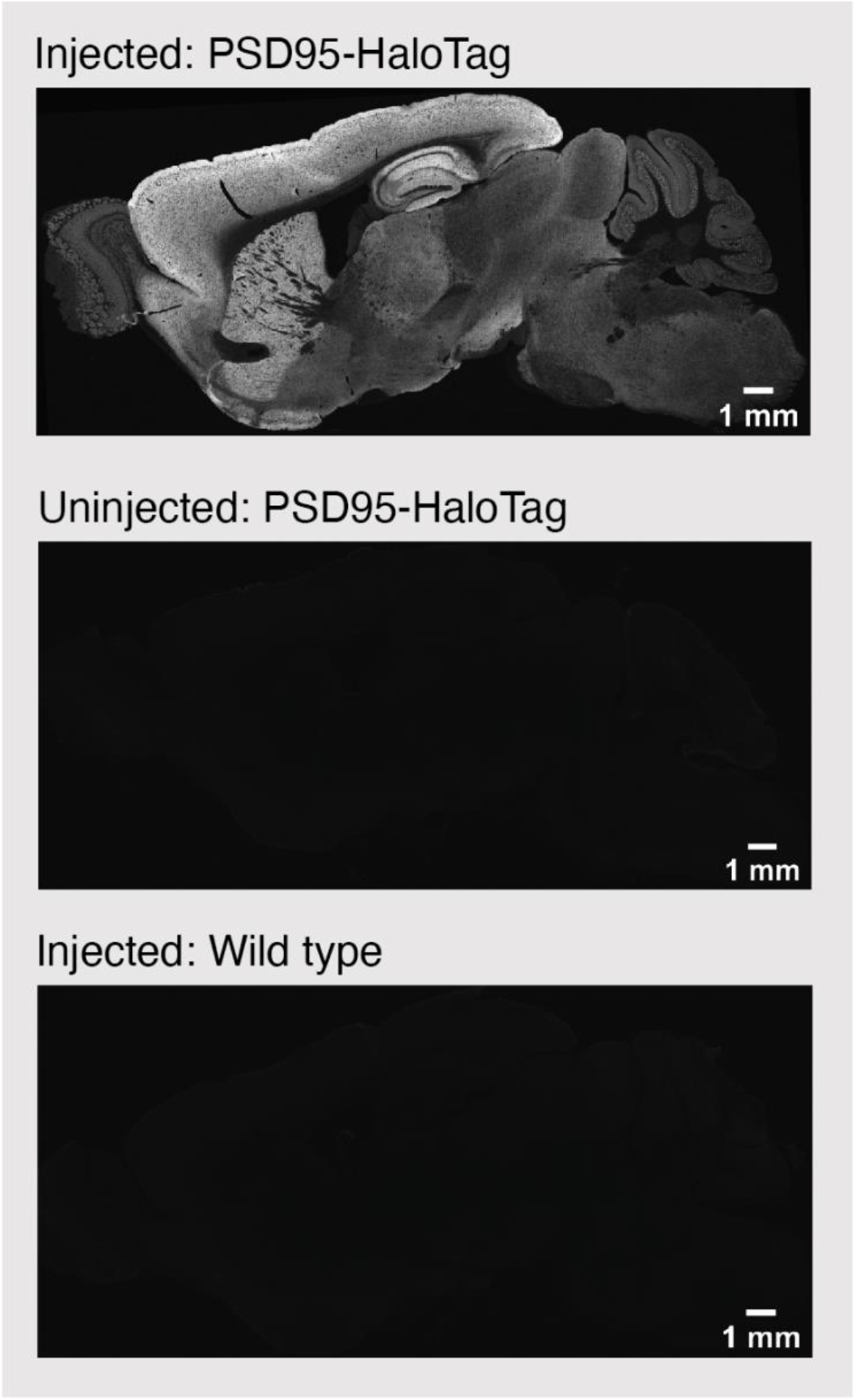
Specificity of labeling with SiR-Halo ligands. Fluorescence images of sagittal brain sections from a SiR-Halo-injected PSD95-HaloTag mouse (top), an uninjected PSD95-HaloTag mouse (middle), and SiR-Halo-injected wild-type mouse (bottom). Scale bars: 1 mm.

**Fig. S6. Related to Fig. 1.**
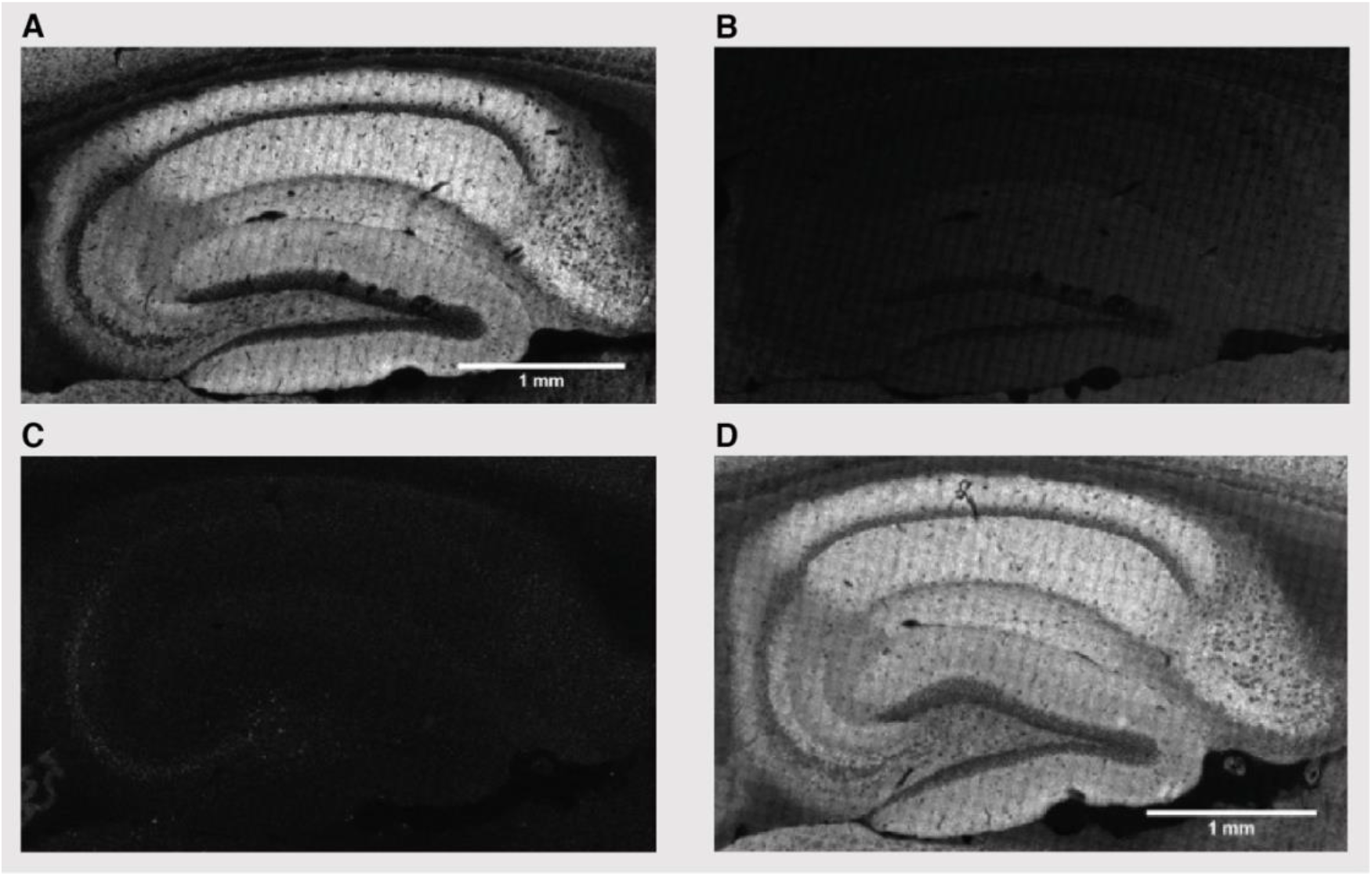
Saturation of labeling by injection of SiR-Halo. (**A**) HPF section from a PSD95-HaloTag mouse injected with 300 nmol SiR-Halo (6 hours post-injection). (**B**) Same section as in (A) stained with TMR-Halo after fixation. The absence of labeling indicates that SiR-Halo has occupied all available Halo ligand binding sites. (**C**) HPF section from a PSD95-HaloTag mouse that was not injected with SiR-Halo ligand. (**D**) Same section as in (C) stained with TMR-Halo after fixation. Note the strong labeling of TMR-Halo. Scale bars: 1 mm.

**Fig. S7. Related to Fig. 1.**
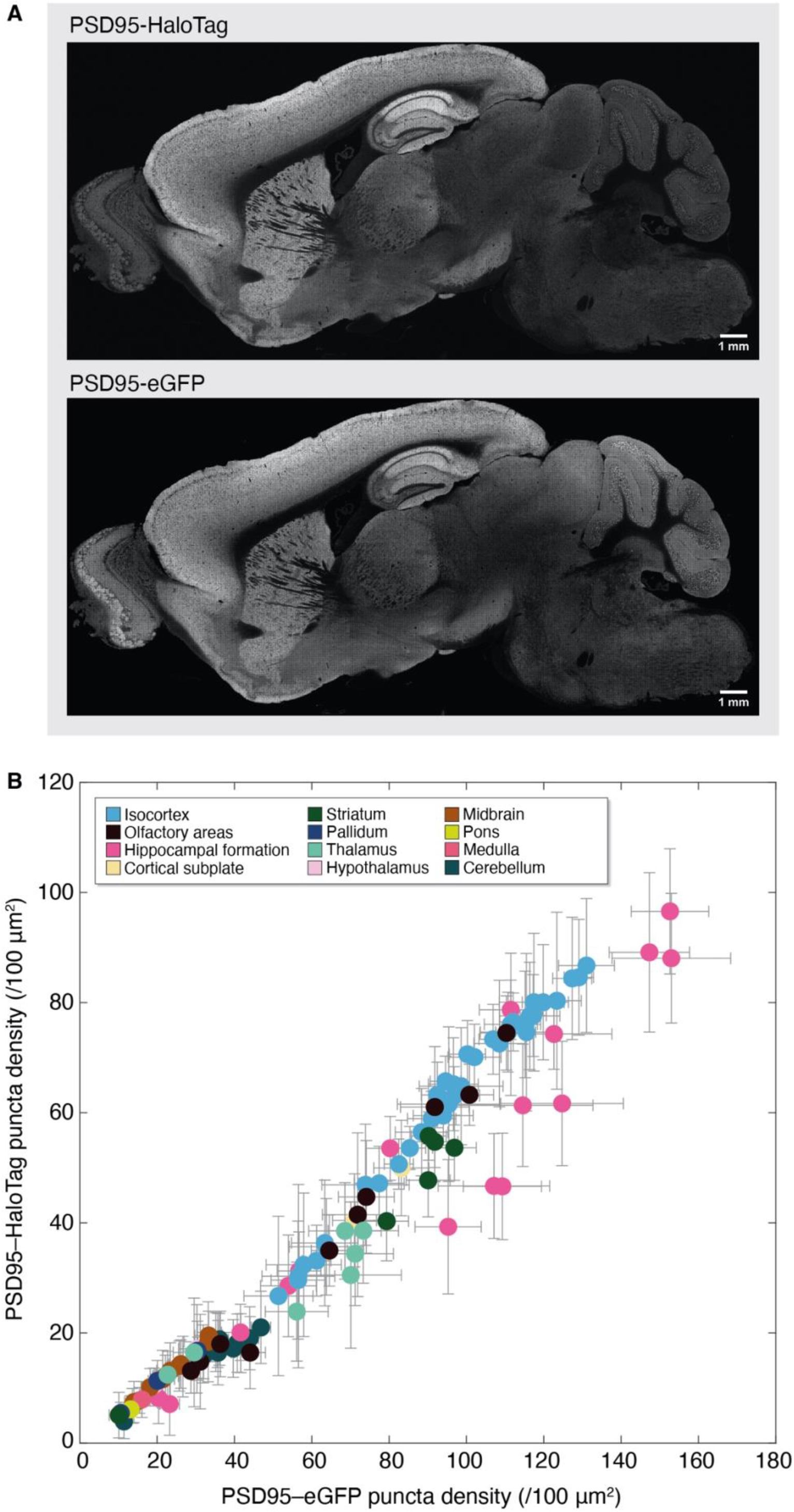
PSD95 fluorescence labeling in *Psd95*^HaloTag/eGFP^ mice. (**A**) Representative fluorescence images of a sagittal section showing PSD95-HaloTag and PSD95eGFP expression (fluorescence) in SiR-Halo-injected *Psd95*^HaloTag/eGFP^ compound heterozygous mouse. Scale bars: 1 mm. (**B**) Correlation of fluorescent puncta densities detected by PSD95eGFP (x-axis) and PSD95-HaloTag (y-axis) across 110 brain subregions (± SD). Each dot represents a brain subregion, and the color corresponds to the overarching brain area. R and P values are from the Pearson’s correlation test.

**Fig. S8. Related to Fig. 1.**
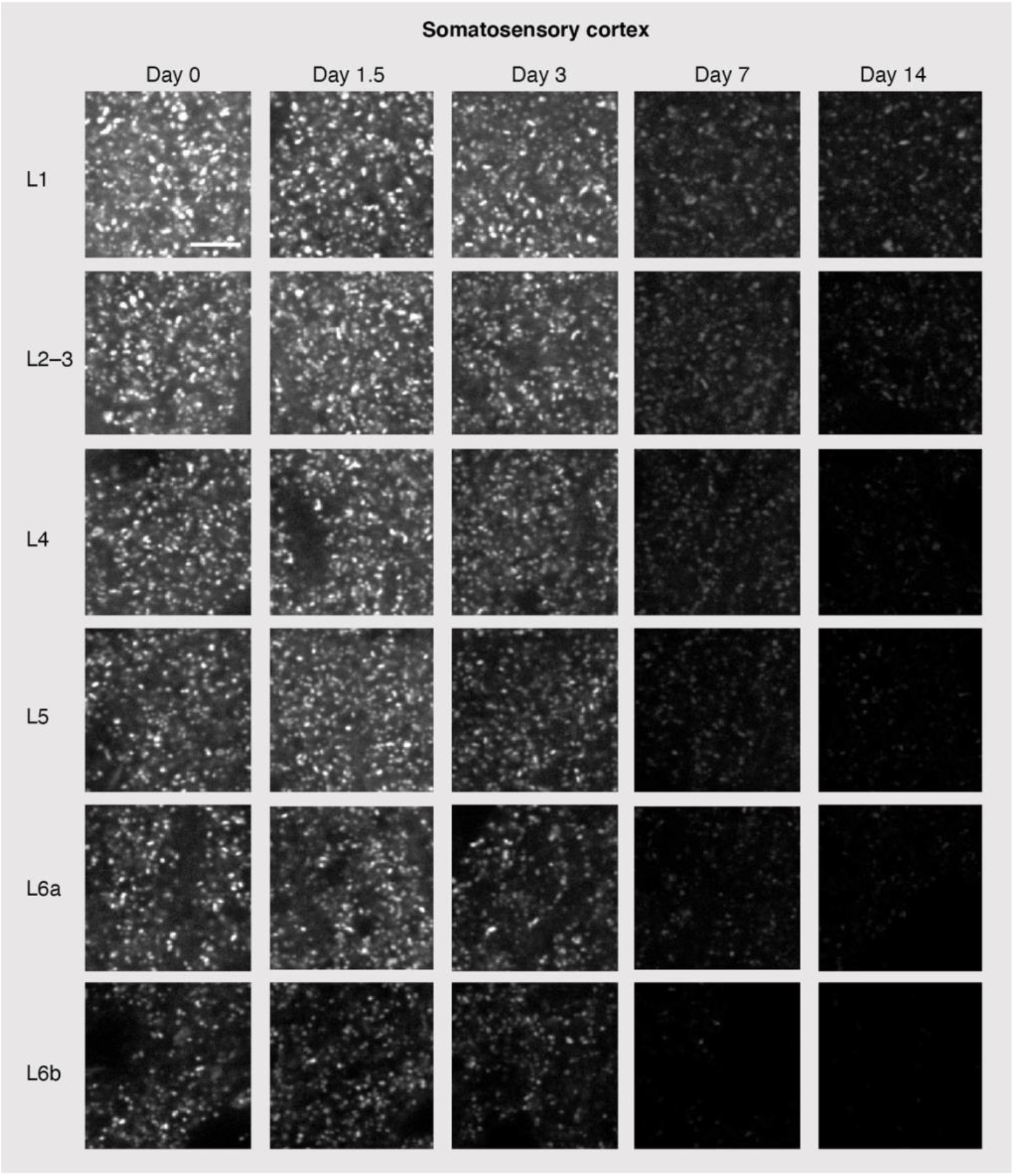
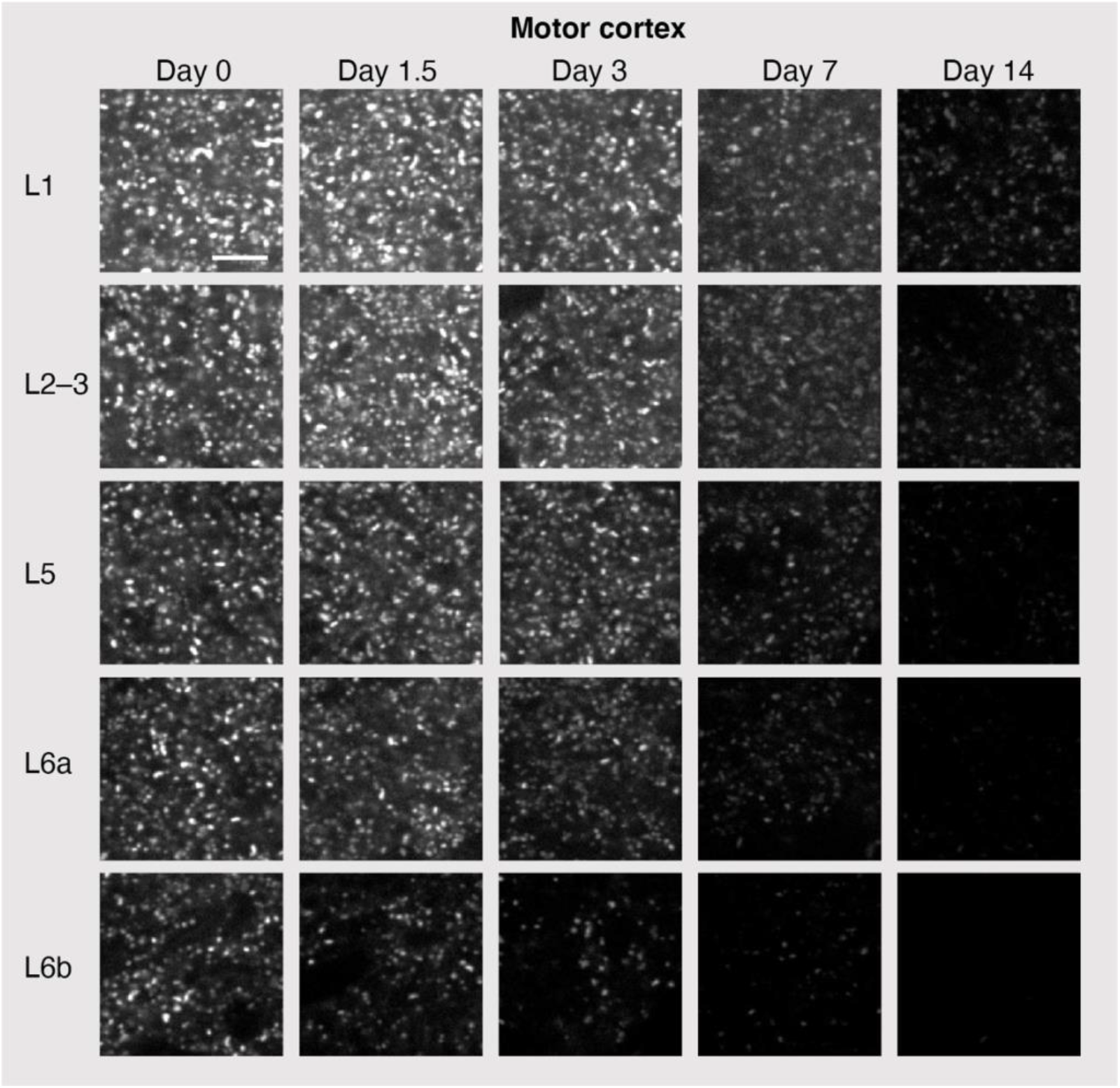
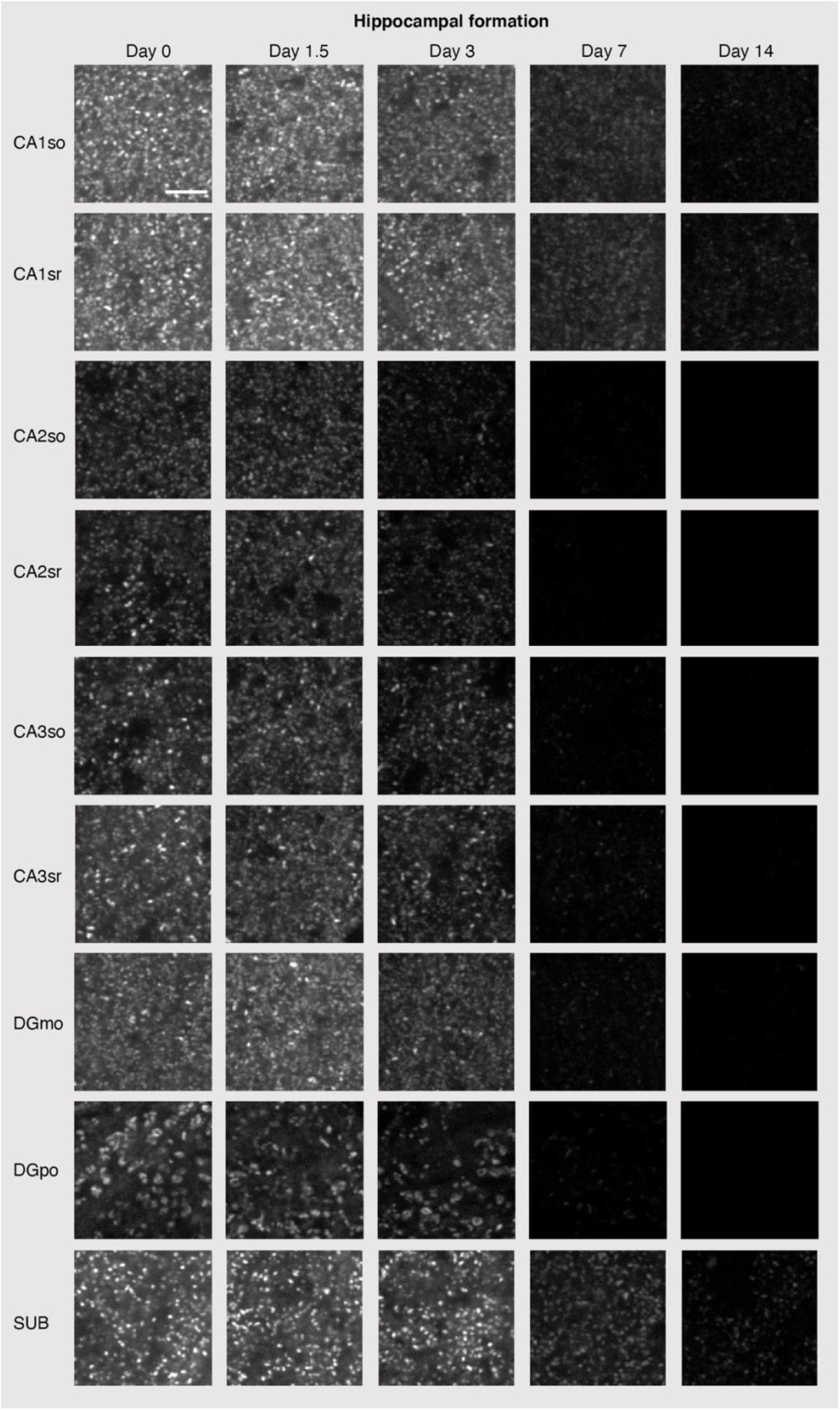
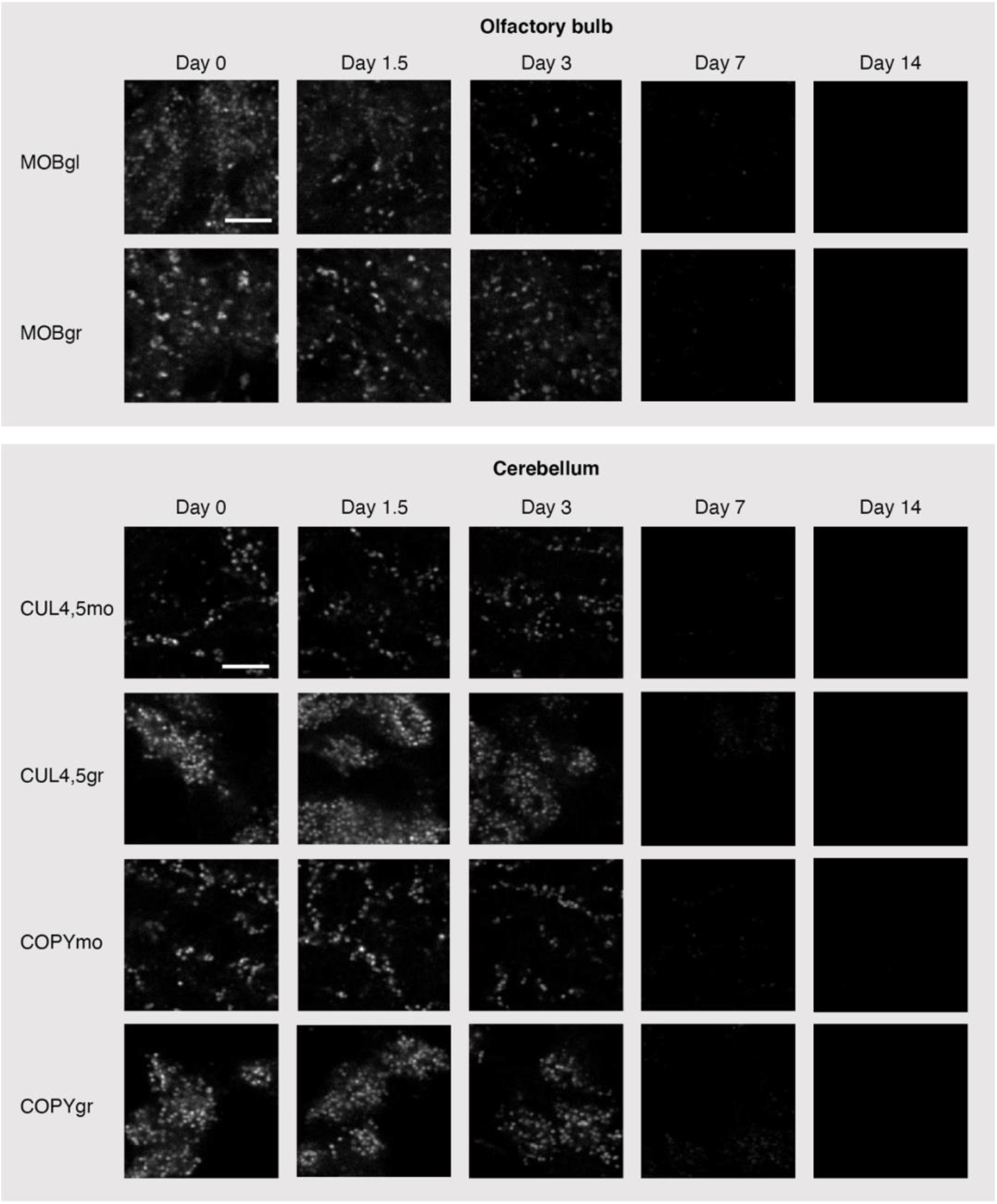
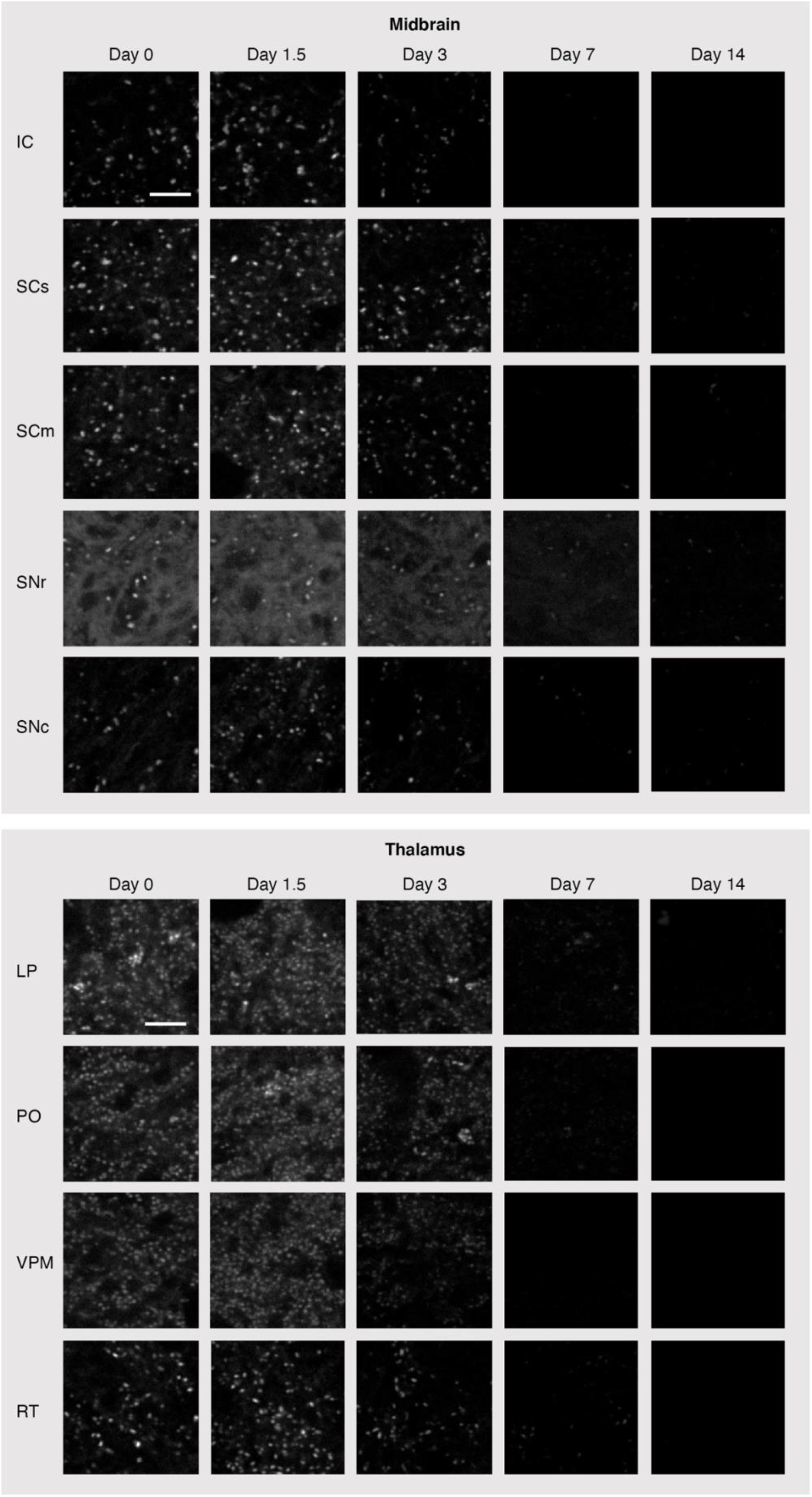
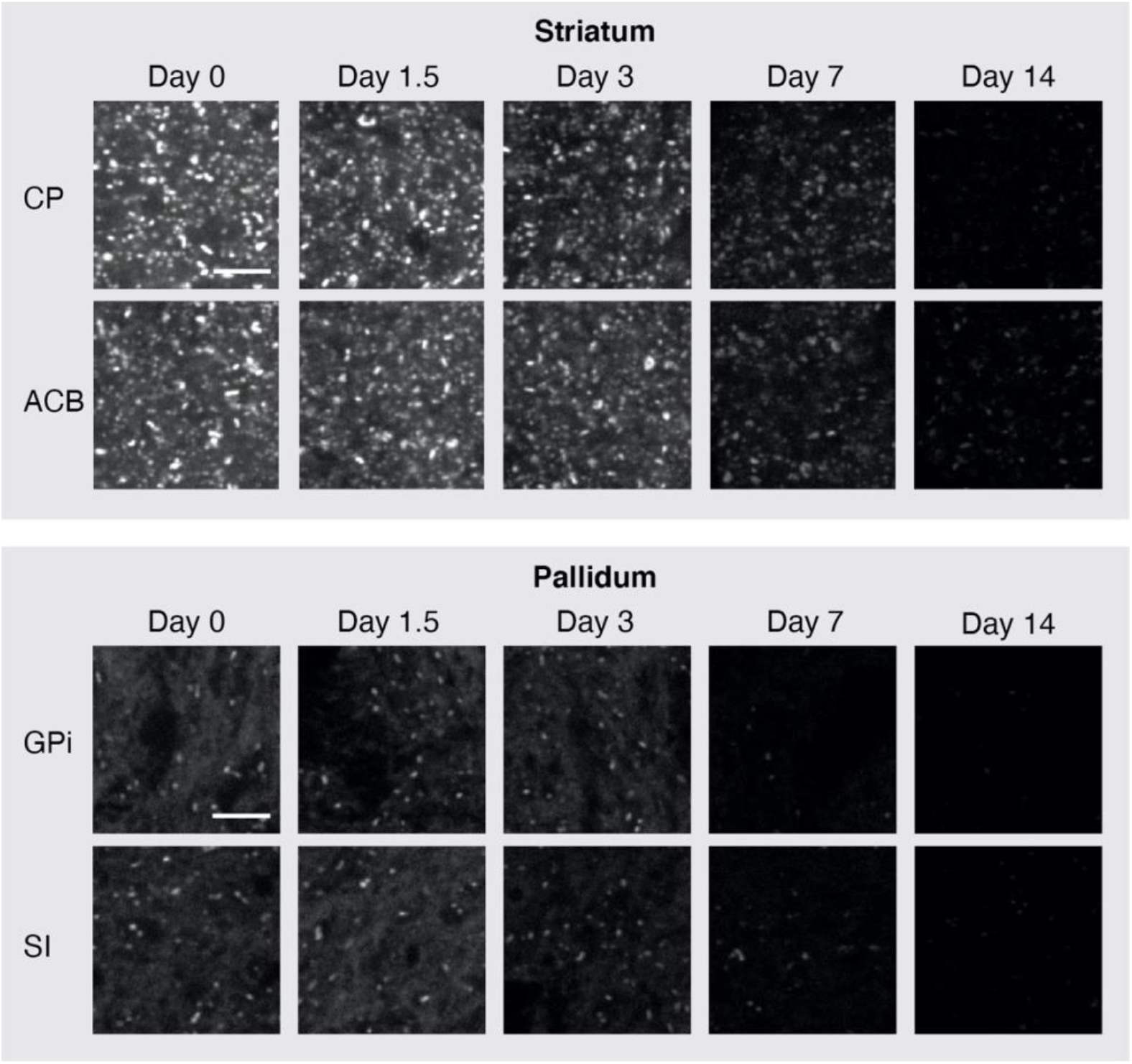
Fluorescent puncta decay in different brain regions and subregions. High-magnification example images of SiR-Halo puncta decay across 2 weeks in regions and layers of the isocortex (somatosensory and motor cortex), subregions of the hippocampal formation, olfactory bulb, cerebellum, midbrain, thalamus, striatum and pallidum. CA1so, CA1 stratum oriens; CA1sr, CA1 stratum radiatum; CA2so, CA2 stratum oriens; CA2sr, CA2 stratum radiatum; CA3so, CA3 stratum oriens; CA3sr, CA3 stratum radiatum; DGmo, dentate gyrus molecular layer; DGpo, dentate gyrus polymorphic layer; SUB, subiculum; MOBgl, glomerular layer of the main olfactory bulb; MOBgr, granular layer of the main olfactory bulb; CUL4,5mo, lobules IV-V molecular layer; CUL4,5gr, lobules IV-V granular layer; COPYmo, copula pyramidis molecular layer; COPYgr, copula pyramidis granular layer; IC, inferior colliculus; SCs, superior colliculus, sensory related; SCm, superior colliculus, motor related; SNr, substantia nigra, reticular part; SNc, substantia nigra, compact part; LP, lateral posterior nucleus of the thalamus; PO, posterior complex of the thalamus; VPM, ventral posteromedial nucleus of the thalamus; RT, reticular nucleus of the thalamus; CP, caudoputamen; ACB, nucleus accumbens; GPi, globus pallidus, internal segment; SI, substantia innominate. Scale bars: 5 µm.

**Fig. S9. Related to Fig. 2.**
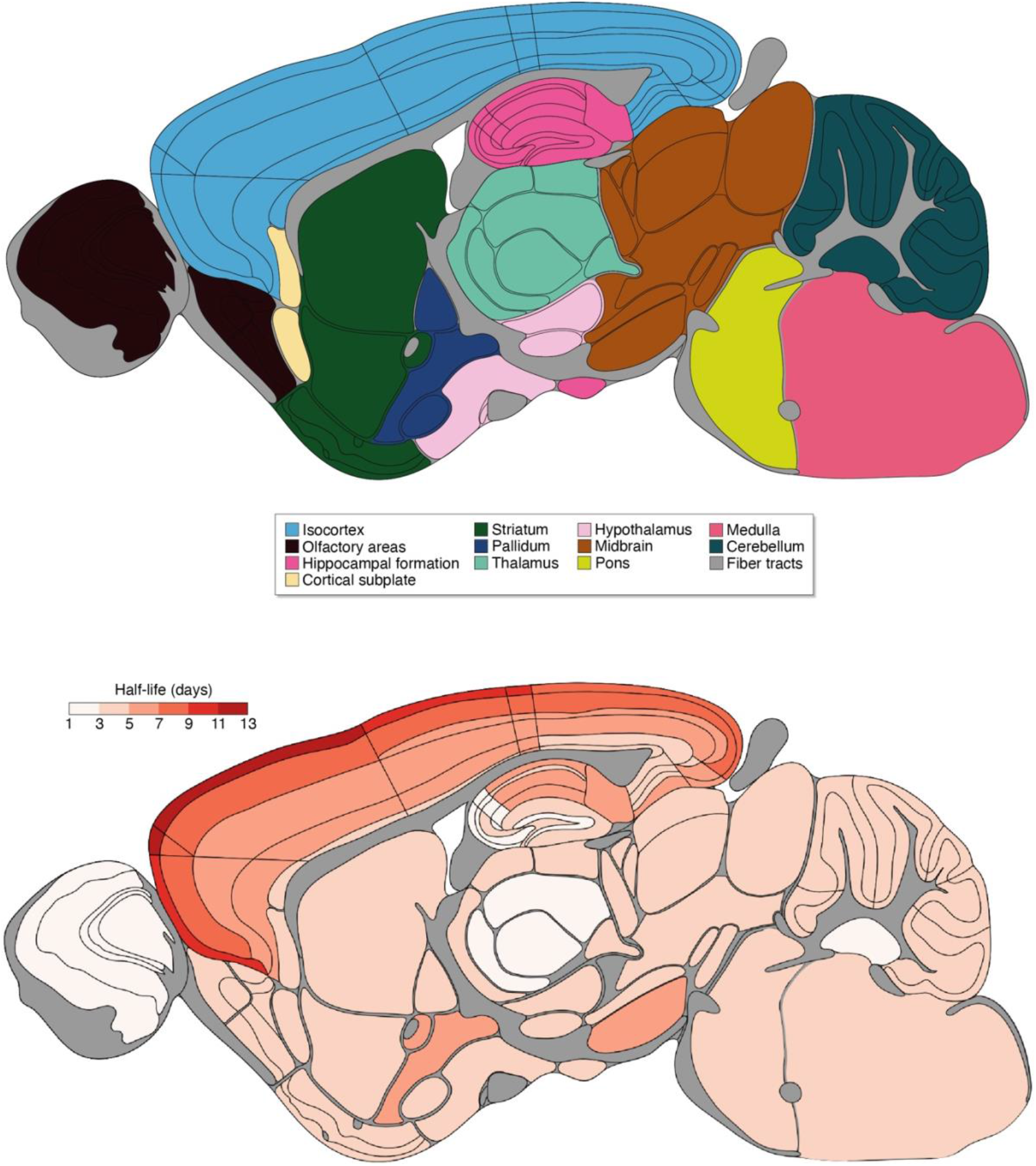
Regional and subregional diversity in PSD95 puncta density half-life in 3M mouse brain. PSD95 puncta lifetime (^PSD95 density^t_1/2_) across 110 subregions of the 3M mouse brain. The main brain regions from which these subregions derive are illustrated at the top. See Protein Lifetime Synaptome Atlas for ^PSD95 density^t_1/2_ values in each subregion (Bulovaite et al., 2021).

**Fig. S10. Related to Fig. 2.**
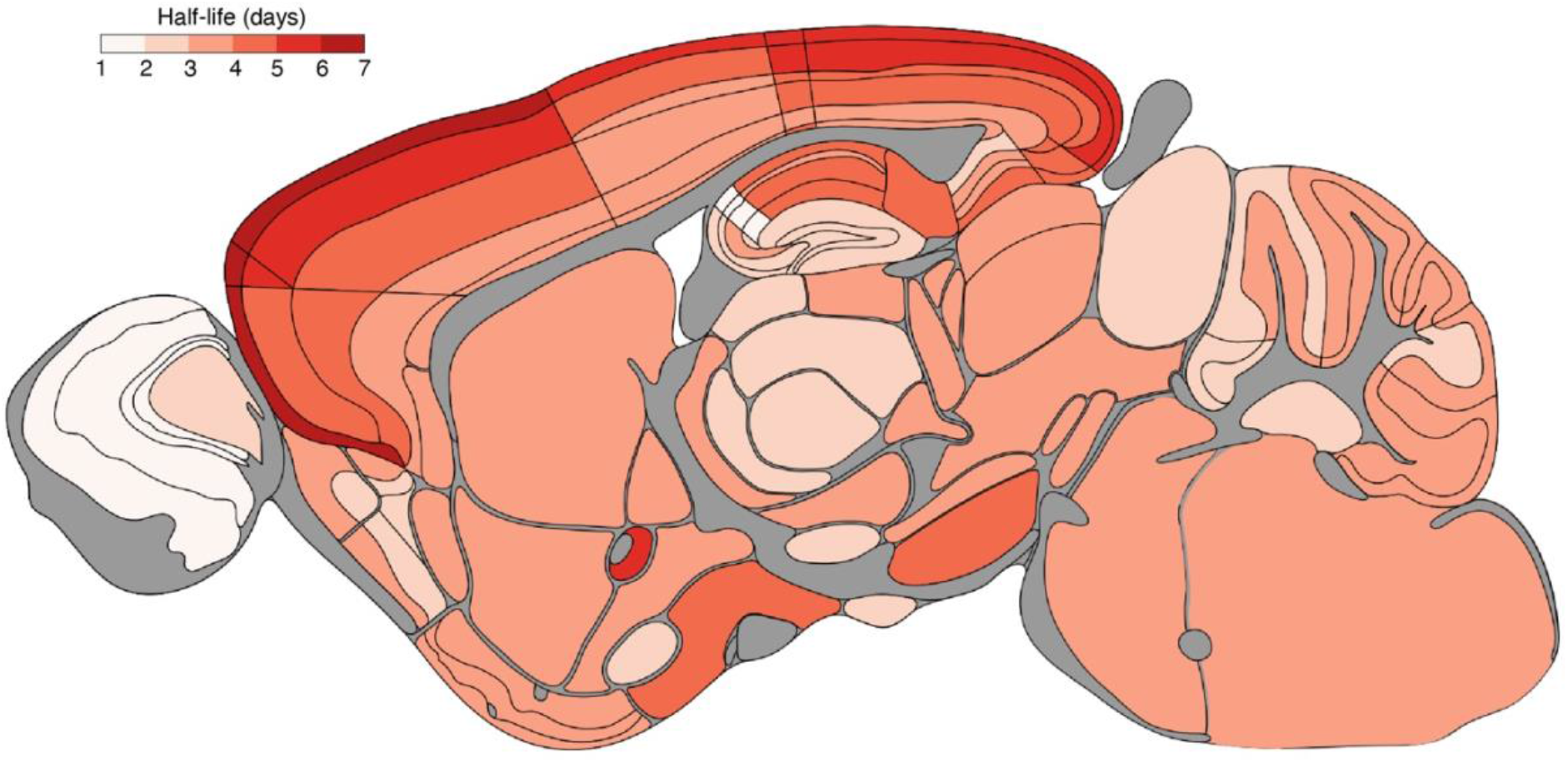
Regional and subregional diversity in synaptic PSD95 protein intensity half-life in 3M mouse brain. Synaptic PSD95 protein lifetime (^PSD95 intensity^t_1/2_) mapped for 110 mouse brain regions. See Protein Lifetime Synaptome Atlas for ^PSD95 intensity^t_1/2_ values in each subregion (Bulovaite et al., 2021).

**Fig. S11. Related to Fig. 2.**
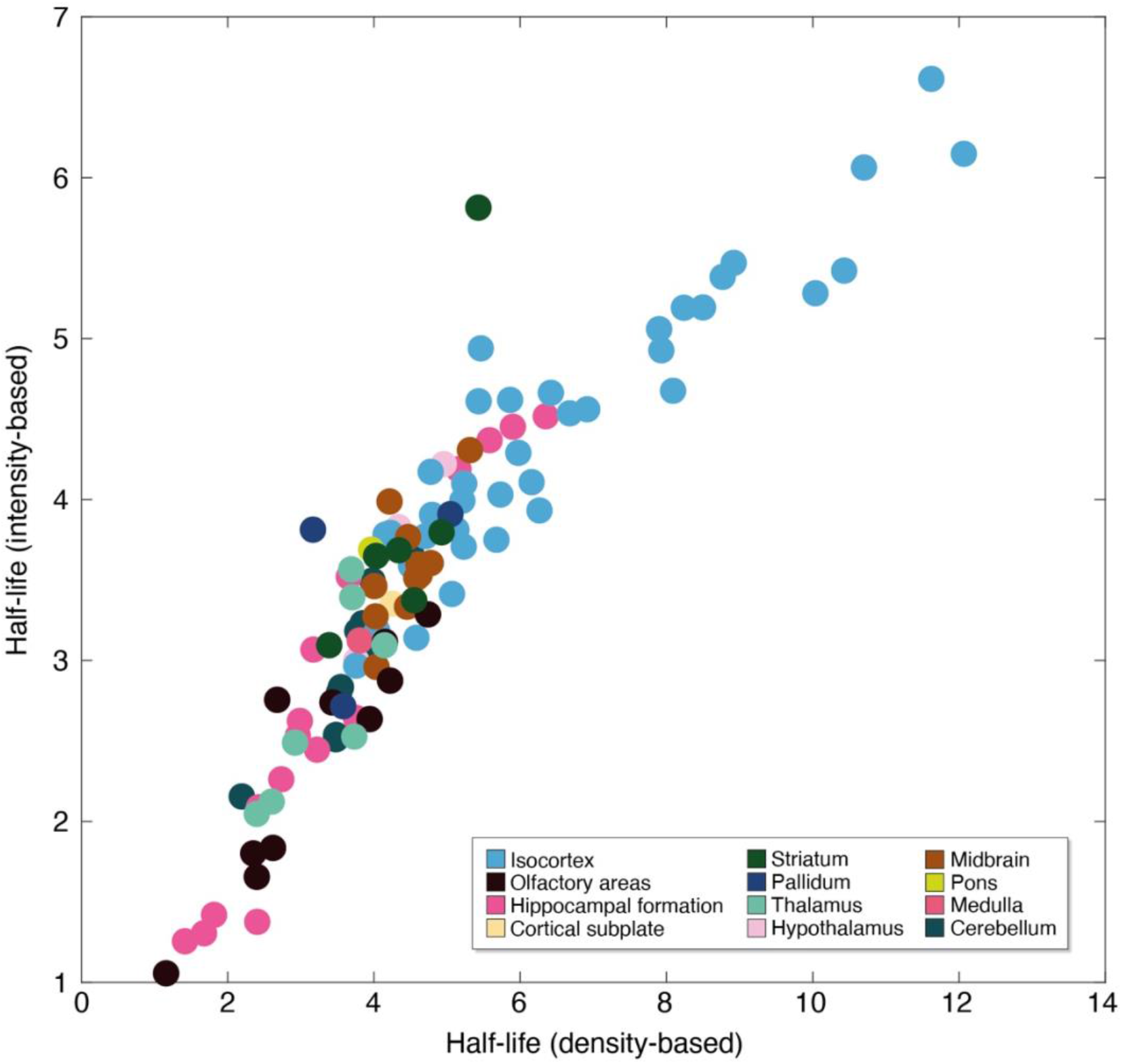
Correlation between density-based and intensity-based half-life estimates. Density-based half-life (x-axis) was plotted against intensity-based half-life (y-axis) for all brain subregions examined. Each dot represents a subregion, color-coded according to the main brain region that they belong to. Half-life is in days. R = 0.9088 and P < 0.0001, Pearson’s correlation test.

**Fig. S12. Related to Fig. 2.**
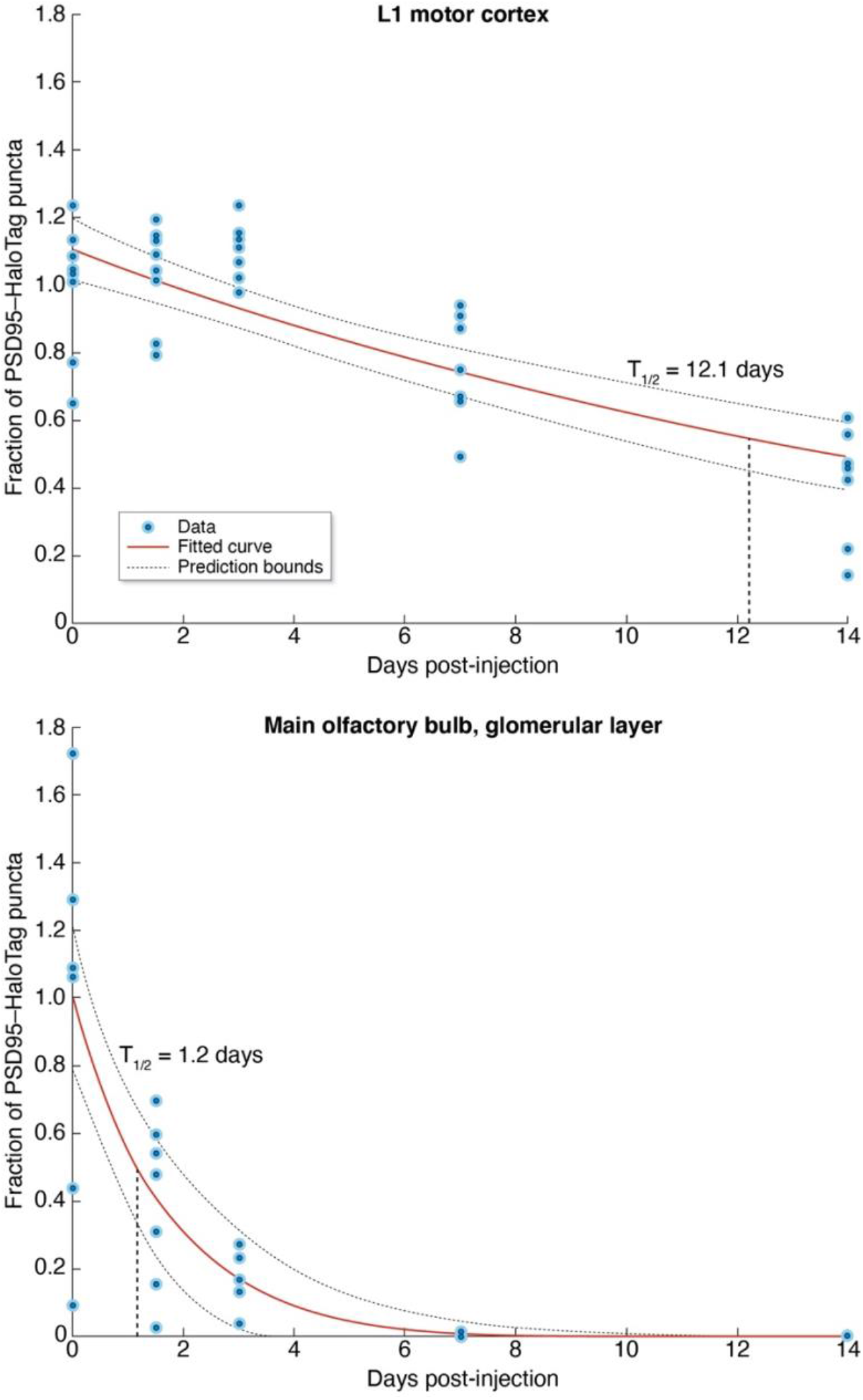
Brain regions with characteristically long and short PSD95 punctum half-lives. Example data-fitted exponential decay curves for regions with especially long-lived (top) and short-lived (bottom) PSD95 puncta (mean ± 95% confidence interval). Dots represent data from individual animals. Data from days 1.5, 3, 7 and 14 are normalized to the mean of day 0 and the fraction of puncta remaining is plotted on the y-axis. Dashed line indicates half-life.

**Fig. S13. Related to Fig. 3.**
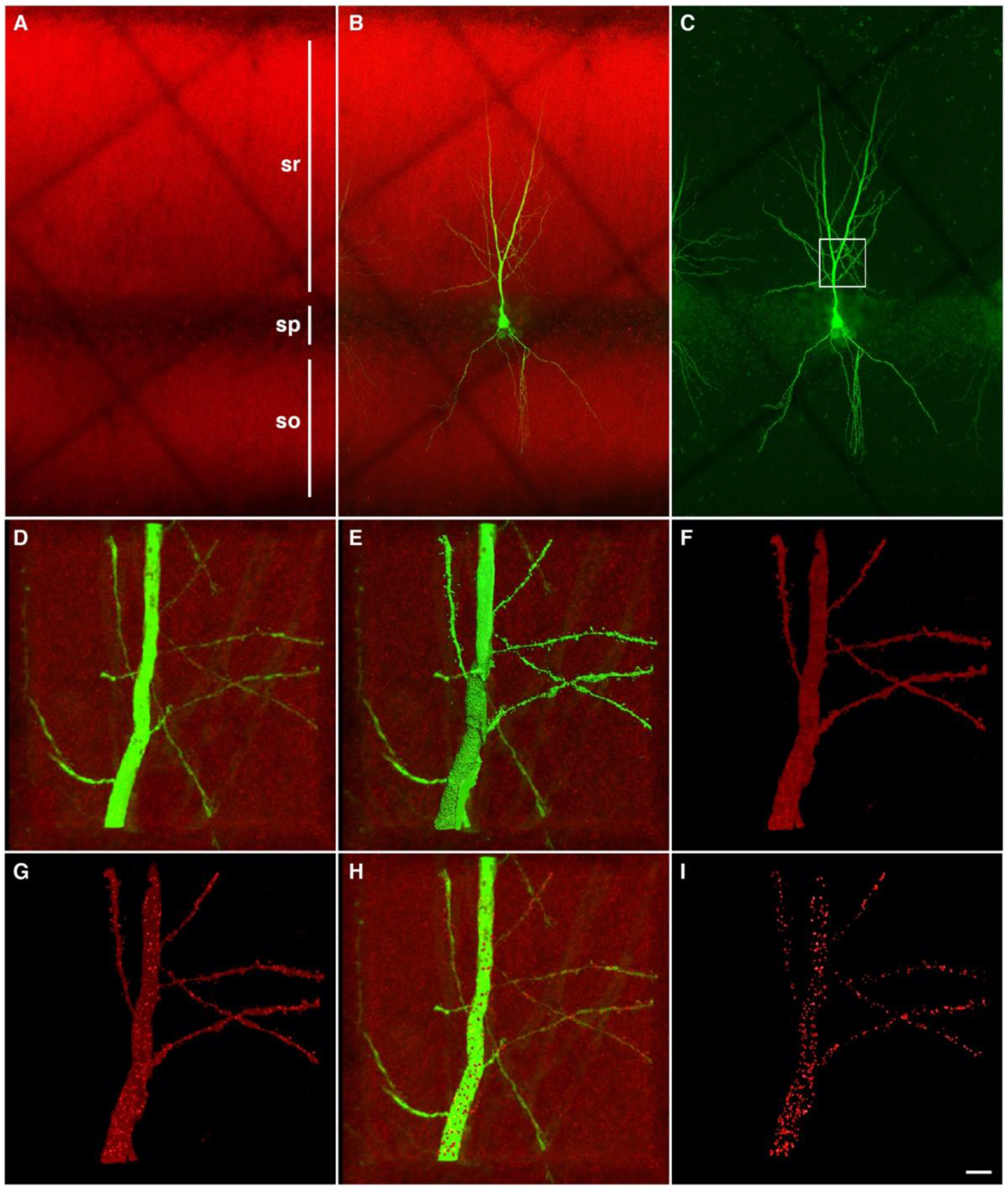
Intracellular injections and SiR-Halo-positive puncta 3D reconstruction. (**A-C**) Confocal microscopy images showing (A) SiR-Halo signal (red) and (B) a CA1 pyramidal neuron intracellularly injected with Alexa 488 (green). (C) Same neuron as in (B) with overexposed green channel to show the total extension of the apical dendritic shaft. White box indicates the area of the analysis. (**D-I**) Representative confocal microscopy images showing the 3D reconstruction of an apical dendrite segment (D). A solid surface was created to match the apical dendritic shaft (E, green). The red signal inside of the solid surface is selected (F) and different intensity threshold surfaces that match the SiR-Halo-positive puncta were created (G, red). Then, the solid surfaces that exactly matched the SiR-Halo-positive puncta were selected for each image stack (H). In (I), only the solid surfaces for the SiR-Halo-positive puncta are shown. Sr, stratum radiatium; sp, stratum pyramidale; so, stratum oriens. Scale bar: 20 µm (A-C), 7 µm (D-I).

**Fig. S14. Related to Fig. 3.**
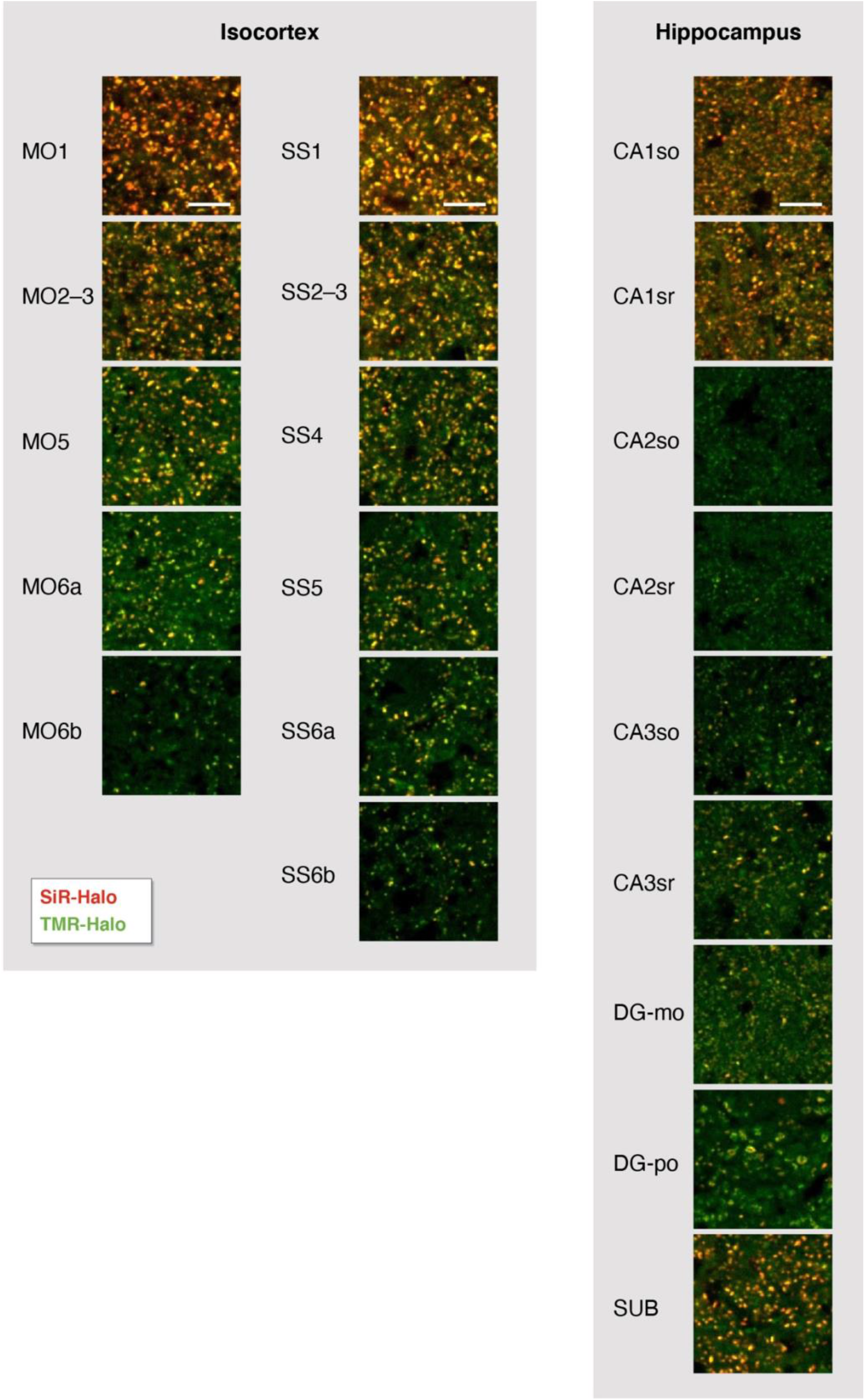
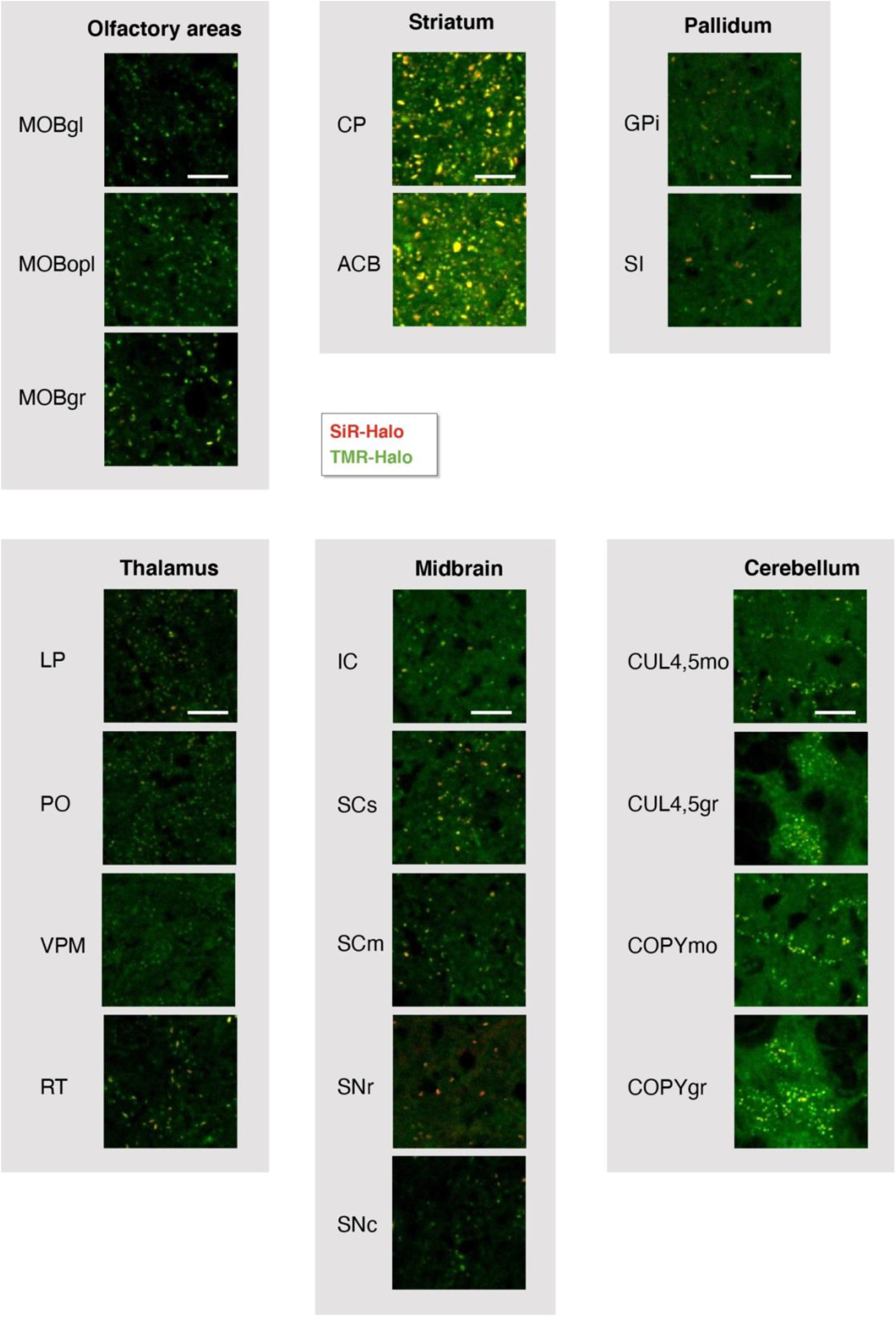
Colocalization of SiR-Halo and TMR-Halo labeling. Representative high-magnification images of SiR-Halo and TMR-Halo fluorescence labeling at day 3 post-injection of SiR-Halo. Examples presented cover subregions of isocortex, hippocampal formation, olfactory bulb, striatum, pallidum, thalamus, midbrain and cerebellum. CA1so, CA1 stratum oriens; CA1sr, CA1 stratum radiatum; CA2so, CA2 stratum oriens; CA2sr, CA2 stratum radiatum; CA3so, CA3 stratum oriens; CA3sr, CA3 stratum radiatum; DGmo, dentate gyrus molecular layer; DGpo, dentate gyrus polymorphic layer; SUB, subiculum; MOBgr, granular layer of the main olfactory bulb; CP, caudoputamen; ACB, nucleus accumbens; GPi, globus pallidus, internal segment; SI, substantia innominate; LP, lateral posterior nucleus of the thalamus; PO, posterior complex of the thalamus; VPM, ventral posteromedial nucleus of the thalamus; RT, reticular nucleus of the thalamus; IC, inferior colliculus; SCs, superior colliculus, sensory related; SCm, superior colliculus, motor related; SNr, substantia nigra, reticular part; SNc, substantia nigra, compact part; CUL4,5mo, lobules IV-V molecular layer; CUL4,5gr, lobules IV-V granular layer; COPYmo, copula pyramidis molecular layer; COPYgr, copula pyramidis granular layer. Scale bars: 5 µm.

**Fig. S15. Related to Fig. 3.**
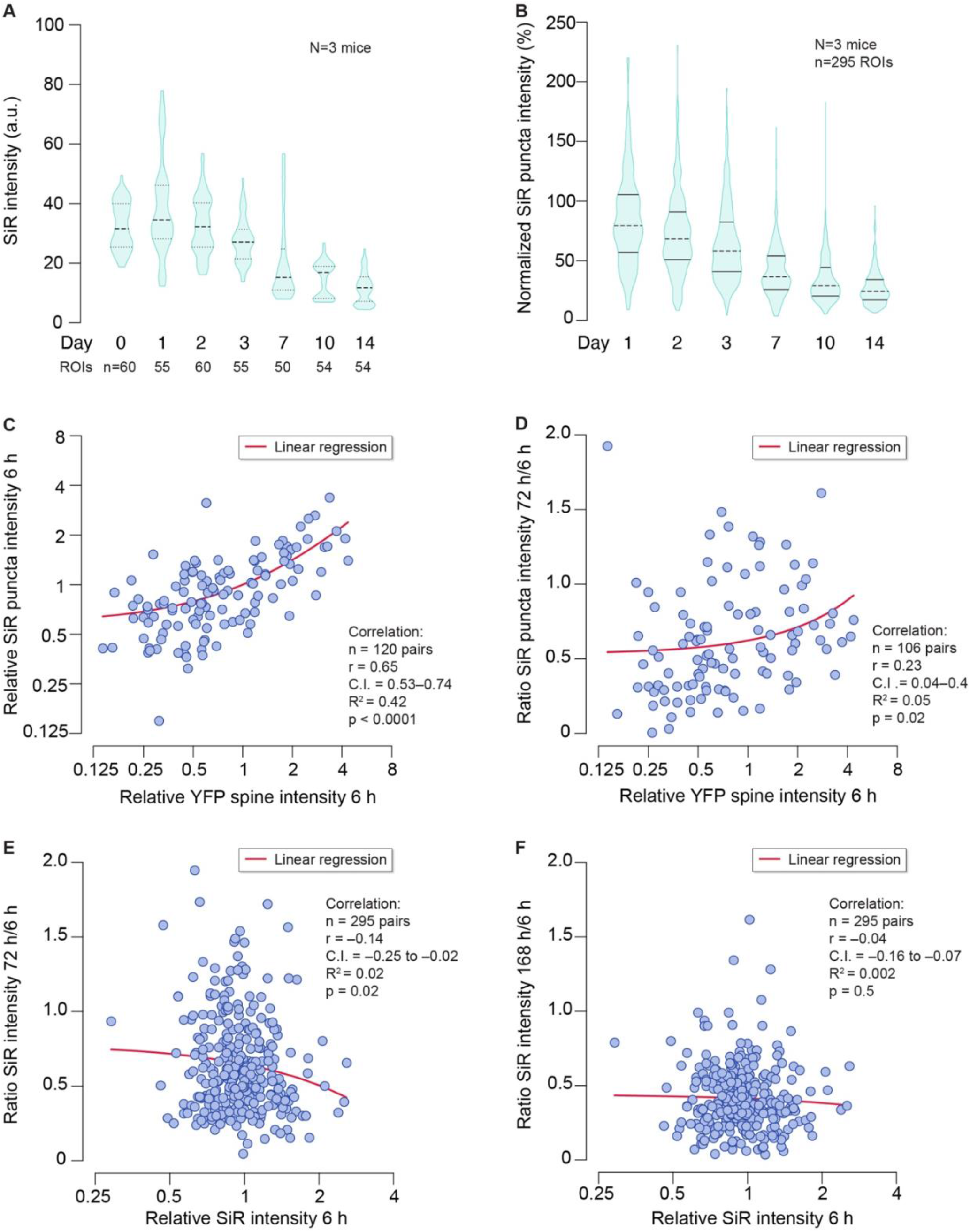
In vivo imaging of dendritic spine and PSD95 turnover. (**A**) Violin plots of SiR ligand intensity (arbitrary units) in 11 μm x 11 μm regions of interest (ROIs) at 0, 1, 2, 3, 7, 10 and 14 days post injection. Each ROI is imaged only once over the entire time course. (**B**) Violin plots of SiR-ligand puncta intensities, normalized to the first imaging time point (day 0). Each punctum is tracked over time. Data correspond to Figure 3F. (**C**) Relative SiR ligand puncta intensities versus relative parent dendritic spine YFP intensities at the first imaging time point (day 0). Each data point represents the intensities relative to the mean puncta and spine intensity in its parent field of view. Line, linear regression, indicating a positive correlation between spine intensity (size) and SiR puncta intensity. (**D**) SiR ligand puncta fluorescence intensity at day 3 post injection as a fraction of their intensity at day 0, versus the relative parent dendritic spine YFP intensity. Line, linear regression, indicating a very weak correlation between the spine intensity (size) and PSD95 retention. (**E**) SiR ligand puncta fluorescence intensity at day 3 post injection as a fraction of their intensity at day 0, versus the relative SiR ligand puncta intensities at day 0. Line, linear regression, indicating a very weak negative correlation between the initial PSD95 content and its retention. (**F**) As (E), but for the day 7 time point. No significant correlation.

**Fig. S16. Related to Fig. 4.**
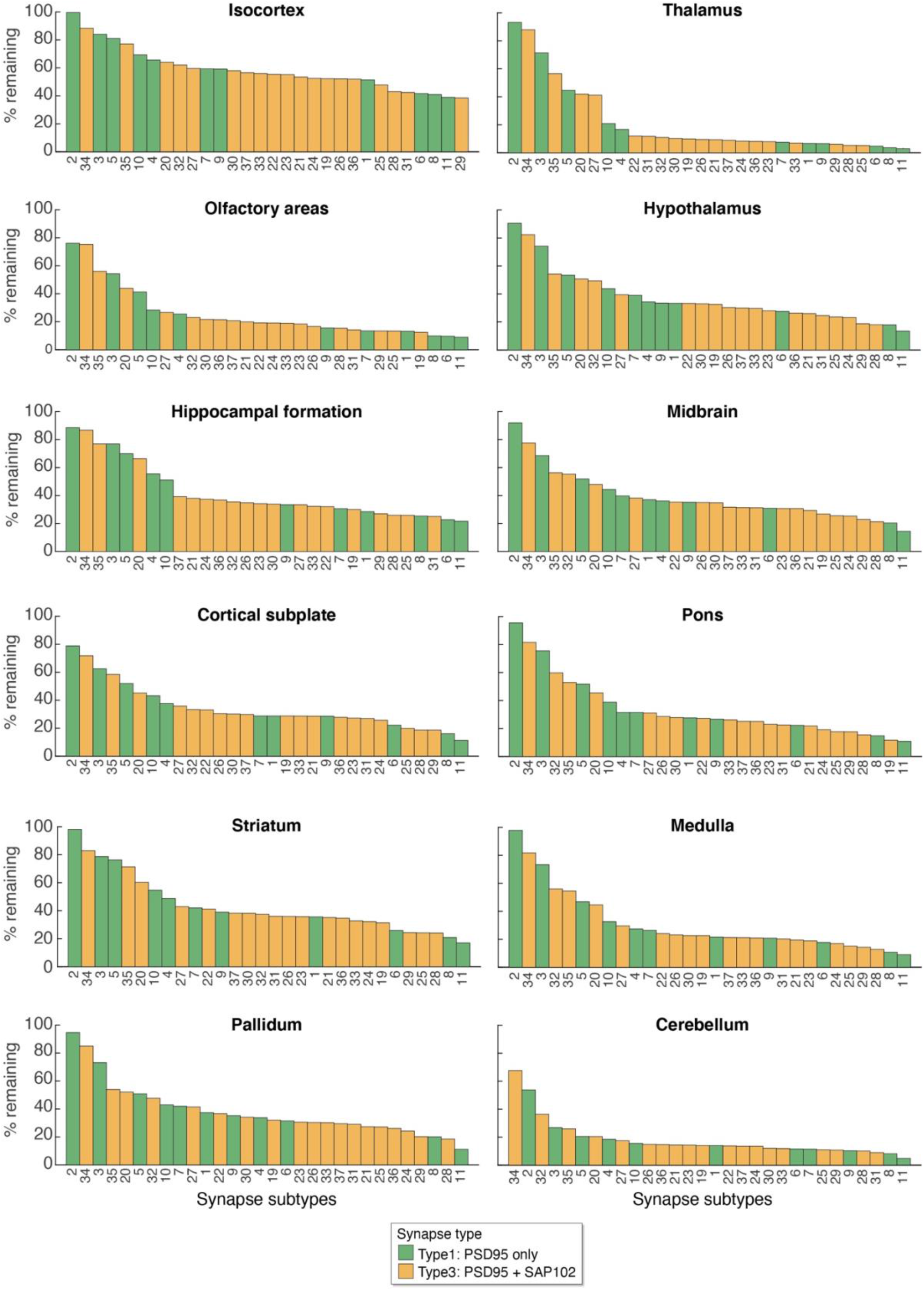
Synapse subtype lifetimes across brain regions. Synapse subtypes are ranked according to their PSD95 protein lifetime in each of the 12 main brain regions. Bar height indicates the percentage of SiR-Halo-positive subtype density remaining at day 7 compared with day 0. Of the 37 synapse subtypes originally defined (Zhu et al., 2018), the 30 studied here are color-coded by their type 1 (PSD95 only) or type 3 (PSD95+SAP102) molecular composition.

**Fig. S17. Related to Fig. 4.**
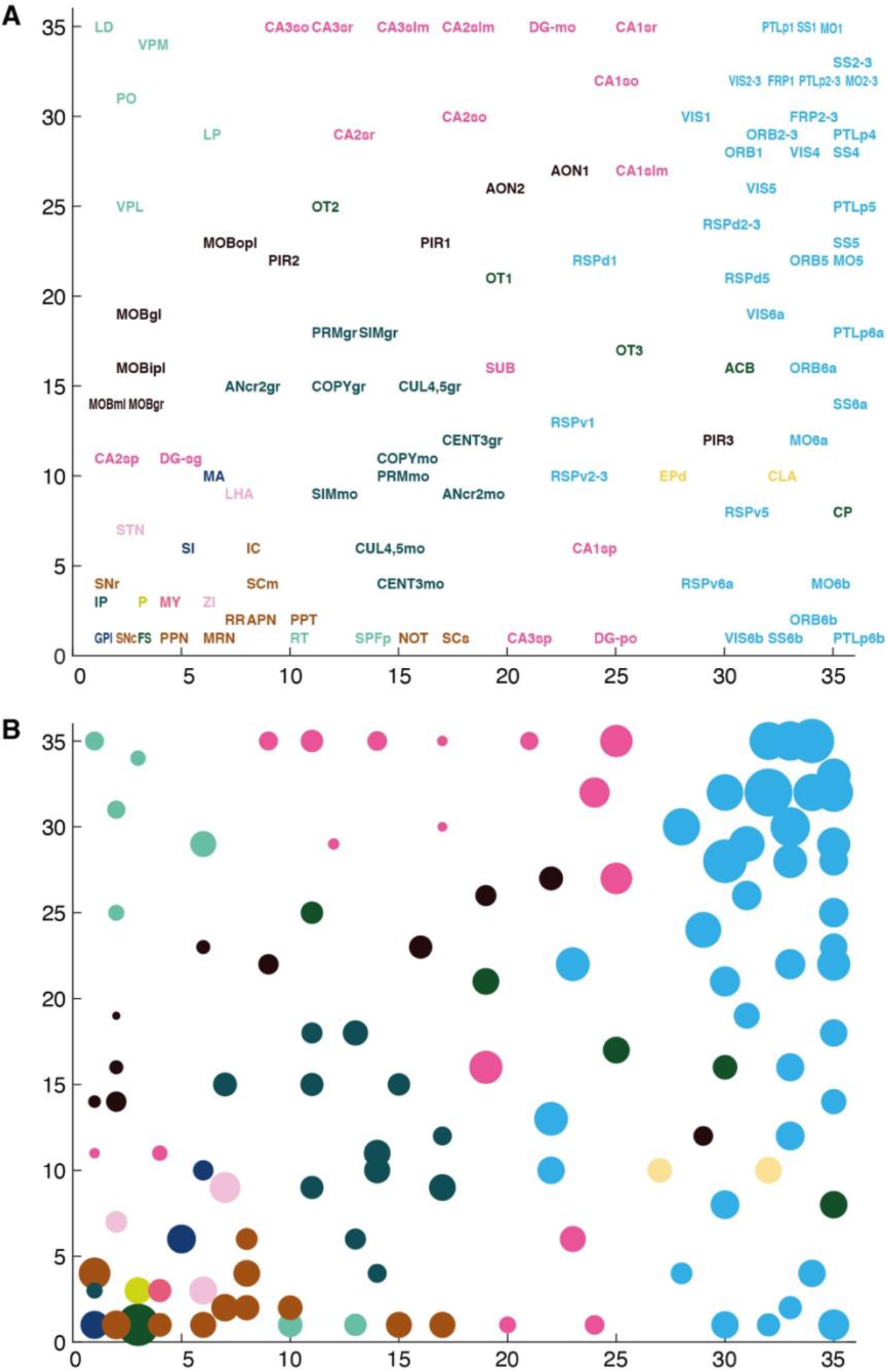
Self-organizing feature maps of synapse subtypes show systematic anatomical organization. Self-organizing feature map (SOM) clustering based on subtype densities. Each region is described by its 37 densities of the subtypes and a resulting 2-dimensional SOM clustering is shown. (**A**) Subregion position is indicated by the lower left corner of the first letter in the subregion name. Subregion names are color-coded according to region (see key in Figure 2A). Axes indicate node index number in the 35×35 SOM node network. (**B**) Same network as in (A) but each subregion is represented by a disc, the center of which corresponds to the winning SOM node position and the radius codes for the half-life (^PSD95 intensity^t_1/2_).

**Fig. S18. Related to Fig. 4.**
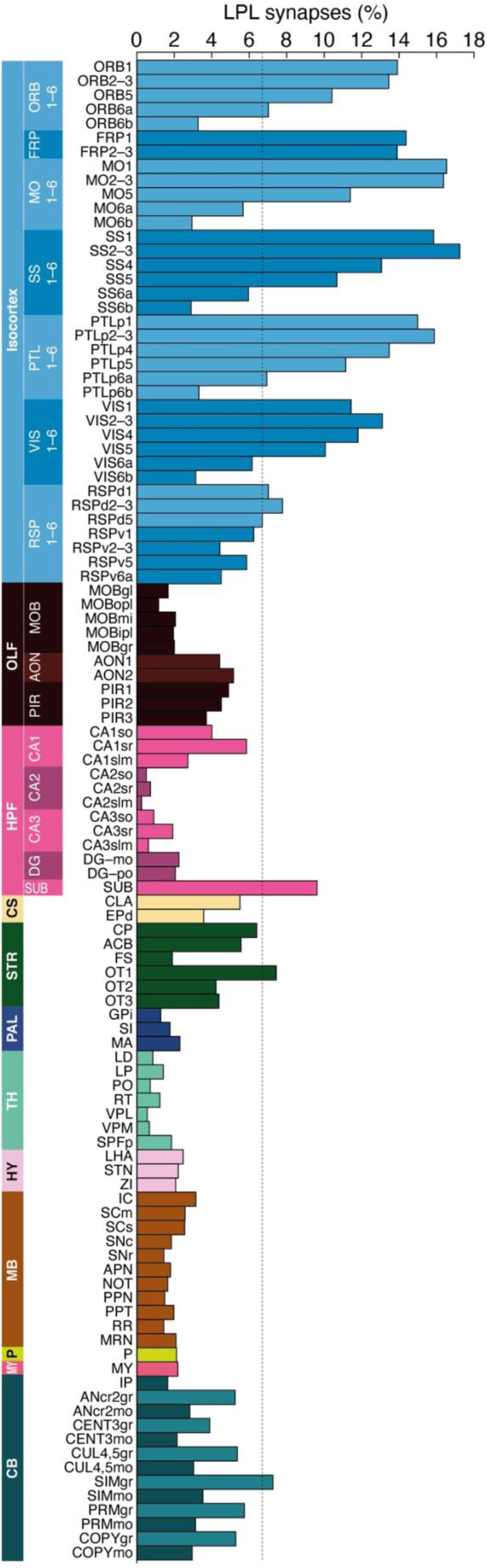
LPL synapse composition of brain subregions. The percentage of LPL synapses in brain regions and subregions. Superficial layers of the isocortex (numbered) contain the most LPL synapses. Dotted line indicates average for whole brain (6.7%). Brain subregions are listed in Table S1.

**Fig. S19. Related to Fig. 5.**
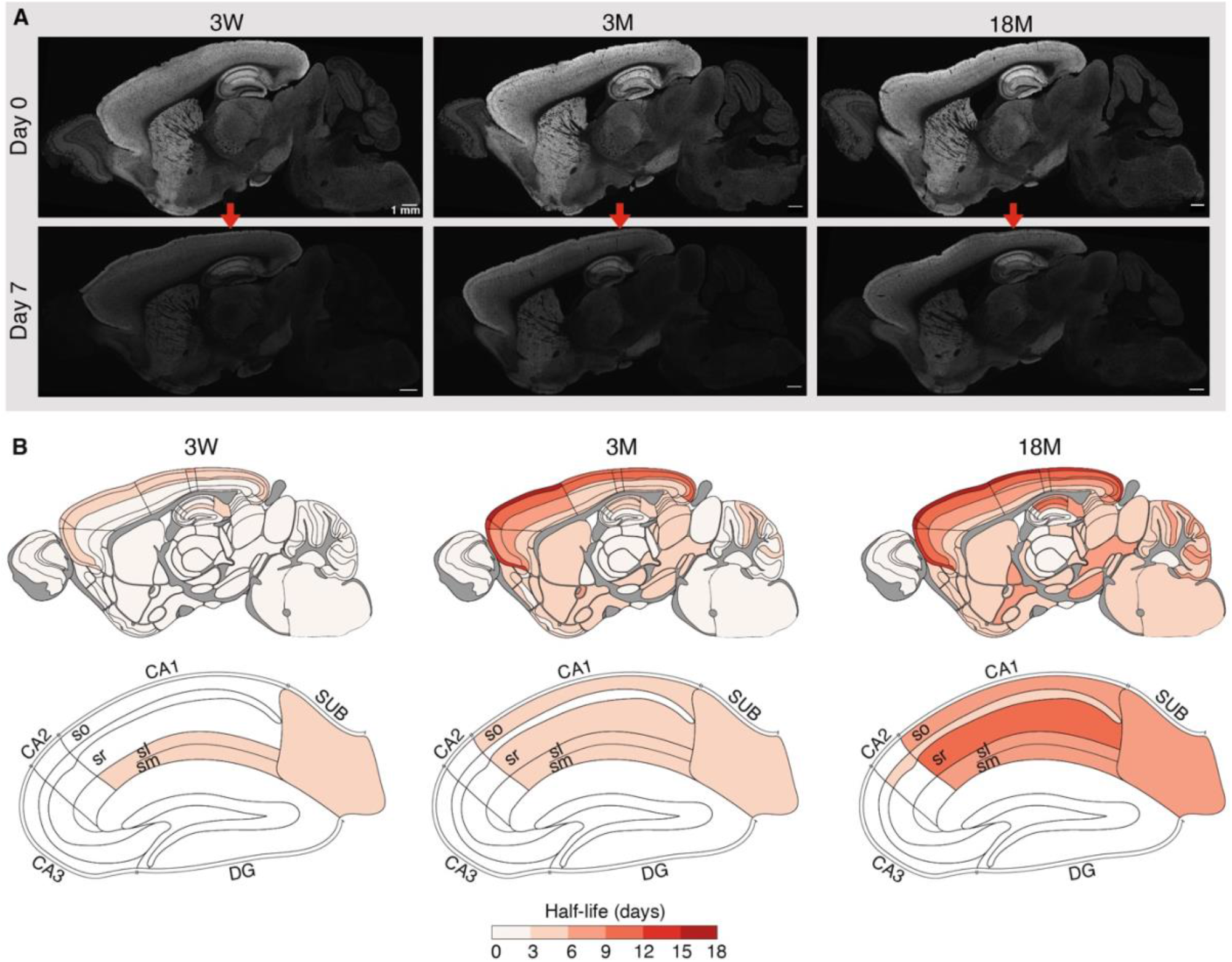
Synapse protein lifetime increases across the lifespan. (**A**) Example images of sagittal sections of PSD95-HaloTag mice injected with SiR-Halo at 3 weeks (3W), 3 months (3M) and 18 months (18M) of age and imaged at day 0 and day 7 post injection. Scale bars: 1 mm. (**B**) Maps of PSD95 puncta half-life (^PSD95 density^t_1/2_) at 3W, 3M and 18M across 110 subregions of the whole mouse brain (top row) and the hippocampal formation (bottom row). See Protein Lifetime Synaptome Atlas for half-life values in each subregion (Bulovaite et al., 2021).

**Fig. S20. Related to Fig. 5.**
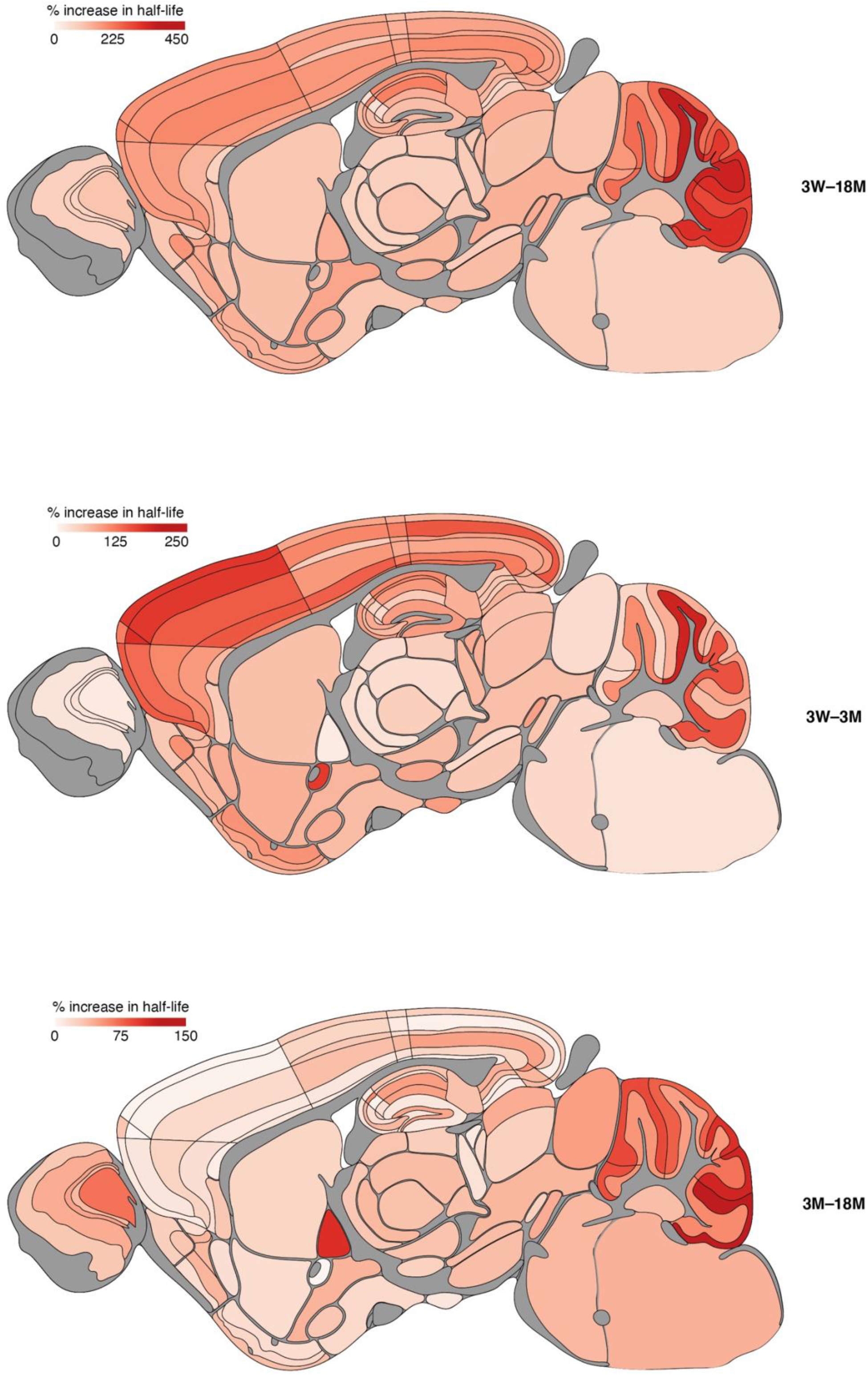
Increases in PSD95 puncta density lifetime between 3W, 3M and 18M. Percentage increases in PSD95 puncta half-life (^PSD95 density^t_1/2_) across 110 brain subregions between the indicated ages. See Protein Lifetime Synaptome Atlas for half-life values in each subregion (Bulovaite et al., 2021).

**Fig. S21. Related to Fig. 5.**
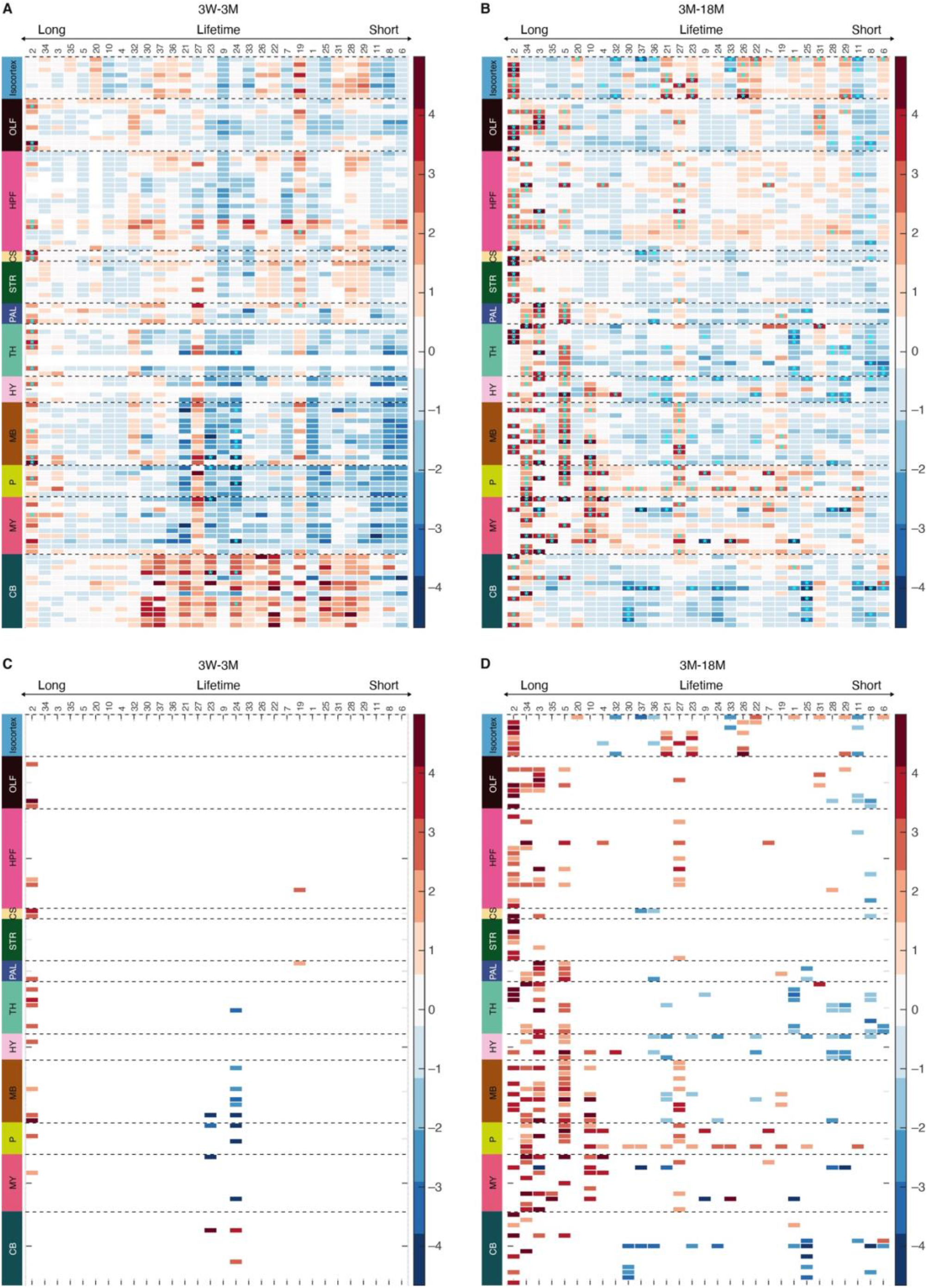
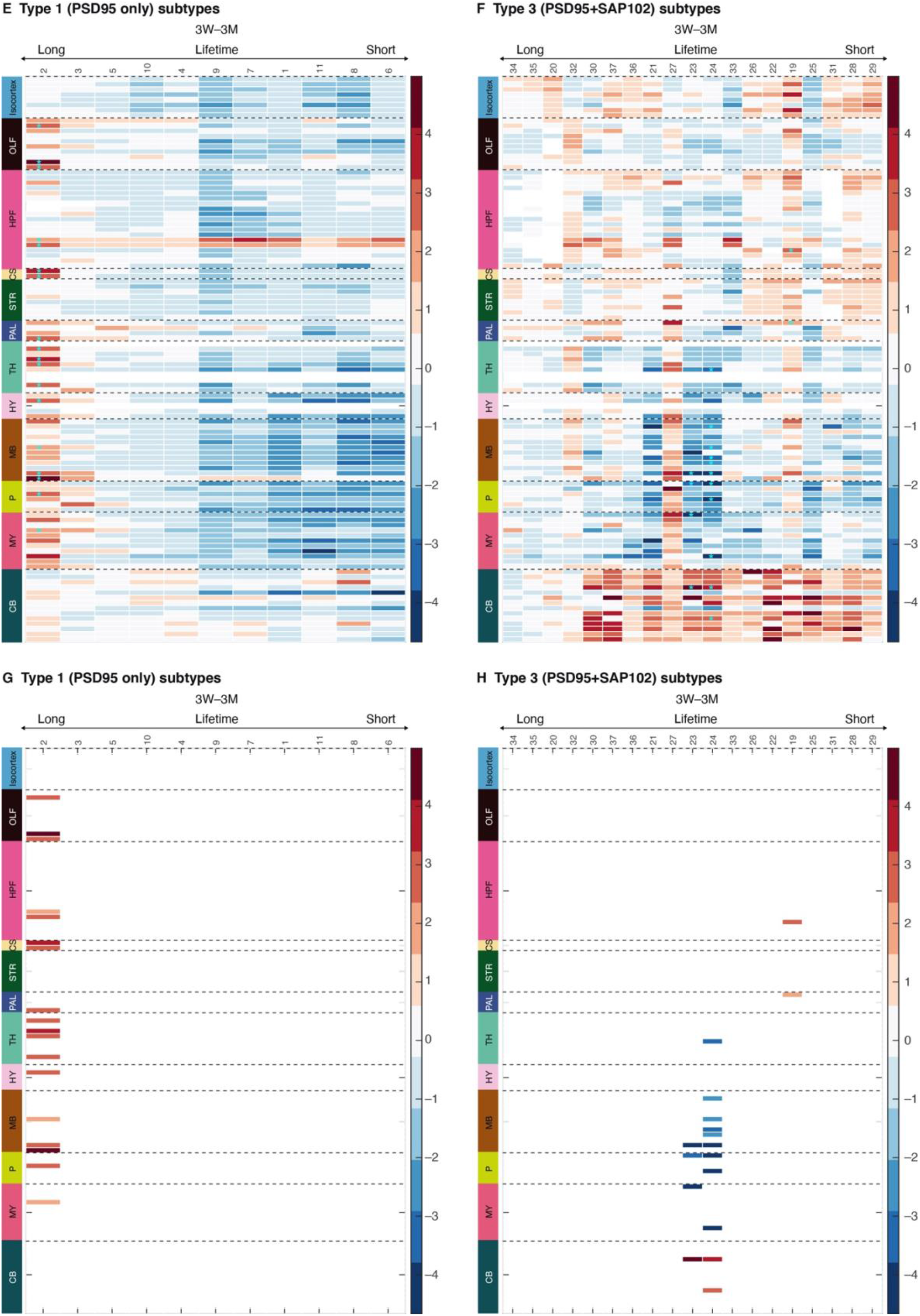
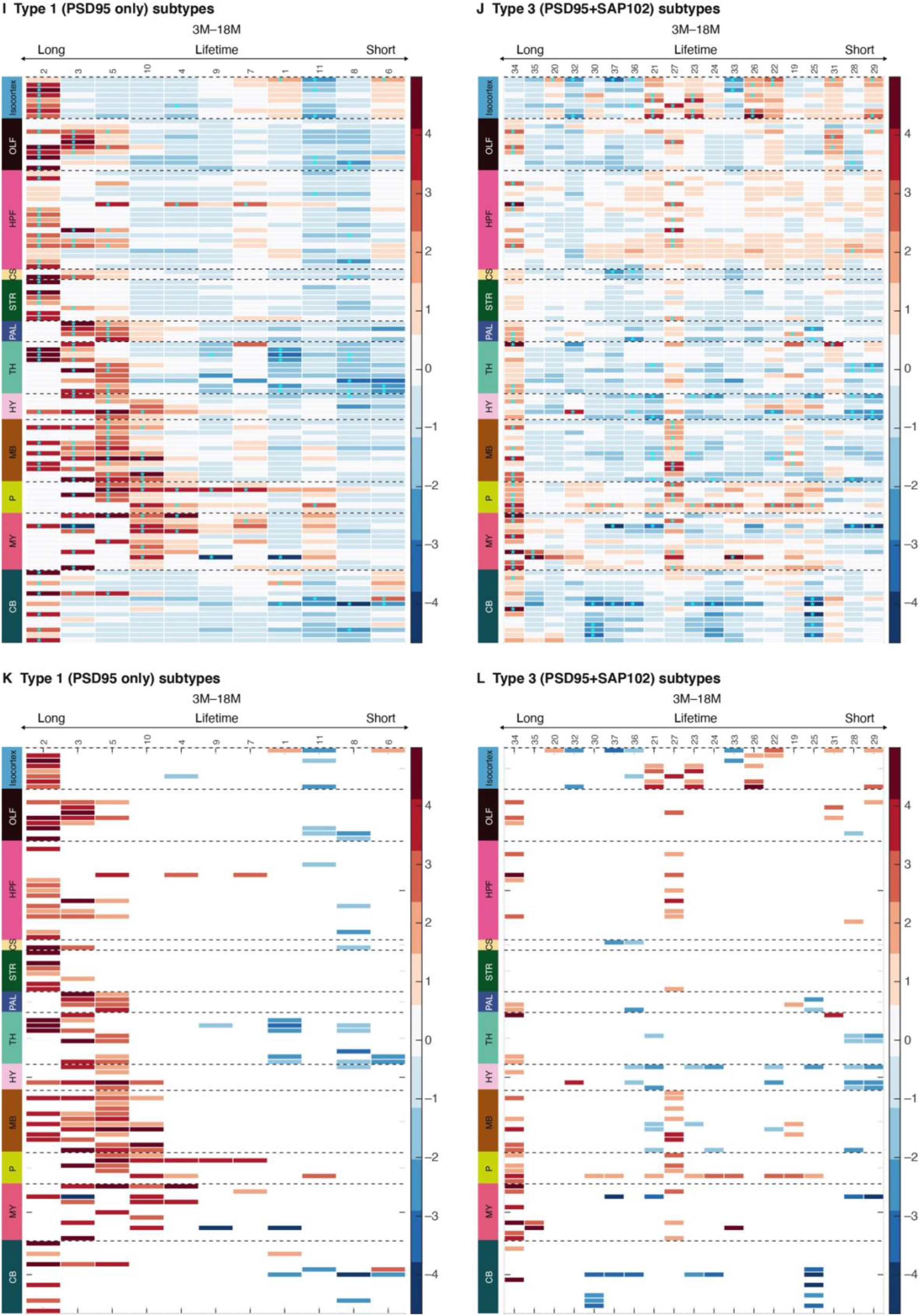
Changing lifespan architecture of synapse subtypes with different protein lifetimes. Heatmaps showing the change (Cohen’s *d* effect size) in synapse subtype (subtype number shown at top) density in 110 brain regions between 3W and 3M (A,C,E-H) and between 3M and 18M (B,D,I-L). Subtypes are ranked with longest lifetime on the left and shortest on the right. The 30 synapse subtypes are shown together (**A-D**) and separated into PSD95-only subtypes (**E**,**G**,**I**,**K**) and PSD95+SAP102 subtypes (**F**,**H**,**J**,**L**). Asterisks (A,B,E,F,I,J) indicate significant differences (P<0.05, Benjamini-Hochberg correction), with these shown alone beneath (C,D,G,H,K,L).

**Fig. S22. Related to Fig. 6.**
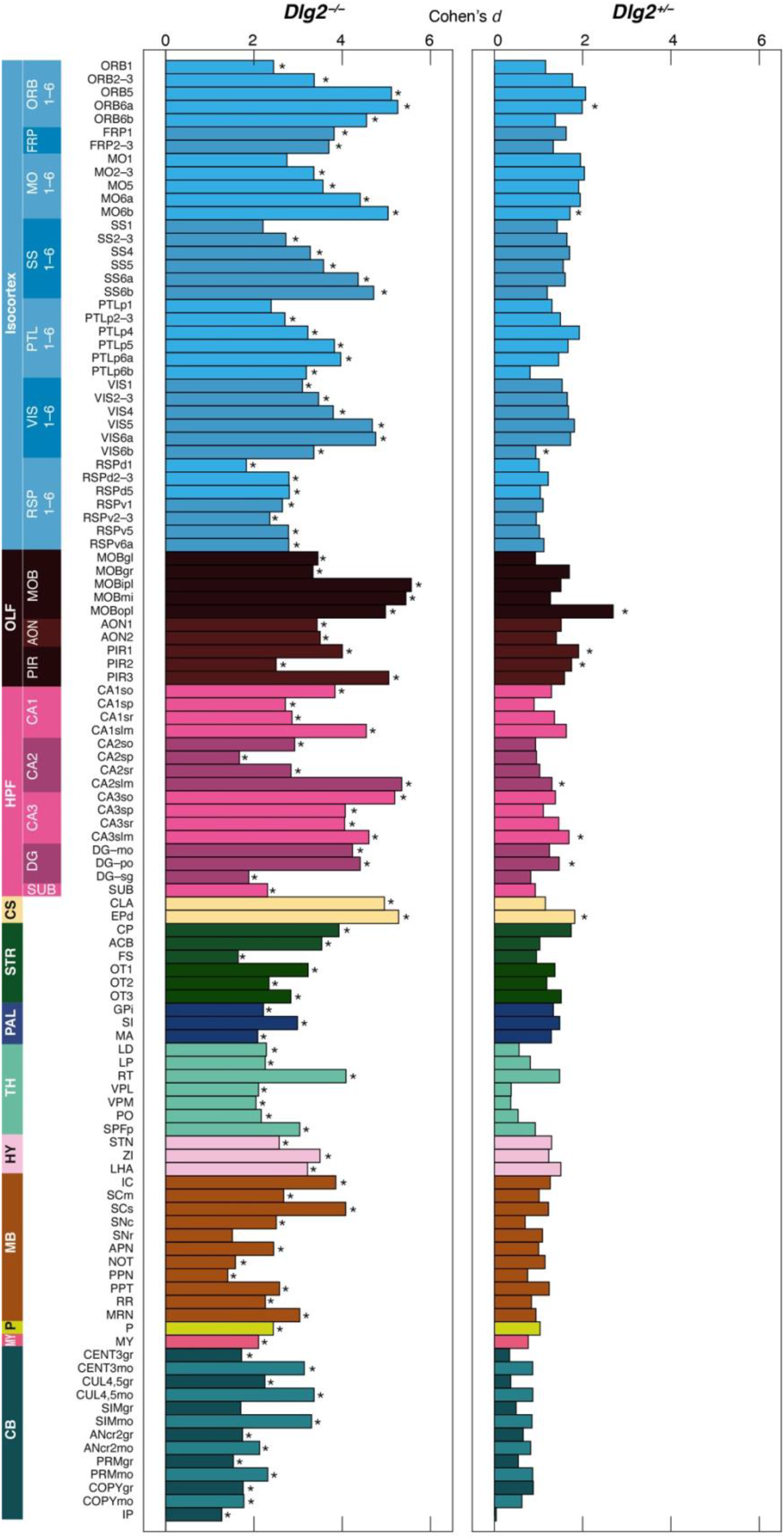
Synapse protein lifetime increases in *Dlg2* mutant mice. The increase (Cohen’s *d* effect size) in PSD95 puncta density half-life in 110 adult brain subregions in mice carrying a heterozygous (right) or homozygous (left) *Dlg2* mutation. Asterisks indicate significant differences from contol (*Dlg2*^+/+^) mice (P<0.05, Benjamini-Hochberg correction).

**Table S1.**
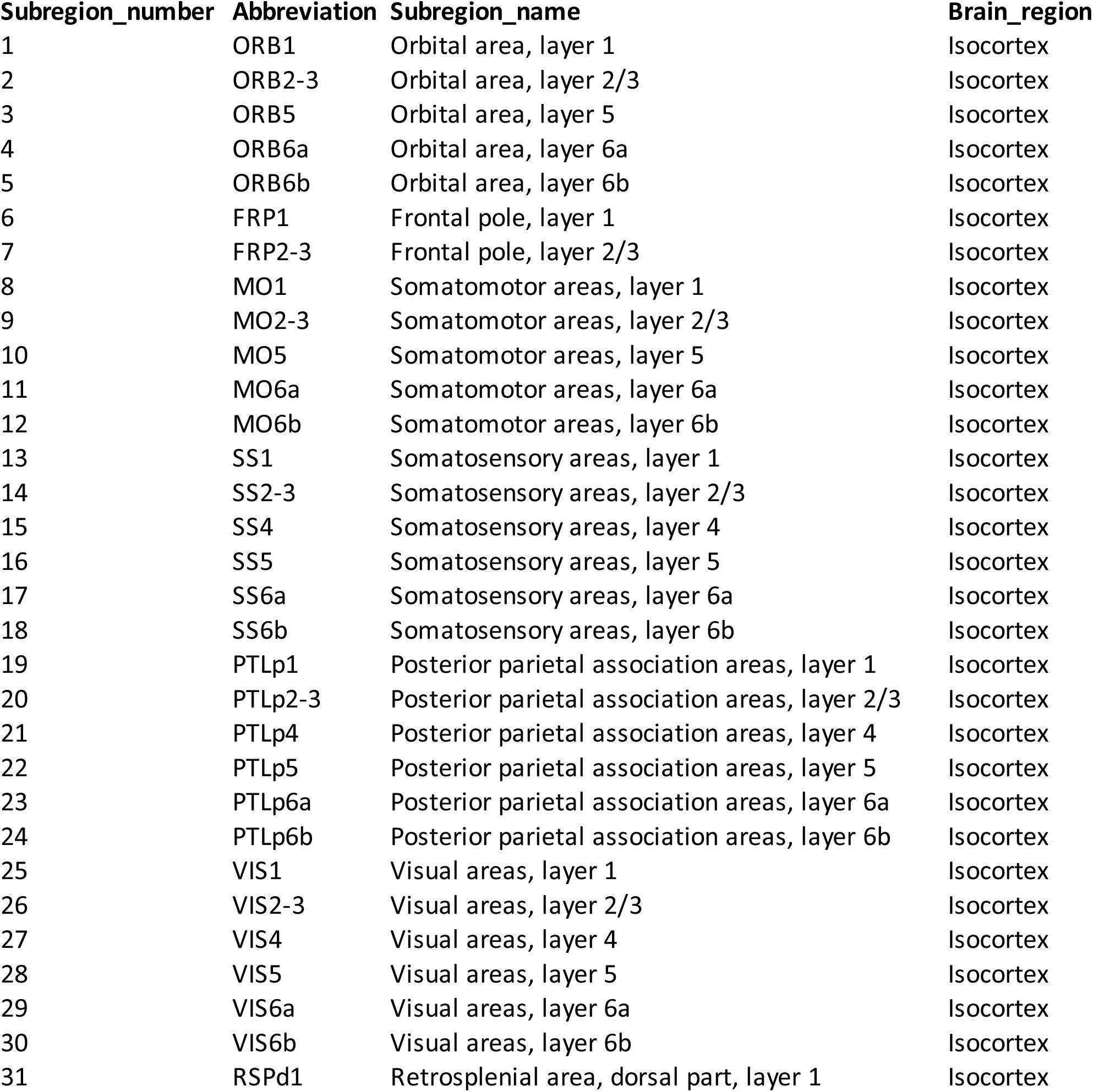
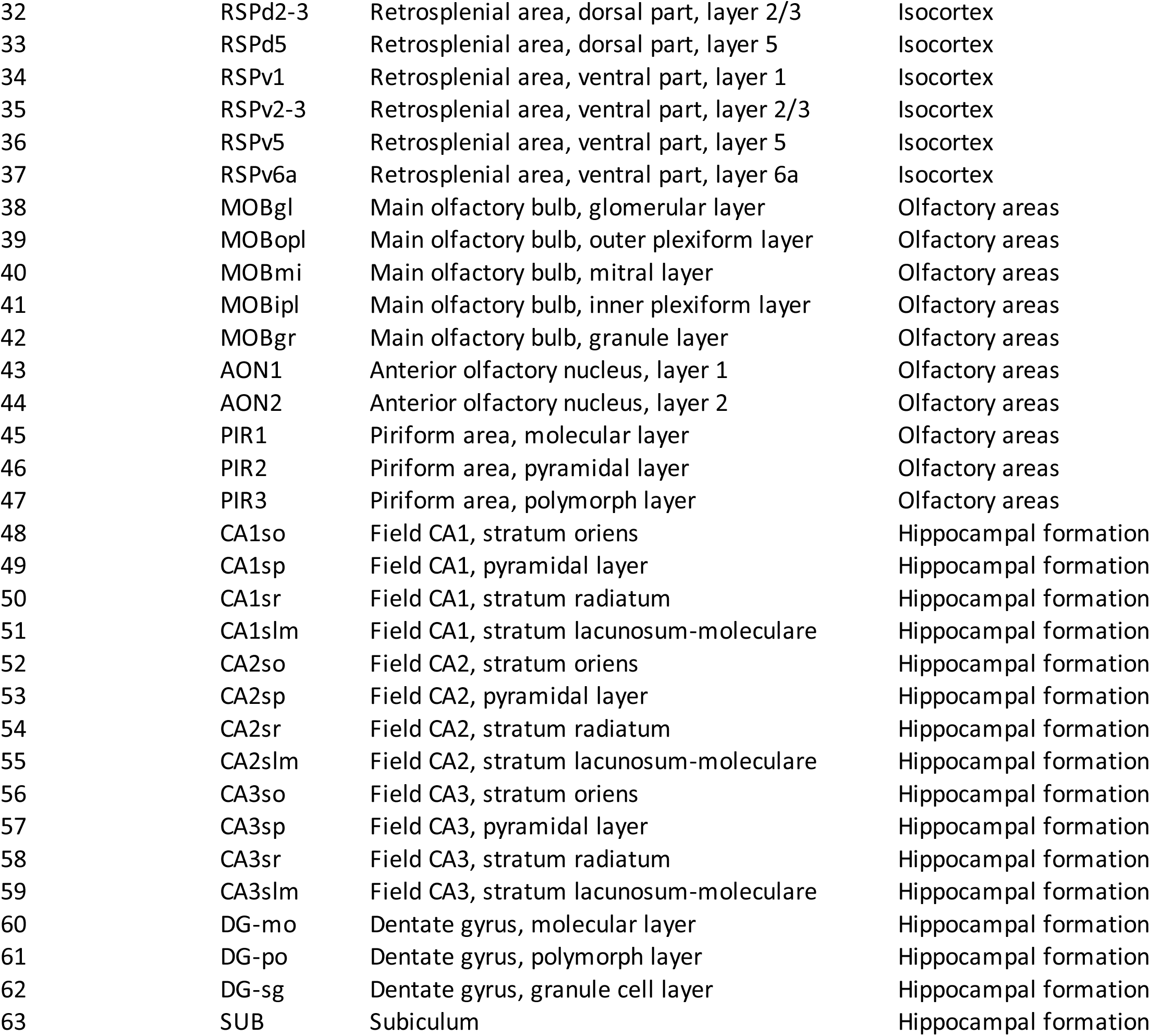
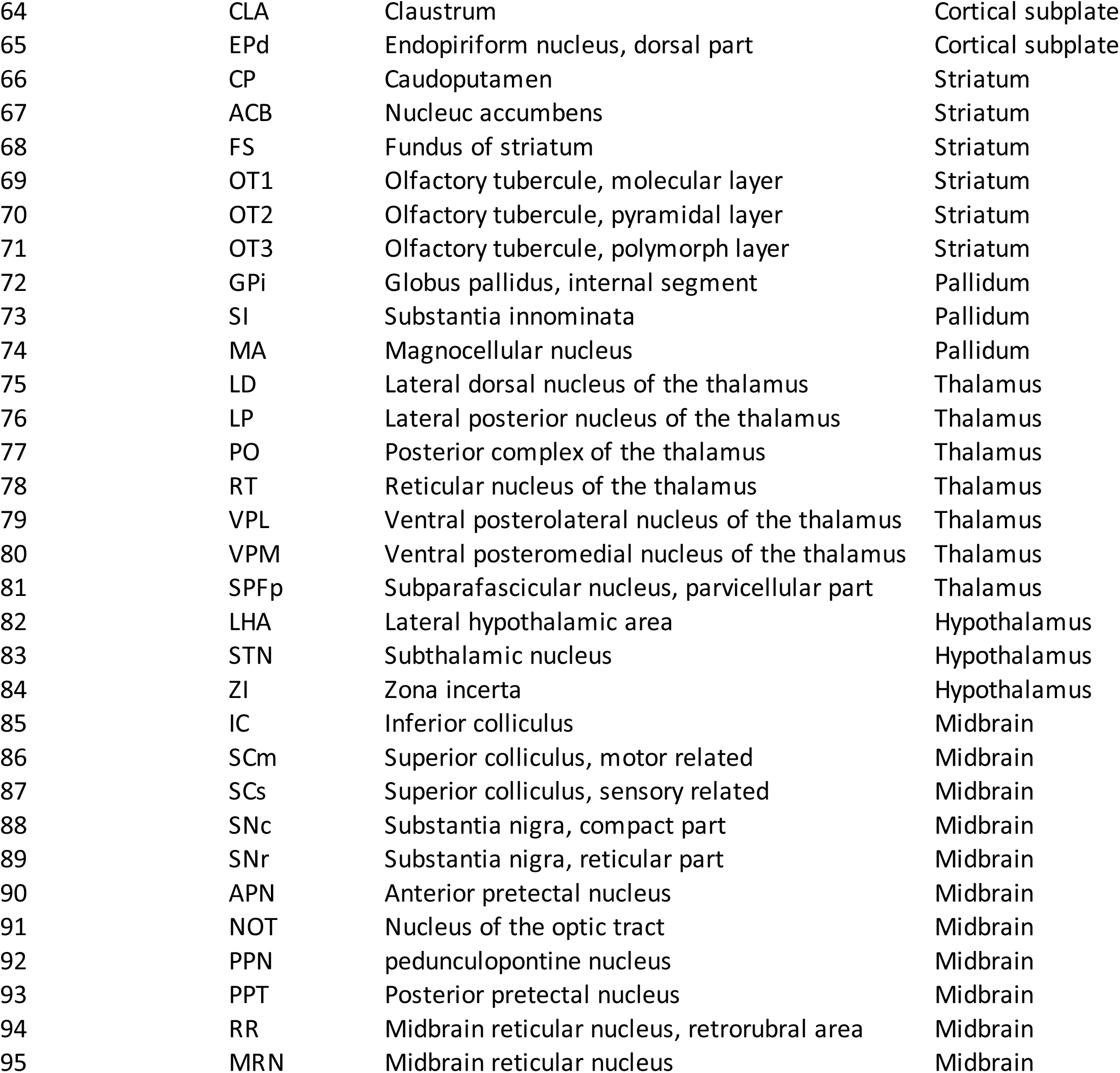
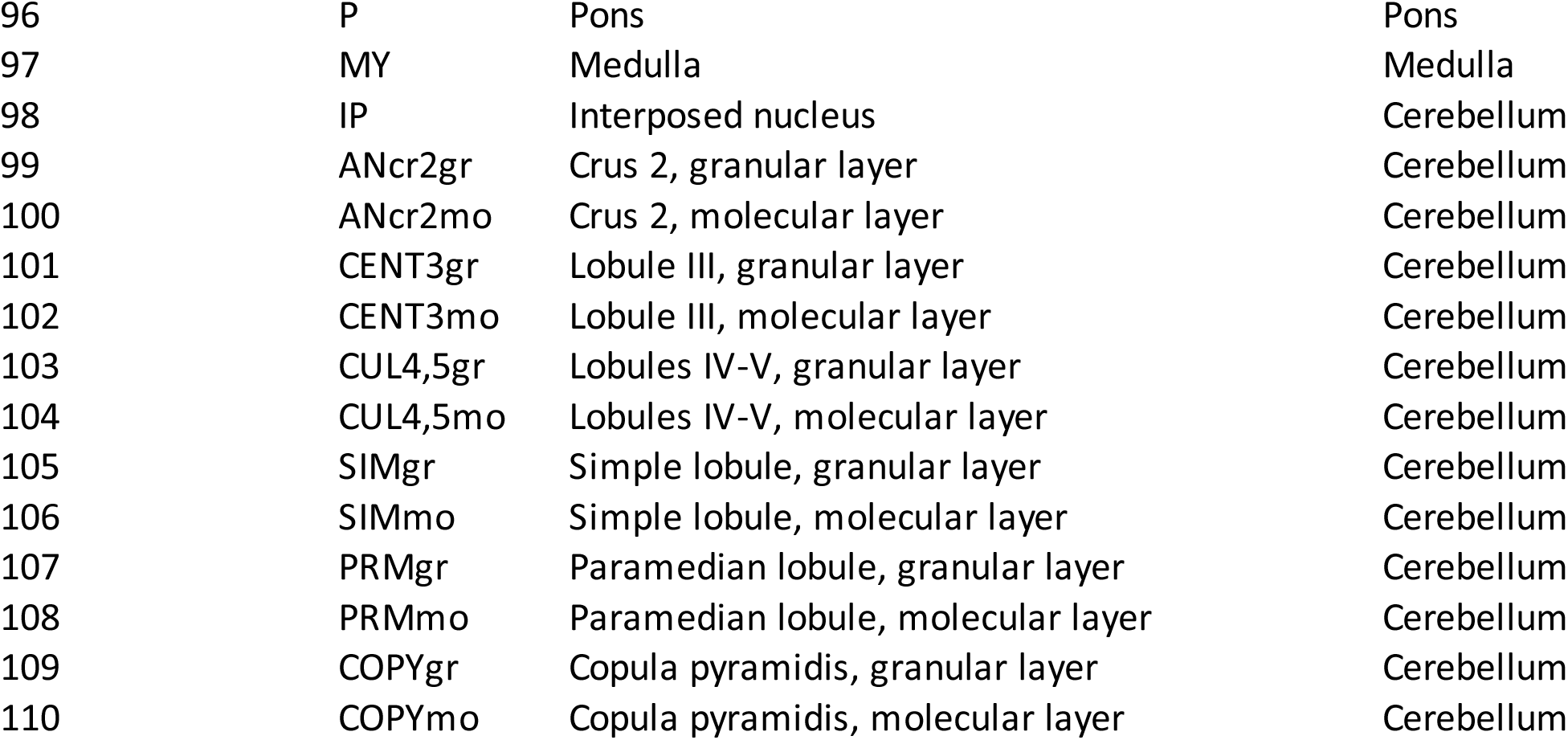
Brain regions and subregions. Column A, subregion number; column B, subregion abbreviation; column C, subregion name; column D, brain region.

## KEY RESOURCES TABLE

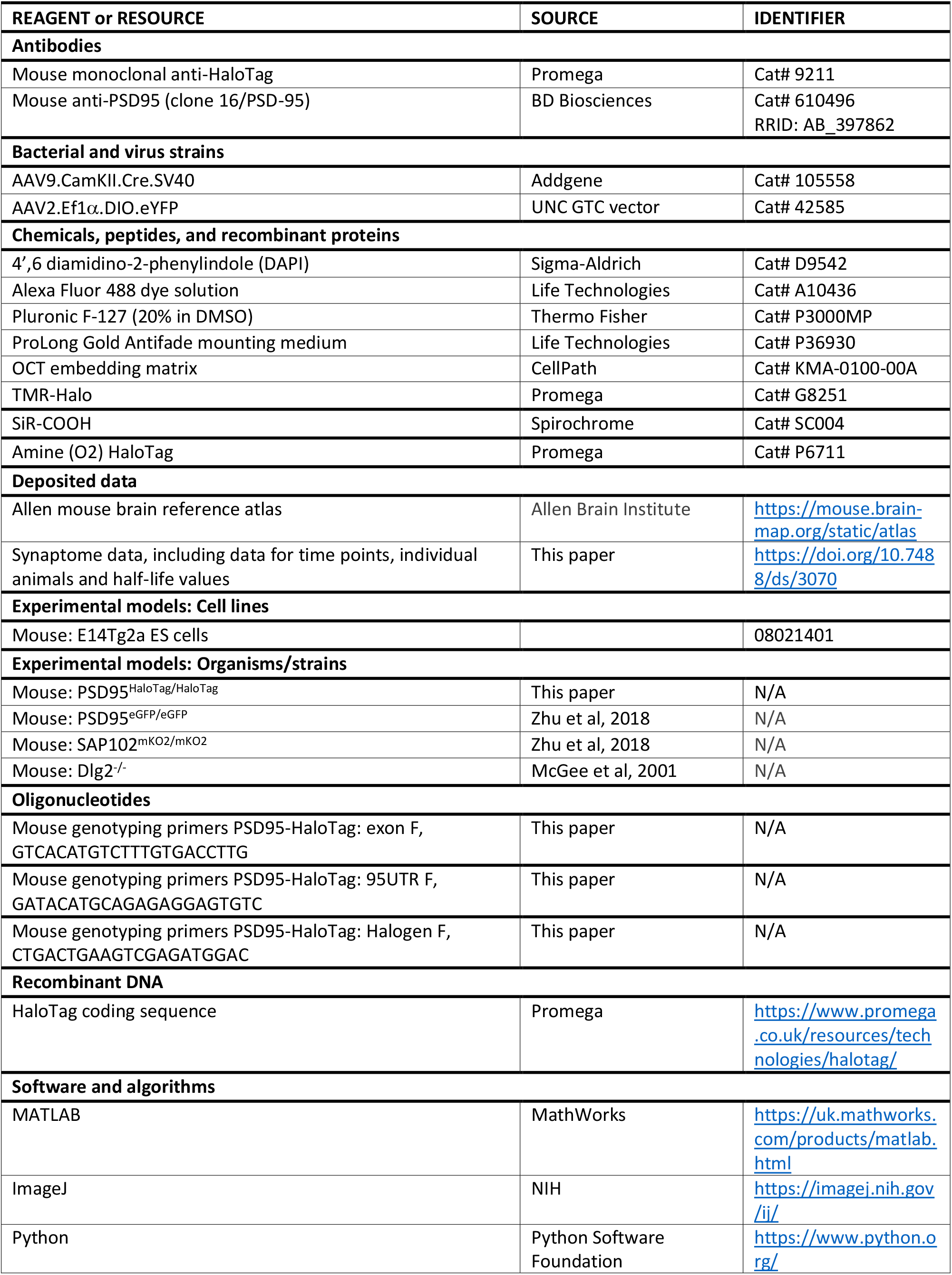
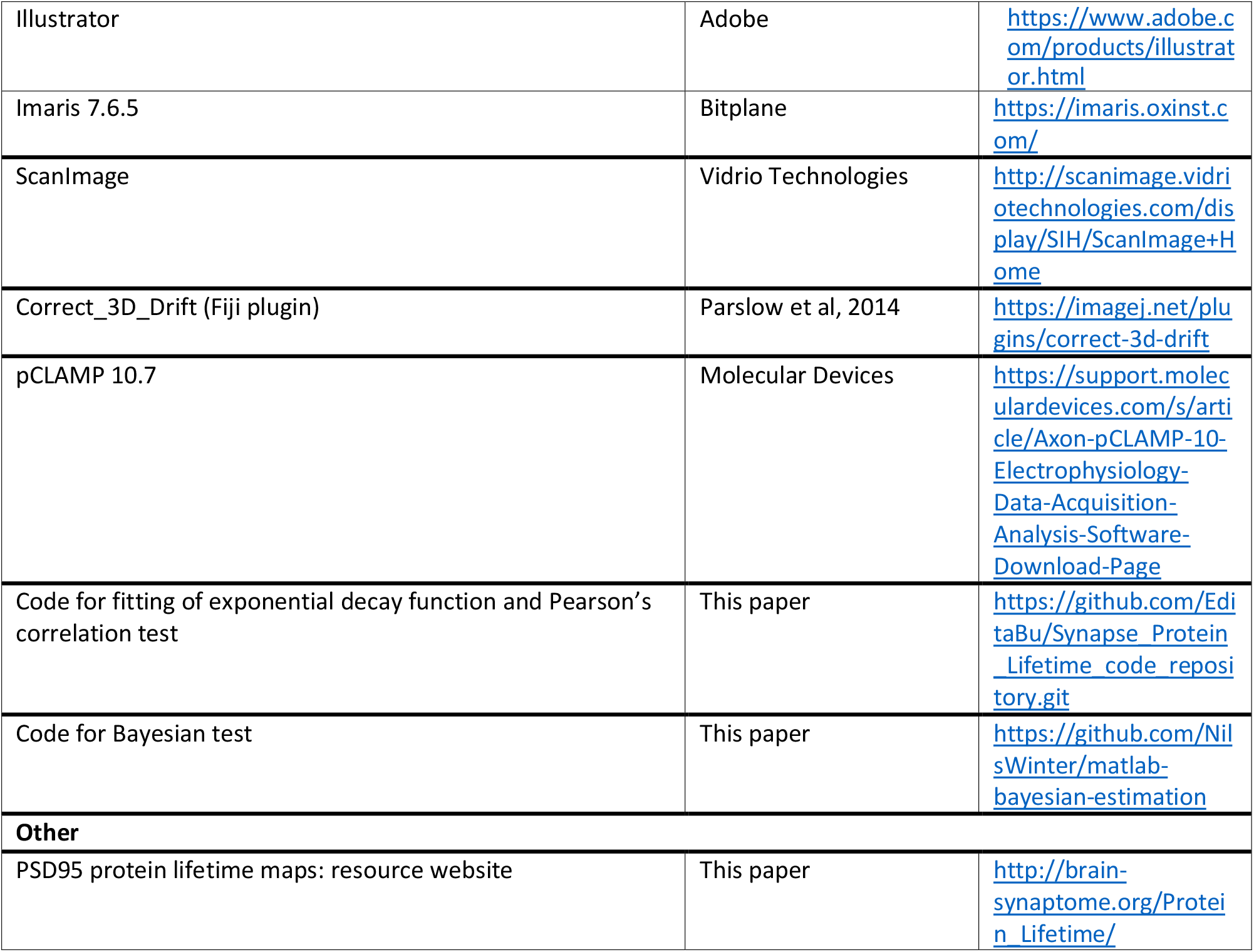

